# Evolution and Diversification of Carboxylesterase-like [4+2] Cyclases in Aspidosperma and Iboga Alkaloid Biosynthesis

**DOI:** 10.1101/2023.10.24.563752

**Authors:** Matthew D. DeMars, Sarah E. O’Connor

**Affiliations:** Department of Natural Product Biosynthesis, Max Planck Institute for Chemical Ecology, 07745 Jena, Germany

**Keywords:** monoterpene indole alkaloids, natural product biosynthesis, [4+2] cycloaddition, carboxylesterase, ancestral sequence reconstruction, enzyme evolution

## Abstract

Monoterpene indole alkaloids (MIAs) are a large and diverse class of plant natural products, and their biosynthetic construction has been a subject of intensive study for many years. The enzymatic basis for the production of aspidosperma and iboga alkaloids, which are produced exclusively by members of the Apocynaceae plant family, has recently been discovered. Three carboxylesterase (CXE)-like enzymes from *Catharanthus roseus* and *Tabernanthe iboga* catalyze regio- and enantiodivergent [4+2] cycloaddition reactions to generate the aspidosperma (tabersonine synthase, TS) and iboga (coronaridine synthase, CorS; catharanthine synthase, CS) scaffolds from a common biosynthetic intermediate. Here, we use a combined phylogenetic and biochemical approach to investigate the evolution and functional diversification of these cyclase enzymes. Through ancestral sequence reconstruction, we provide evidence for initial evolution of TS from an ancestral CXE followed by emergence of CorS in two separate lineages, leading in turn to CS exclusively in the *Catharanthus* genus. This progression from aspidosperma to iboga alkaloid biosynthesis is consistent with the chemotaxonomic distribution of these MIAs. We subsequently generate and test a panel of chimeras based on the ancestral cyclases to probe the molecular basis for differential cyclization activity. Finally, we show through partial heterologous reconstitution of tabersonine biosynthesis using non-pathway enzymes how aspidosperma alkaloids could have first appeared as “underground metabolites” via recruitment of promiscuous enzymes from common protein families. Our results provide insight into the evolution of biosynthetic enzymes and how new secondary metabolic pathways can emerge through small but important sequence changes following co-option of preexisting enzymatic functions.

## Introduction

Monoterpene indole alkaloids (MIAs) comprise a large class of secondary metabolites found primarily in select plants of the Gentianales order. With more than 3000 unique compounds reported to date (1), MIAs encompass a large diversity of chemical scaffolds that endow them with a broad range of biological activities (e.g., anti-cancer, anti-inflammatory, anti-malarial, anti-hypertensive) (2, 3). Despite their diverse structures, virtually all known MIAs derive from the common biosynthetic precursor strictosidine (4) (**Figure 1**). MIAs are organized into unique structural classes, with members of the corynanthe, strychnos, aspidosperma, and iboga groups representing a large proportion of the MIA scaffolds that have been isolated to date (2, 5). Notably, MIAs belonging to certain structural classes have only been isolated from plants in specific families. For instance, the aspidosperma and iboga MIAs are only found in plants belonging to the Apocynaceae family. Natural products bearing these scaffolds possess potent pharmaceutically relevant bioactivities such as anti-cancer (e.g., vinblastine) (6–8) and anti-addiction (e.g., ibogaine) (9–11) (**Figure 1**).

**Figure 1.**
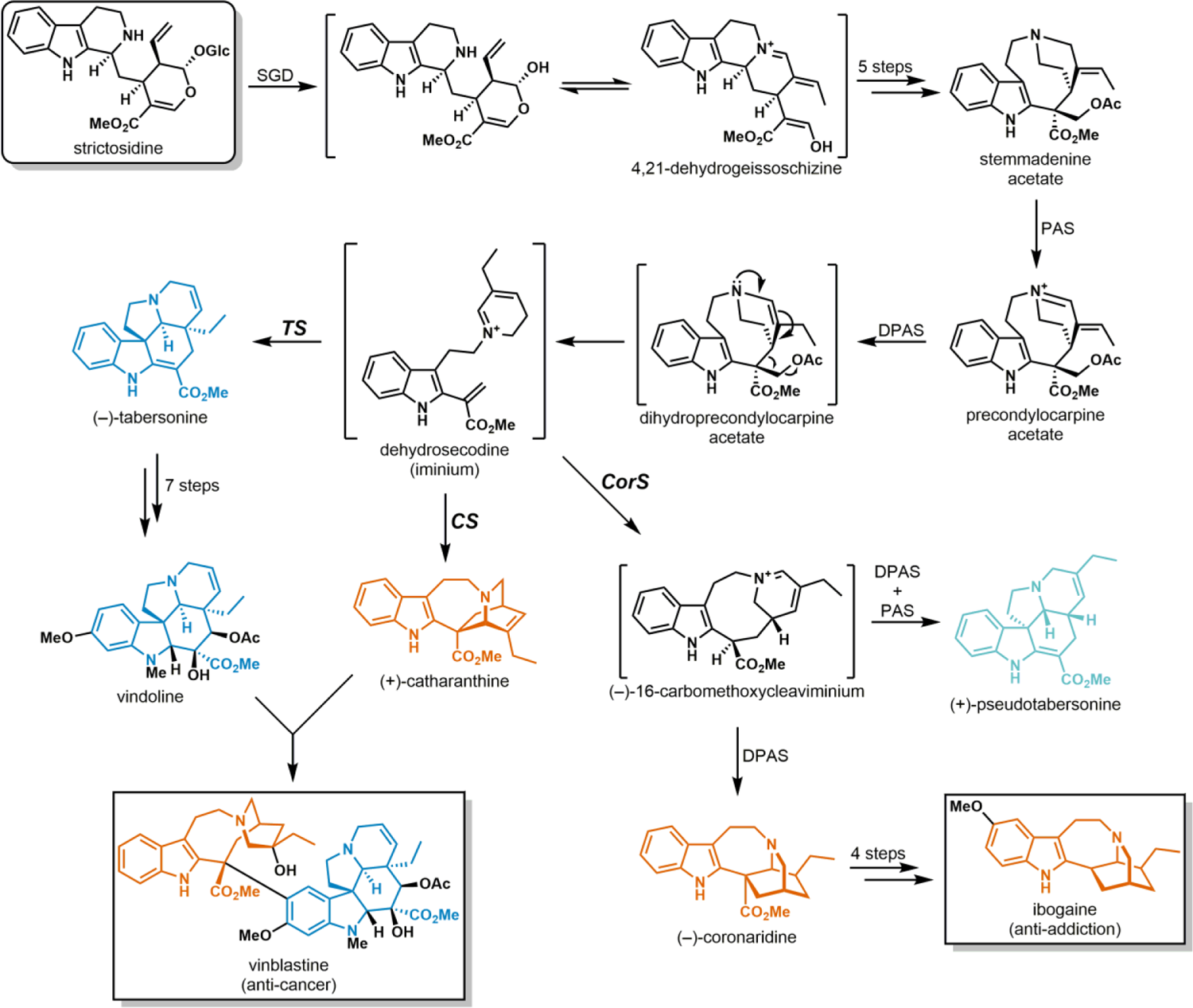
Abbreviated MIA biosynthetic pathways in *Catharanthus roseus* (vinblastine) and *Tabernanthe iboga* (ibogaine). Aspidosperma and iboga alkaloid scaffolds are shown in blue and red, respectively. The pseudoaspidosperma alkaloid (+)-pseudotabersonine is shown in light blue. Enzyme abbreviations: SGD (strictosidine β-D-glucosidase), PAS (precondylocarpine acetate synthase), DPAS (dihydroprecondylocarpine acetate synthase), TS (tabersonine synthase), CS (catharanthine synthase), CorS (coronaridine synthase).

Despite initial investigations into the biosynthesis of the aspidosperma and iboga alkaloids dating back six decades (12), only recently have the biosynthetic steps involved in their construction been unveiled in complete detail (13–16). In the vinblastine producer *Catharanthus roseus*, the pathway leading from strictosidine to the first aspidosperma or iboga scaffold requires nine individual enzymatic steps (Figure 1). In the key scaffold-forming step, one of two α/β hydrolase-type cyclases catalyzes a [4+2] cycloaddition reaction to generate either the aspidosperma alkaloid (–)-tabersonine (tabersonine synthase, TS) or the iboga alkaloid (+)-catharanthine (catharanthine synthase, CS) from an unstable intermediate (dehydrosecodine).

Recently, we identified and characterized an additional cyclase (coronaridine synthase, CorS) in the ibogaine producer *Tabernanthe iboga* responsible for formation of both (–)-coronaridine (iboga type) and (+)-pseudotabersonine (pseudoaspidosperma type) (16, 17). We demonstrated that this enzyme acts on the same substrate to generate an iminium-bearing intermediate en route toward these two structurally distinct compounds (Figure 1). We also recently solved the crystal structures of TS, CS, and CorS and performed extensive mutational analysis of these enzymes, which provided some key insights into their catalytic mechanism and cyclization specificity (18).

Here, we explore the evolutionary basis for MIA structural diversity by investigating the phylogenetic distribution of and relationships between these cyclases and their closest extant homologs among diverse MIA- and non-MIA-producing plants. We then use ancestral sequence reconstruction as a tool to understand the natural evolution and diversification of cyclase function in the Apocynaceae. Our results provide evidence for an evolutionary trajectory starting from an ancestral carboxylesterase that loses its original catalytic function before being co-opted for aspidosperma MIA biosynthesis. In two separate lineages, gene duplication and functional diversification lead from cyclases that produce aspidosperma alkaloids to those enabling the generation of pseudoaspidosperma and iboga alkaloids. Furthermore, we show through targeted mutagenesis studies how relatively minor amino acid changes in individual enzymes can lead to the emergence and enhancement of new chemistry, thus fundamentally altering and expanding upon preexisting biosynthetic pathways. We also gain broader insight into MIA biosynthetic pathway evolution by demonstrating how the highly reactive substrate for these cyclization reactions can be readily produced from a stable biosynthetic intermediate using extant enzymes from different pathways.

## Results

### Phylogenetic analysis of MIA cyclases and their closest homologs

On the basis of their primary amino acid sequences, the cyclases from *C. roseus* and *T. iboga* are classified as 2-<u>h</u>ydroxyisoflavanone <u>d</u>ehydratase (HID)-like proteins. In leguminous plants, HIDs are carboxylesterase-like enzymes that catalyze dehydration of 2-hydroxyisoflavanones as one of the steps in isoflavone biosynthesis (19). Along with the acylsugar acylhydrolases (ASHs) from *Solanum* spp. (20), the tuliposide-converting enzymes (TCEs) from *Tulipa gesneriana* (21, 22), the gibberellin receptors (GID1s) (23, 24), and a number of other characterized and as yet uncharacterized plant proteins (**Figure S1**), HIDs can be broadly classified as class I carboxylesterases (CXEs) (25, 26). Members of this class all adopt the α/β-hydrolase fold and typically share some key sequence signatures, including the His-Gly-Gly-Gly motif comprising the oxyanion hole and the Ser-His-Asp catalytic triad. While most class I CXEs are thought to catalyze ester hydrolysis, there is a growing list of enzymes in this group that perform other types of chemistry (e.g., HIDs and TCEs) or that have lost catalytic function entirely (e.g., GID1s) (**Figure S1**).

We began our study by collating a group of putative HID-like cyclases and closely related sequences from online and in-house genome and transcriptome databases using TS from *C. roseus* (CrHID3) as a search query (**Table S1**). We then constructed a phylogenetic tree based on an alignment of 258 sequences, which included class I CXEs from *Arabidopsis thaliana* (27), HIDs from the Fabaceae (legume) family (19), ASHs from *Solanum* spp. (20), TCEs from *T. gesneriana* (21, 22), and a representative selection of GID1s (23, 24) among other previously described proteins with known and unknown functions (Figures 2a and **S2**). The known cyclases from *C. roseus* and *T. iboga* are found in a well-supported clade (Clade 1) consisting of 28 sequences from a total of 10 species in the Apocynaceae family (Figure 2b). Because they have relinquished their putative ancestral function, non-ester-hydrolyzing CXEs sometimes exhibit differences from canonical CXEs in certain key regions of their amino acid sequence. We have previously noted that the catalytic triad of the cyclases is disrupted, with the aromatic residues Tyr or Phe substituting for His (18). In addition, the characteristic His-Gly-Gly-Gly of the oxyanion hole is replaced with His-Gly-Ala-Gly in the cyclases. Among the sequences in our phylogenetic analysis, 16 share both of these features (although Gly also substitutes for the catalytic Ser in MtHID2), and each of the 9 species from which they derive is known to produce aspidosperma and, in some cases, iboga alkaloids. Notably, while MIAs are produced by a number of other species that are represented in our phylogeny, production of the aspidosperma and iboga types is limited to just these nine species (**Table S2**). Along with three additional sequences (TdHID3, TeHID3, TiHID3) that also lack a functional catalytic triad but that retain the His-Gly-Gly-Gly oxyanion hole, the 16 cyclase-like sequences comprise a strongly supported subclade of the one described above (Clades 1a and 1b in Figure 2). The nine sequences remaining in Clade 1 (Clade 1c in Figure 2) bear all of the canonical sequence signatures of catalytically active CXEs.

**Figure 2.**
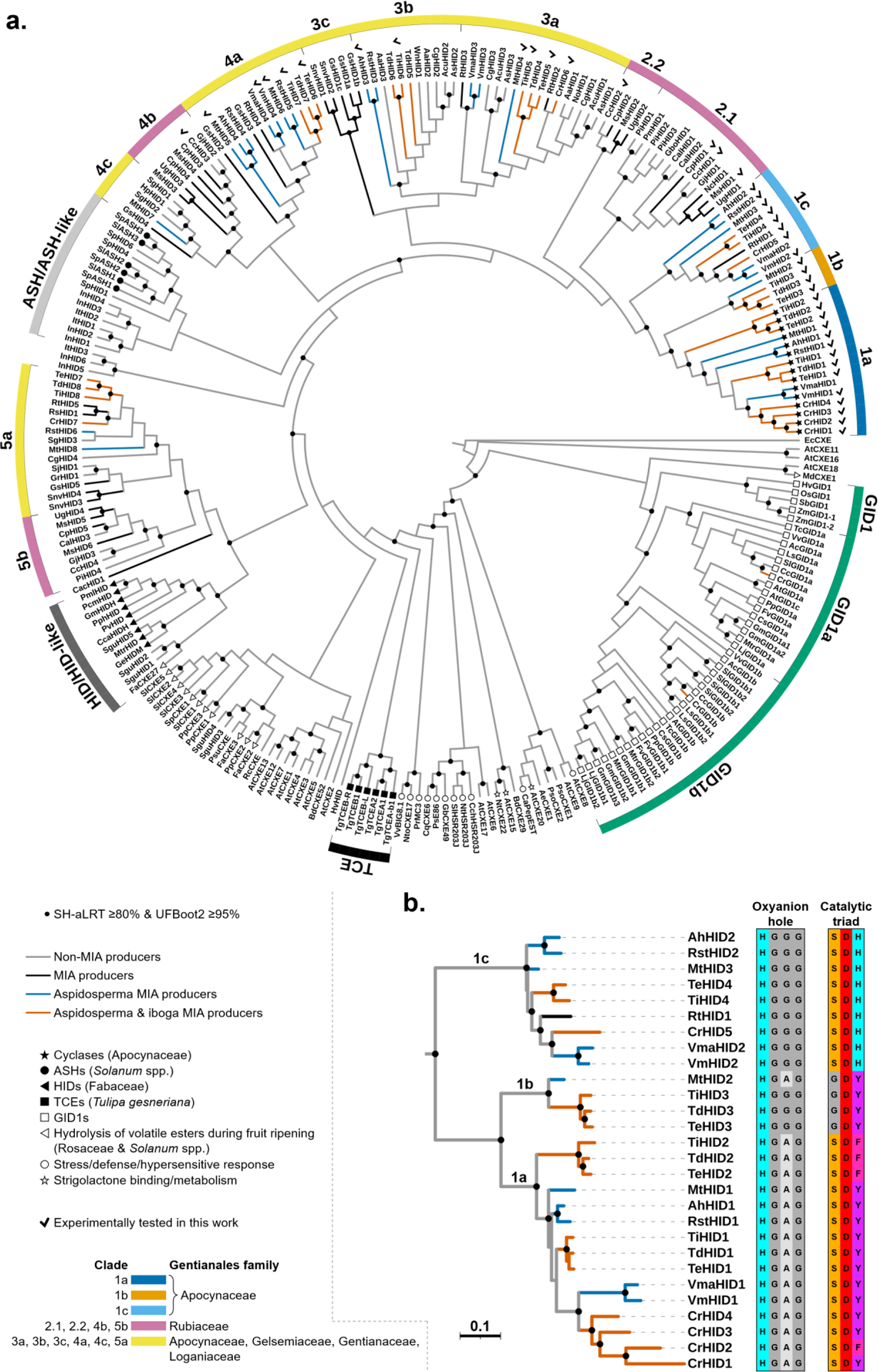
Phylogenetic analysis of plant class I carboxylesterases (CXEs). (**a**) Maximum likelihood tree (cladogram), including major characterized CXE subclasses and their reported in vivo functions (shown as symbols at the tips). Black circles at the nodes indicate strong support for the respective clade (SH-aLRT support ≥80% and ultrafast bootstrap (UFBoot2) support ≥95%). Clades 1-5 are divided into subclades on the basis of enzymatic function and phylogenetic distribution within the Gentianales order. Clades are labeled and colored as in Figure 4. Checkmarks appear next to all enzymes experimentally tested in the present work. See **Figure S2** for a full phylogram with SH-aLRT/UFBoot2 support values and branch lengths. Abbreviations: ASH (acylsugar acylhydrolase), HID (2-hydroxyisoflavanone dehydratase), TCE (tuliposide-converting enzyme), GID1 (gibberellin receptor). (**b**) Enlarged view of Clade 1 with oxyanion hole and catalytic triad residues shown next to each sequence name. Scale bar length represents 0.1 amino acid substitutions per site.

Two somewhat weakly supported clades composed of 17 closely related HID-like CXEs from Rubiaceae spp. appear close to the putative cyclases (Clades 2.1 and 2.2 in Figure 2a). Additional clades branching off earlier in the tree (Clades 3–5) consist of CXEs from species in the five families that together comprise the Gentianales order (Apocynaceae, Gelsemiaceae, Gentianaceae, Loganiaceae, and Rubiaceae) (28, 29). While the vast majority of these CXEs possess the typical oxyanion hole and catalytic triad residues (**Table S1**), their in vivo or in vitro catalytic function has never been investigated. The most closely related enzymes that have been functionally characterized are the *Solanum* ASHs (20) and the HIDs from *Glycine max*, *Glycyrrhiza echinata*, and *Pueraria montana* var. *lobata* (19, 30). In the latter, Thr substitutes for the catalytic Ser (**Table S1**) and is required for optimal activity. The sequences of the *T. gesneriana* TCEs and the *Arabidopsis* CXEs (AtCXEs) are more divergent and are thus located further away from the cyclases and their closest relatives (Figure 2a). Finally, the GID1s appear closest to the root of the tree and are thought to have evolved from an ancestral catalytic CXE (23, 24). These proteins retain most of the canonical residues with the exception of the catalytic His, which is replaced with either Ile or Val (**Table S1**).

### In vitro functional analysis of extant HID-like cyclases and CXEs

With a suitable phylogeny of the HID-like cyclases and CXEs in hand, we next verified the catalytic function of the different enzymes represented in each major clade of the tree (Figure 2a). We selected a representative set of enzymes to express, purify, and test in small-scale in vitro reactions with angryline (an acid-stable constitutional isomer of the dehydrosecodine substrate) (18) along with 4-nitrophenyl butyrate (4-NPB) and 4-methylumbelliferyl butyrate (4-MUB) to serve as model substrates for any active esterases.

Given the strong correlation between the presence of proteins with key cyclase-like sequence signatures and the chemotaxonomic distribution of the aspidosperma and iboga classes of MIAs, we initially hypothesized that Clades 1a and 1b would contain all of the HID-like cyclases present in the nine representative aspidosperma/iboga alkaloid producers included in our phylogenetic analysis. With the exception of CrHID4, which we deemed a pseudogene (**Figure S3**), all of the enzymes tested from Clade 1a exhibited high angryline cyclization activity (Figure 3 and **Table S3**). CrHID1 (previously characterized CrCS (13, 15, 18)) formed catharanthine whereas TiHID2 (previously characterized TiCorS (16–18)) along with CrHID2, TdHID2, and TeHID2 all formed (–)-16-carbomethoxycleaviminium (16-cmc) as the major product. All of the other Clade 1a enzymes showed high activity and cyclized angryline to selectively form tabersonine. Consistent with previous results (18), none of these cyclases could catalyze hydrolysis of the model esterase substrates (**Table S4**). Moreover, despite their high similarity to the cyclases in Clade 1a (average = 69% ID), we did not observe turnover of angryline or the esterase substrates when we tested the four proteins in Clade 1b (Figure 3 and **Table S4**). One key feature that distinguishes the sequences in Clade 1b from those in Clade 1a is the lack of *both* a catalytic Ser and a catalytic His in the active site (Figure 2b). Restoring the catalytic Ser negatively impacted the soluble expression levels of MtHID2 and TiHID3, and no gain of catalytic function was observed in either case (**Figure S4** and **Table S4**).

**Figure 3.**
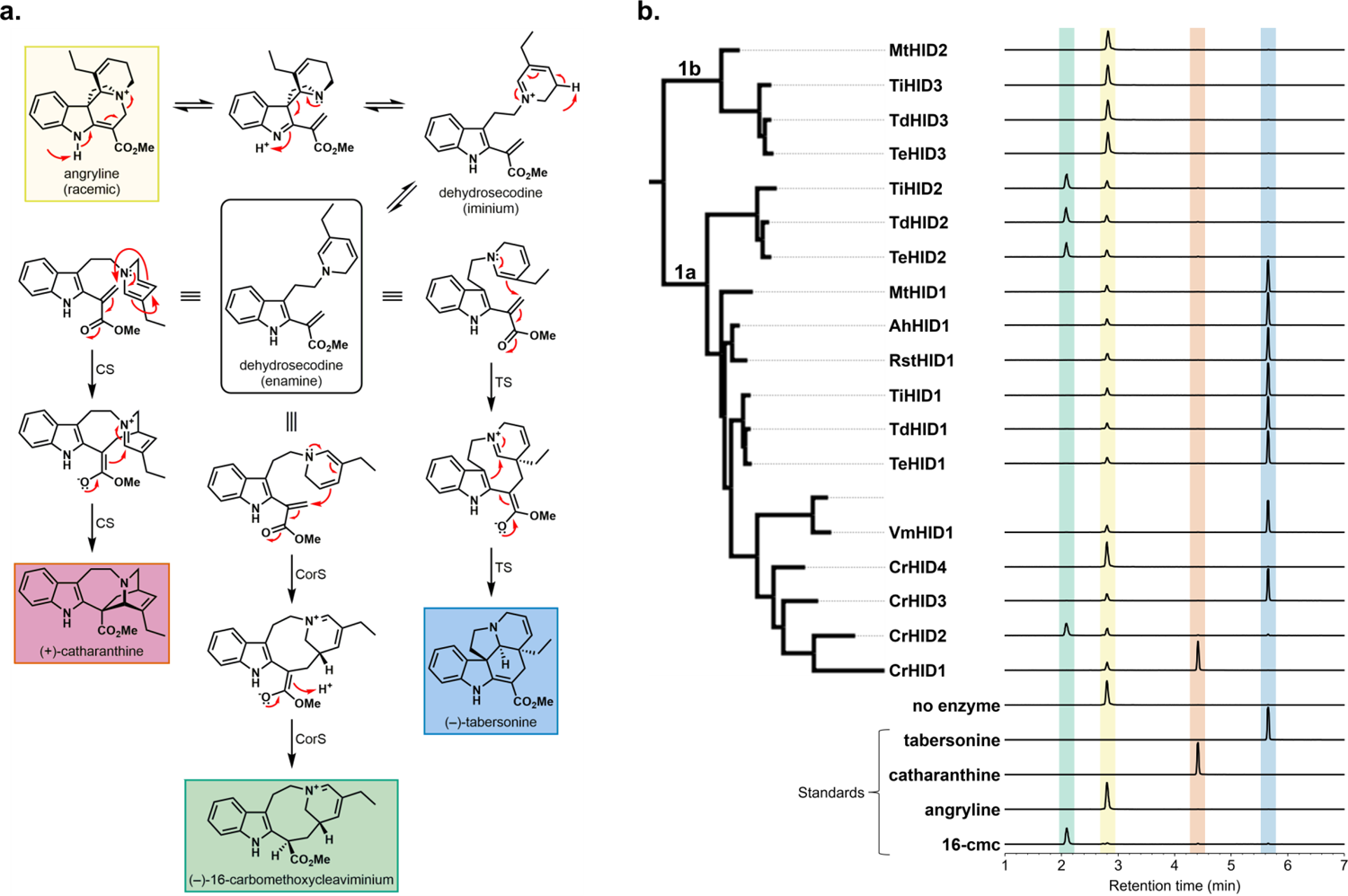
In vitro functional analysis of HID-like cyclases. (**a**) Cyclization reactions catalyzed by CS, CorS, and TS. Angryline, (+)-catharanthine, and (–)-tabersonine are the only stable and isolable compounds depicted in this scheme. (–)-16-carbomethoxycleaviminium (16-cmc) is stable enough to be observed as a distinct peak by LC-MS, but it is not isolable. (**b**) LC-MS traces (EIC 337.1911 ± 0.01) of reactions between angryline and enzymes from Clades 1a and 1b of the phylogeny. Reactions were performed at 37 °C for 30 min in 50 mM Tris (pH 9.0) with 50 µM angryline and either 1 µM (Clade 1a) or 5 µM (Clade 1b) enzyme. The 16-cmc standard is the reaction with TiHID2 (10 µM) performed at 37 °C for 1 h.

In contrast to the proteins in Clades 1a and 1b, we anticipated that those in Clade 1c would exhibit at least some level of esterase activity because the residues of both the oxyanion hole and the catalytic triad are identical to those present in most class I CXEs (Figure 2b). However, these enzymes exhibited virtually no hydrolytic activity toward 4-NPB and 4-MUB (**Table S4**). AlphaFold models of these CXE-like proteins revealed that the catalytic Asp may be positioned outside of the active site, offering one plausible explanation for their poor activity (**Figure S5**). However, we cannot rule out the possibility that the physiological substrates for these enzymes differ considerably from the model substrates we employed in this study. As expected, the Clade 1c proteins were also unable to catalyze cyclization of angryline (**Figure S6**). From all of the remaining clades, 16 representative enzymes showed low to very high rates of hydrolysis when presented with the model substrates (**Table S4**). While angryline did not cyclize in the presence of these esterases, we did observe moderate levels of substrate depletion in some cases (**Figure S7**).

### Reconstruction and functional analysis of ancestral HID-like enzymes

In order to gain further insight into the evolutionary history of the cyclases, we sought to reconstruct ancestral sequences corresponding to several key nodes in a more focused phylogeny consisting only of sequences from Clades 1–5 as well as those from the ASH/ASH-like and HID/HID-like clades (Figures 4a and **S8**). We inferred a total of nine ancestral nodes by maximum likelihood (labeled AncHID1–AncHID9 in Figure 4a), and all of the corresponding proteins were expressed at high-levels in *E. coli*, purified, and tested in vitro with angryline and the model esterase substrates. The ancestor of all extant cyclases (AncHID4) displayed TS activity, cyclizing angryline to generate tabersonine as the major product (Figure 4b) along with low levels of 16-cmc when tested at a higher concentration (**Table S3**). Ancestral TS activity is consistent with our constructed phylogeny as well as the reported chemotaxonomic distribution of the aspidosperma alkaloids, which are found in many more Apocynaceae species than the pseudoaspidosperma and iboga alkaloids. Almost all of the residues within or close to the predicted substrate binding pocket of AncHID4 are identical to those found in all of the experimentally verified extant TS enzymes (**Figure S9**). One key exception is the presence of a Gly instead of a Trp at position 171 of the ancestral sequence; all extant TSs with the exception of VmHID1 have a Trp at this position whereas all extant CorSs and the vast majority of closely related extant CXEs have a Gly (**Figure S9**). Mutating this Gly in AncHID4 to Trp (G171W) led to a marked increase in cyclase activity to produce tabersonine that was concomitant with elimination of 16-cmc as a minor side product (Figure 4b and **Table S3**). Although this residue is located outside of the substrate binding pocket (**Figure S10**), the positive impact of the GèW substitution on substrate turnover and product specificity must have been significant enough to ensure its retention throughout the course of further TS evolution.

**Figure 4.**
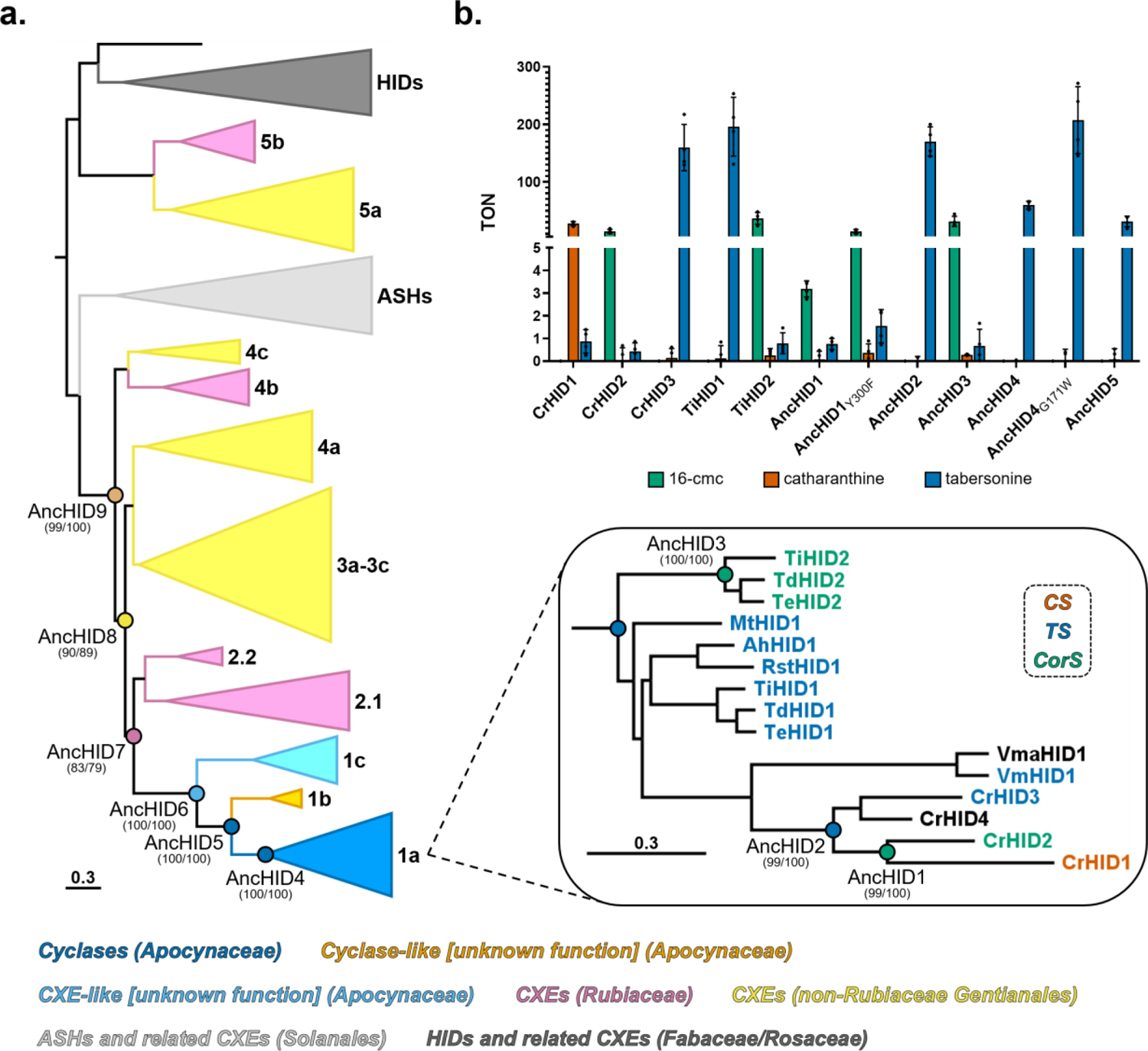
Reconstruction and in vitro functional analysis of ancestral HID-like enzymes. (**a**) Phylogeny with nodes targeted for ancestral sequence reconstruction highlighted. Clades are labeled and colored according to enzymatic function/taxonomic distribution as in Figure 2. Values at each of the reconstructed nodes represent SH-aLRT support (%) / ultrafast bootstrap support (%). Scale bar length represents 0.3 nucleotide (nt) substitutions per codon site (equivalent to 0.1 nt substitutions per nt). Clade 1a containing all of the extant cyclases is enlarged at the bottom right, and the names of individual enzymes are colored according to experimentally verified function. CrHID4 exhibits minimal TS activity, and VmaHID1 was not tested but is predicted to be a TS. See **Figure S8** for a full phylogram with SH-aLRT/UFBoot2 support values and branch lengths. (**b**) Angryline cyclization activity of selected extant and ancestral cyclases expressed as turnover number (TON = mol product/mol enzyme). Colored bar height indicates the mean of four independent experiments. The results of individual experiments are represented by dots (error bars = SD). All reactions were performed at 37 °C for 30 min in 50 mM Tris (pH 9.0) with 50 µM angryline and 40 nM enzyme.

Given that *C. roseus* itself possesses each of the three unique cyclases, we sought to determine the function of the ancestor from this organism (AncHID2). In reactions with angryline, this enzyme produced tabersonine in a highly selective manner; no other products were detected even in trace amounts (Figure 4b and **Table S3**). This result was unsurprising considering the large number of shared substrate binding pocket and surrounding residues between AncHID2, AncHID4, and the extant TSs (**Figure S9**). While the CrHID1/CrHID2 (CS/CorS) ancestor AncHID1 is most closely related to AncHID2 (TS) at the sequence level (93% ID, 21 amino acid differences), it bears the highest similarity to CrHID2 (CorS) specifically among the amino acid residues located in and around the substrate binding pocket (**Figure S11**). One key difference is the presence of a **Tyr**-Phe-Glu motif in the ancestral sequence (where **Tyr** occupies the position of the catalytic His in the esterases), which is also found in CrHID1 (CS) and almost all of the extant and ancestral TSs (**Figure S9**). In contrast, in all extant CorSs, this motif is replaced with **Phe**-Phe-Asp. When assayed with angryline, AncHID1 formed 16-cmc as the major product along with very low levels of tabersonine, but its overall activity was significantly lower than that of CrHID2 and the other extant CorS enzymes (Figure 4b and **Table S3**). Replacing the Tyr in the **Tyr**-Phe-Glu motif with a Phe (Y300F) enhanced the overall activity of this ancestral cyclase (Figure 4b and **Table S3**). As predicted given the presence of the **Phe**-Phe-Asp motif as well as its high sequence identity to the extant CorS enzymes from the Tabernaemontaneae tribe (97% ID, 9–11 amino acid differences) (**Figure S11**), AncHID3 produced 16-cmc at relatively high levels as the major product in reactions with angryline (Figure 4b and **Table S3**).

Although we were unable to establish any in vitro functions for the proteins in Clade 1b of the phylogeny (Figure 3b), the common ancestor of these proteins and the cyclases (AncHID5) showed TS activity (Figure 4b) and also produced low amounts of 16-cmc when tested at a higher concentration (**Table S3**). Most of the 27 amino acid differences between AncHID5 and AncHID4 occur outside of the predicted substrate binding pocket (**Figure S12**). One noteworthy exception is the oxyanion hole motif, which appears as His-Gly-Gly-Gly in AncHID5 and most active CXEs but is curiously replaced with His-Gly-Ala-Gly in all of the extant and ancestral cyclases (**Figure S9**). Mutating the oxyanion hole Gly in AncHID5 to Ala (G84A) had no effect on TS activity but led to a minor increase in 16-cmc as a side product (**Figure S12** and **Table S3**). The reciprocal mutation in AncHID4 (A84G) resulted in increased TS activity, but the low level of 16-cmc present in reactions with the wild-type ancestor was further reduced (**Figure S12** and **Table S3**). Thus, while cyclase function is not strictly dependent on modifications to the canonical oxyanion hole, the change from His-Gly-Gly-Gly to His-Gly-Ala-Gly appears to have helped marginally broaden the product profile of the cyclases.

The proteins in Clade 1c of the phylogeny are the closest extant relatives of the cyclases that possess the canonical oxyanion hole and catalytic triad residues (Figure 2b) but that nonetheless exhibit extremely low esterase activity (**Table S4**). Testing the common ancestor of these proteins and the cyclases (AncHID6) revealed that it too was incapable of catalyzing significant levels of hydrolysis (**Table S4**). However, unlike its descendants in Clade 1c, this ancestor did display very low (TON <1) but reproducible TS activity (**Figure S6** and **Table S3**). To identify an ancestor with high esterase and no cyclase activity, we chose to reconstruct several early ancestral CXE-like sequences (AncHID7, AncHID8, and AncHID9) due to suboptimal support values at the node corresponding to AncHID7, which could represent the most recent common ancestor of the cyclases and the most active extant CXEs (Figure 4a). These three ancestors are highly similar to one another (≥94% ID) (**Figure S11**), with all non-conservative differences confined to regions outside of the predicted substrate binding pocket (**Figure S13**). Each of these proteins showed exceptionally high rates of hydrolysis when acting on the model substrates (**Table S4**) and no cyclase activity toward angryline (**Figure S7**). On the basis of these results, we surmise that basal cyclase (TS) activity must have arisen at some point between AncHID7 and AncHID6, with significant enhancement of this low-level activity occurring between AncHID6 and AncHID5. Notably, as AncHID5 is the first ancestor for which the catalytic His has been replaced (**Figure S9**), the observed boost in TS activity appears to have been concomitant with substitution of this important residue.

### Exploring the molecular basis for cyclase evolution and diversification

We next sought to uncover the molecular-level details underlying the evolution of a cyclase (TS) from an ancestral CXE as well as the subsequent change in cyclization regioselectivity. First, we used our previously solved crystal structures and docking models of the cyclases (18) to locate all residues within 6 Å of bound substrate. Multiple sequence alignment of the extant and ancestral proteins then allowed us to identify seven contiguous regions that we would target for mutagenesis in each of the ancestors (Figure 5 and **Figure S9**). We began by working backward along the evolutionary trajectory, first investigating the evolution of the only known CS (CrHID1) from a CorS precursor (AncHID1) in *C. roseus*. Of the 52 total amino acid differences between these two proteins, less than half (23) are located in the seven regions in and around the substrate binding pocket (**Figure S14**). Replacing 19 of these residues in AncHID1 with those found in CrHID1 (including the MISTTP extended loop just before the catalytic Asp) resulted in a cyclase (AncHID1_CrHID1_-M1) that generated catharanthine as the primary product, retaining very little residual CorS activity (Figure 5c and **Table S3**). Reverting two amino acids back to those of the wild-type ancestor (AncHID1_CrHID1_-M2) had virtually no impact on the product profile (Figure 5c and **Table S3**). We further dissected the regions containing these 17 changes sufficient for switching from CorS to CS activity: the loop between alpha helices 4 and 5 (α4/α5 loop; 4 changes), alpha helix 5 (α5; 5 changes), and the extended loop/catalytic Asp (8 changes). Although the four differing residues in the α4/α5 loop are located outside of the substrate binding pocket (**Figure S15**), the mutant lacking these changes (AncHID1_CrHID1_-M3) exhibited a significant drop in CS activity concomitant with increased production of 16-cmc and tabersonine (Figure 5c and **Table S3**). Testing other possible combinations of changes revealed that residues in all three regions play an important role in influencing the product specificity of CS (**Figure S15** and **Table S3**).

**Figure 5.**
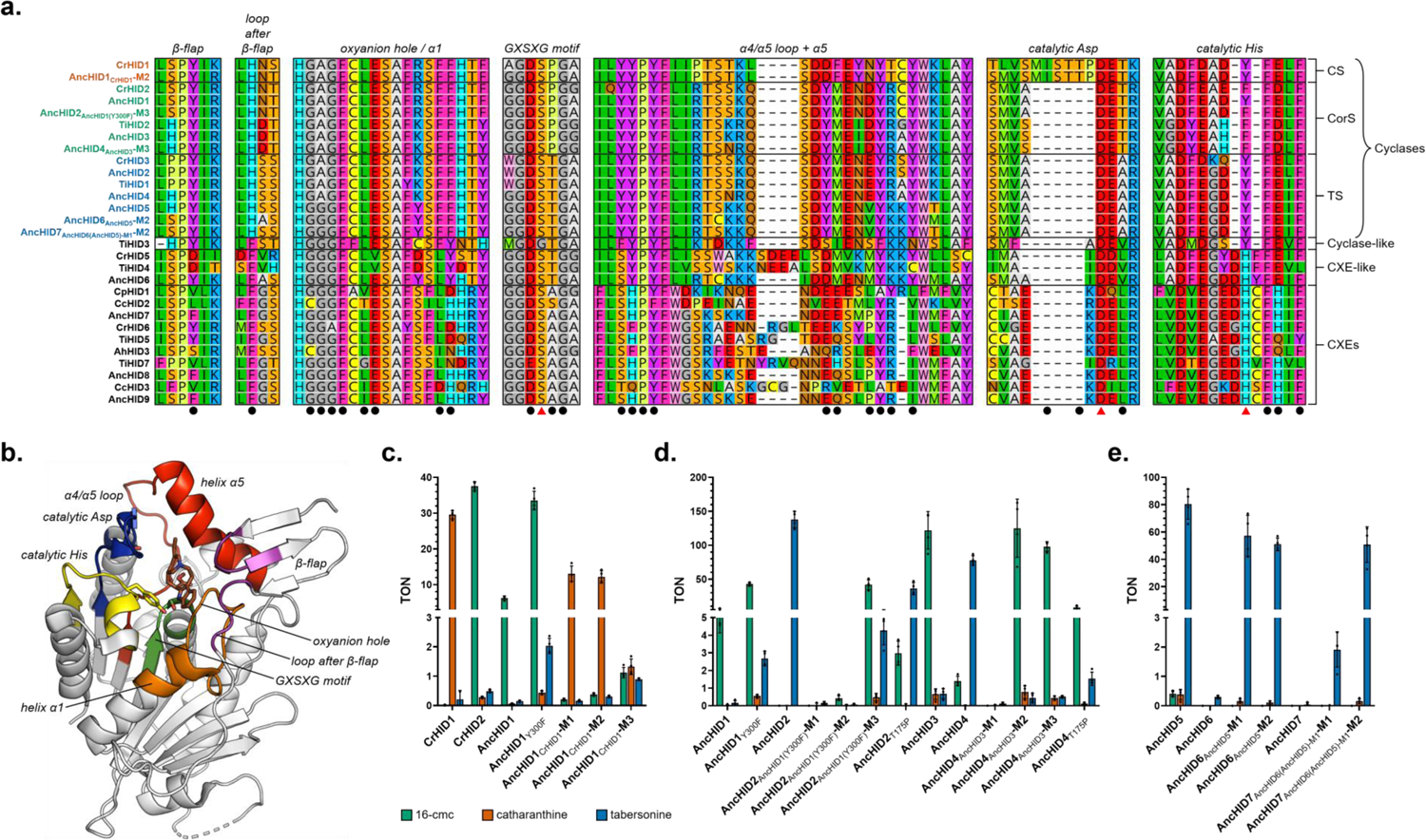
Multiple sequence alignment (MSA) of extant and ancestral enzymes and cyclization activity of key ancestral HID mutants. (**a**) MSA of selected extant and ancestral HID-like enzymes. Seven contiguous regions encompass all amino acid residues located within 6 Å of the substrate binding pocket (black circles). Red triangles are located below the three residues making up the catalytic triad. The names of the individual cyclases are colored according to the identity of their major cyclization product (red = catharanthine, green = 16-cmc, blue = tabersonine). CXEs with experimentally verified esterase activity from each of the major subclades (2.1, 2.2, 3a, 3b, 4a, 4b) are included in the alignment. See **Figure S9** for an expanded version with additional sequences. (**b**) Crystal structure of CrHID1 (**PDB 6RT8**) with each of the seven key regions surrounding the substrate binding pocket highlighted. Bound intermediate (16-carbomethoxycleaviminium, brown) and the side chains of the catalytic triad residues (Ser = green, Tyr = yellow, Asp = blue) are shown as sticks. (**c**) Evolution of CS (CrHID1) from CorS (AncHID1) in *C. roseus*. Angryline cyclization activity of wild-type and selected mutant cyclases is expressed as turnover number (TON = mol product/mol enzyme). Colored bar height indicates the mean of four independent experiments. The results of individual experiments are represented by dots (error bars = SD). Reactions were performed at 37 °C for 30 min in 50 mM Tris (pH 9.0) with 50 µM angryline and 1 µM enzyme. (**d**) Evolution of CorS (AncHID1, AncHID3) from TS (AncHID2, AncHID4). Reactions were performed at 37 °C for 30 min in 50 mM Tris (pH 9.0) with 50 µM angryline and 200 nM enzyme. (**e**) Evolution of TS (AncHID5, AncHID6) from an ancestral CXE (AncHID7). Reactions were performed as described in **d**.

Our phylogenetic analysis and ancestral sequence reconstruction results demonstrated that CorS activity evolved independently at least twice: once in the Tabernaemontaneae and again in *Catharanthus*. Evidently, the path from AncHID2 (TS) to AncHID1 (CorS) in *C. roseus* was relatively short, consisting of only 21 amino acid changes. As wild-type AncHID1 could generate only low levels of 16-cmc, we targeted the corresponding Y300F mutant, which exhibited much higher turnover numbers (Figure 5d and **Table S3**). Of the 22 differences between AncHID2 and AncHID1_Y300F_, 12 are located in the 7 previously identified regions, and only 4 of the corresponding residues are expected to form part of the substrate binding pocket (**Figure S16**). Mutating these four amino acids as well as three additional residues found within the same four regions of AncHID2 (AncHID2_AncHID1(Y300F)_-M1) almost completely abolished cyclase activity toward angryline (Figure 5d and **Table S3**). Introducing the remaining five mutations and thus fully exchanging all seven regions between AncHID2 and AncHID1_Y300F_ resulted in an enzyme (AncHID2_AncHID1(Y300F)_-M2) exhibiting only a minor increase in CorS activity (Figure 5d and **Table S3**). Consequently, we elected to replace an additional four residues, leading to an AncHID2 mutant (AncHID2_AncHID1(Y300F)_-M3) with a total of 16 amino acid changes and differing from AncHID1_Y300F_ in only 6 positions located far from the substrate binding pocket (**Figures S16** and **S17**). This mutant displayed the same CorS activity as AncHID1_Y300F_ (Figure 5d and **Table S3**), indicating that amino acid residues outside of the substrate binding pocket can have a substantial impact on cyclase activity. Furthermore, systematic reversion of each of these residues showed that no single site was fully responsible for the observed change in product specificity (**Figure S17** and **Table S3**). However, it is interesting to note that replacing the active site Thr (present in all TSs) with Pro (present in all CorSs and CS) led to a single-site mutant (AncHID2_T175P_) capable of producing 16-cmc at low levels (Figure 5d and **Table S3**).

We similarly probed the evolution of CorS activity in the Tabernaemontaneae by generating mutants of AncHID4 (TS) targeting AncHID3 (CorS). Although AncHID4 can already produce low levels of 16-cmc, AncHID3 exhibits much higher CorS activity than AncHID1 and even AncHID1_Y300F_ (Figure 5d and **Table S3**). A total of 34 amino acid differences separate these two ancestral cyclases, 16 of which are located in the 7 key regions in and around the substrate binding pocket (**Figure S18**). Replacing all 16 of these residues in AncHID4 with those found in AncHID3 yielded a protein (AncHID4_AncHID3_-M1) with poor soluble expression levels and virtually no cyclase activity (Figure 5d and **Table S3**). Accordingly, we exchanged all but eight residues that we deemed too far removed from the substrate binding pocket to likely have any impact on enzyme activity, leading to a mutant (AncHID4_AncHID3_-M2) whose product profile was identical to that of AncHID3 (Figure 5d and **Table S3**). Reverting two amino acids back to those of the wild-type ancestor (AncHID4_AncHID3_-M3) led only to a modest reduction in overall activity (Figure 5d and **Table S3**). Several of the residues that were changed to achieve the switch in cyclase activity did not appear to have a significant effect when probed individually (**Figure S19** and **Table S3**). However, reversion of six mutations in alpha helix 8 (α8) at the C-terminus of the protein resulted in very low expression yields and complete abolition of activity. Similar to its effect on AncHID2, the T175P mutation in AncHID4 led to increased levels of 16-cmc at the expense of tabersonine (Figure 5d and **Table S3**). AncHID4_T175P_ generated 16-cmc as the major product, indicating that cyclase product specificity could be reversed via a single mutation (**Figure S20**).

We previously noted that AncHID6, while resembling a catalytically active CXE at the sequence level, shows negligible hydrolytic activity and can produce only very low amounts of tabersonine in reactions with angryline. According to our phylogeny and the reconstructed ancestral sequences, AncHID6 accumulated 49 changes in its amino acid sequence along the path toward AncHID5, a fully active TS (**Figure S21**). Only 19 of these changes are found in the 7 regions previously identified, indicating that portions of the protein further away from the substrate binding pocket were modified the most during the evolution of the first highly active cyclase (**Figure S21**). We found that substituting only 18 (AncHID6_AncHID5_-M1) or 14 (AncHID6_AncHID5_-M2) residues in 5 of the regions located in proximity to the substrate binding pocket was sufficient to recapitulate 71% or 63% of the TS activity of AncHID5, respectively (Figure 5e and **Table S3**). This result indicates that the low-level cyclase activity already present in AncHID6 could be readily enhanced through a relatively small number of amino acid substitutions.

In contrast, the de novo evolution of basal TS activity in an ancestral CXE is not as easy to explain. AncHID7 differs from AncHID6 at 59 positions (82% ID) and from AncHID5 at 95 positions (71% ID) in its amino acid sequence (**Figure S11**). Because AncHID6_AncHID5_-M1 exhibits much higher TS activity than wild-type AncHID6 and is more similar to AncHID7 at the sequence level than is AncHID5 (66 differences, 79% ID) (**Figure S22**), we generated and tested AncHID7_AncHID6(AncHID5)-M1_-M1 for cyclase activity. Despite replacing all 42 residues found in the 7 key regions surrounding the substrate binding pocket (**Figures S22**), we only observed very low levels of tabersonine in assays with angryline (Figure 5e and **Table S3**). Exchanging 12 additional residues (AncHID7_AncHID6(AncHID5)-M1_-M2) led to enhanced activity, with tabersonine levels matching those of the AncHID6_AncHID5_ mutants (Figure 5e and **Table S3**). However, we found that a shorter mutational pathway between high-level esterase and cyclase activity could be achieved if we employed AncHID6 as a template to generate a more active CXE. Replacing 35 residues in AncHID6 with the corresponding residues from AncHID7 fully endowed this enzyme (AncHID6_AncHID7_) with high-level esterase activity (**Figure S23** and **Table S4**). AncHID6_AncHID7_ and AncHID6_AncHID5_-M2 (TS) differ from one another at 41 positions (87% ID), all of which are confined to the 7 regions around the substrate binding pocket (**Figure S24**).

Notably, almost half of these changes appear in the α4/α5 loop and α5, highlighting this region as having played a particularly important role in cyclase evolution. Other critical modifications occurred in α1 and near the catalytic Asp and His residues, the latter conspicuously having been replaced with a Tyr as previously noted.

### Partial heterologous reconstitution of an ancestral MIA biosynthetic pathway

In order to place the evolution of the MIA cyclases in the broader context of pathway evolution, we considered how the substrate for these enzymes—dehydrosecodine (angryline)—might have arisen as a result of the promiscuous activity of other pathway enzymes. The biosynthesis of dehydrosecodine from the stable intermediate stemmadenine acetate requires the action of two enzymes: the FAD-dependent oxidase precondylocarpine acetate synthase (PAS) and the alcohol dehydrogenase (ADH) dihydroprecondylocarpine acetate synthase (DPAS) (13) (Figure 1). In our previous work, we noted that the heterologous host plant *Nicotiana benthamiana* harbors an enzyme that is capable of performing the same reaction as PAS (13, 17, 31). Moreover, the ADH geissoschizine synthase (GS) has been reported as a dual-function enzyme that can reduce precondylocarpine acetate to generate dehydrosecodine (15). Encouraged by these results, we infiltrated stemmadenine acetate into *N. benthamiana* leaves heterologously expressing GS or two other ADHs—heteroyohimbine synthase (HYS) and tetrahydroalstonine synthase (THAS)—that act upstream in the MIA pathway to generate corynanthe-type alkaloids (32, 33). As anticipated, we could detect small amounts of angryline in the leaf extracts, which was converted to tabersonine upon co-expression of extant and ancestral TS enzymes (Figures 6, **S25**, and **S26**). These results lend support to a plausible scenario in which enzymes from other metabolic pathways could, through adventitious reactivity toward upstream precursors, generate low levels of dehydrosecodine that an evolving CXE-like enzyme with cyclase activity could act upon. We imagine that the enzymes recruited for this new biosynthetic pathway would co-evolve to enable increasingly efficient production of aspidosperma alkaloids, secondary metabolites belonging to a new structural class that could provide a competitive advantage for the host plant.

**Figure 6.**
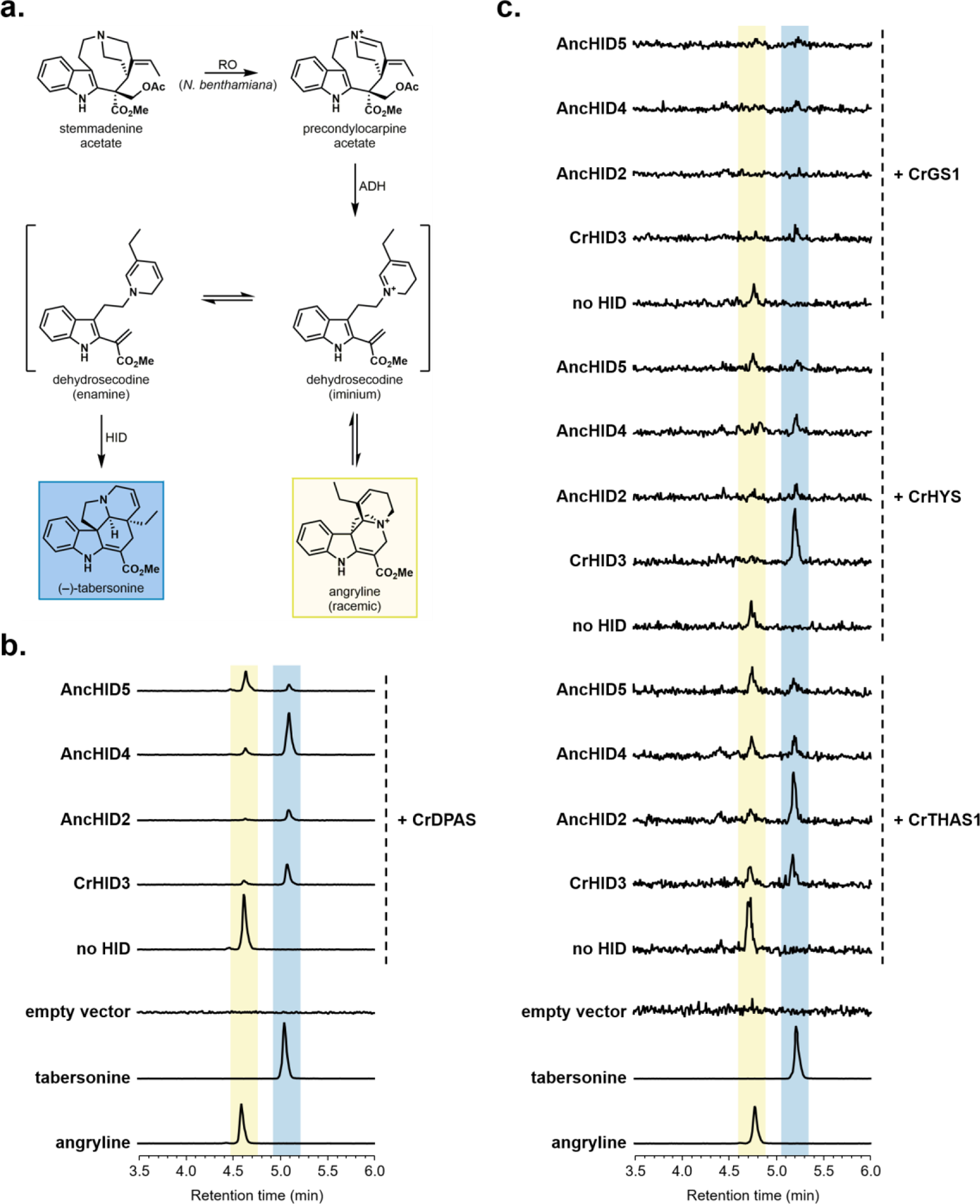
Reconstitution of dehydrosecodine (angryline) and tabersonine biosynthesis from stemmadenine acetate in *Nicotiana benthamiana*. (**a**) Biosynthesis of angryline and (–)-tabersonine from stemmadenine acetate. Enzyme abbreviations: RO (PAS-like reticuline oxidase), ADH (DPAS-like alcohol dehydrogenase), HID (2-hydroxyisoflavanone dehydratase-like cyclase). (**b**) LC-MS traces (EIC 337.1911 ± 0.01) of leaf extracts following infiltration of CrDPAS alone (no HID) or with extant (CrHID3) or ancestral (AncHID2, AncHID4, AncHID5) TS. (**c**) LC-MS traces (EIC 337.1911 ± 0.01; signal intensity magnified ∼10x) of leaf extracts following infiltration of the ADHs CrTHAS1, CrHYS, or CrGS1 alone (no HID) or with extant (CrHID3) or ancestral (AncHID2, AncHID4, AncHID5) TS. See **Figures S25** and **S26** for results from additional trials.

## Discussion

The aspidosperma and iboga classes of MIAs, which are composed of over 400 distinct molecules, have a wide range of important biological activities (5, 34, 35). Notably, production of these types of alkaloids is restricted to the Rauvolfioideae subfamily of the large and diverse Apocynaceae plant family. For decades, it had been postulated that the biogenetic construction of these scaffolds involves the action of enzymes catalyzing regiodivergent [4+2] cycloaddition reactions on the unstable intermediate dehydrosecodine (36, 37). The distinctive phylogenetic distribution of the aspidosperma/iboga alkaloids along with the recent identification and in vitro characterization of the cyclases directly responsible for their biosynthesis (13–18) motivated us to explore the evolution of this pathway using a combined phylogenetic and biochemical approach.

Phylogenetic analysis of the previously characterized MIA cyclases places them in a distinct clade among plant class I CXEs. With the exception of *Rauvolfia tetraphylla* (38, 39), all of the species represented in this clade (Clade 1 in Figure 2) are known to produce aspidosperma alkaloids. In contrast, among the Clade 1 species, production of iboga and pseudoaspidosperma alkaloids (both derived biosynthetically from 16-cmc) is limited to *C. roseus* (tribe Vinceae) and *T. iboga*, *Tabernaemontana divaricata*, and *Tabernaemontana elegans* (tribe Tabernaemontaneae). It is noteworthy that aspidosperma alkaloids are considerably more widespread than the iboga type, both in terms of number of known compounds and with respect to their distribution among species in the Apocynaceae family (40). Functional analysis of the Clade 1 proteins showed that only a subset of them (Clade 1a) are capable of cyclizing dehydrosecodine (angryline) to form catharanthine (CS), tabersonine (TS), or 16-cmc (CorS) (Figure 3). The product specificities of the cyclases are consistent with the chemotaxonomic distribution of the aspidosperma and iboga alkaloids: *C. roseus* and the three Tabernaemontaneae species each possess both a TS and a CorS while the remaining species only have a TS. In accordance with the fact that catharanthine is only produced by species in the *Catharanthus* genus, the only enzyme exhibiting CS activity is CrHID1 from *C. roseus*.

Furthermore, we noted two key sequence signatures that are peculiar to the cyclases: the His-Gly-Ala-Gly oxyanion hole and the Ser-(Tyr/Phe)-Asp catalytic triad (Figure 2b). Some residues can also be used to reliably distinguish between enzymes with TS or CorS activity (e.g., **Ser**-Thr and **Tyr** (TS) vs. **Ser**-Pro and **Phe** (CorS); bolded residues are part of the catalytic triad). While the Ser-His-Asp catalytic triad remains intact in all of the Clade 1c proteins (Figure 2b), these enzymes do not display any appreciable levels of esterase activity (**Table S4**). Instead, the closest extant relatives of the cyclases with significant hydrolytic capability are found in Clade 2, which consists of representative CXEs from the Rubiaceae.

On the basis of these results, we surmised that an ancestral CXE evolved into the first cyclase (TS), which subsequently diversified to give rise to the modern CS, TS, and CorS enzymes. Accordingly, TS activity represents an ancestral function from which CS and CorS activity subsequently evolved, indicating that the biosynthesis of aspidosperma alkaloids preceded that of iboga alkaloids. We were able to provide support for this hypothesis by resurrecting ancestral proteins represented at important nodes in the phylogeny and testing the catalytic activity of these enzymes with respect to hydrolysis and cyclization. Mutagenesis of selected ancestors in key regions of their amino acid sequence provided further insight into the molecular basis for cyclase evolution.

Here, we delineate a plausible trajectory for the evolution of the MIA cyclases that is supported by our data. The Rubiaceae family is thought to have diverged from the rest of the Gentianales around 87–92 million years ago (41). Shortly thereafter, in a lineage that would subsequently become the Apocynaceae family, a CXE (AncHID7) started to acquire a number of mutations that would ultimately lead to loss of its original catalytic function. At the same time, it serendipitously developed a latent TS activity (AncHID6) that was selected for and optimized in the Rauvolfioideae lineage. During this process, any remaining hydrolytic activity was lost due to substitution of Tyr for His in the catalytic triad. Enhancement of TS activity was concomitant with the appearance of low-level CorS activity as seen in AncHID5 and AncHID4. Gene duplication then enabled the specialization of one paralog for the production of 16-cmc (AncHID3), but only in the Tabernaemontaneae lineage. Over time, further specialization of TS also occurred, and orthologs finely tuned for this specific activity are likely to be found in all extant species that produce aspidosperma alkaloids. In *C. roseus* (or in an ancestral *Catharanthus* sp.), duplication of the ancestral TS (AncHID2) allowed for a neofunctionalized CorS enzyme with low activity (AncHID1) to emerge. A second duplication event then occurred, enabling simultaneous optimization of CorS activity (leading to CrHID2) and evolution of an enzyme with a completely novel function: CS (CrHID1).

This evolutionary trajectory combines various features of the classical neo- and subfunctionalization models for the evolution of new enzyme functions. Often, a multifunctional or generalist ancestor with the ability to carry out more than one type of reaction or to act on multiple different substrates is subfunctionalized following duplication to give separate specialists that each perform only one of the ancestral functions (42, 43). In many cases, the ancestor may not be multifunctional in the sense that its secondary activities provide a clear fitness advantage for the host organism. Rather, it may simply exhibit some level of promiscuity, which from an evolutionary biochemical perspective refers to the coincidental ability of an enzyme to perform side reactions that are physiologically irrelevant and thus not subject to natural selection (44–47). Although it can be difficult to establish the biological relevance of a given secondary enzymatic activity, many studies have described the co-option of a preexisting promiscuous or minor function as a means by which a novel physiologically relevant enzymatic activity can evolve (48–53).

How the first MIA cyclase evolved to catalyze a seemingly complicated cyclization reaction starting from an ancestral CXE is one of the central questions that motivated our study. We postulate that adventitious TS activity first arose in AncHID6 due to co-option of the oxyanion hole inherited from a CXE ancestor (AncHID7). In the likely stepwise cyclization mechanism employed by TS (Figure 3a), Michael addition of the dihydropyridine ring to the methyl acrylate portion of the substrate generates an oxyanion, which is stabilized by the oxyanion hole. The subsequent Mannich-like reaction to form the second C–C bond may then occur spontaneously in solution following product release or in the enzyme active site. We note that dehydrosecodine (angryline) can also cyclize spontaneously to form tabersonine under basic pH conditions, albeit at a slow rate (**Figure S27**). An enzyme promoting this reaction would thus be selected for if tabersonine and/or any downstream metabolites conferred an advantage on the host plant (*vide infra*).

Unlike tabersonine, the intermediate required for the biosynthesis of iboga and pseudoaspidosperma alkaloids (16-cmc) does not appear to form spontaneously in solution (**Figure S27**). We suspect that the ability of AncHID5 and AncHID4 to generate low levels of 16-cmc arose as an accidental consequence of optimizing the TS activity present in AncHID6. If this promiscuous activity provided an additional selective advantage, gene duplication enabling enhancement of CorS activity independent of TS activity would have been favored. Such a scenario is consistent with both the innovation-amplification-divergence (IAD) and escape from adaptive conflict (EAC) models of gene functional divergence (54, 55). In the IAD model, gene amplification resulting in increased protein expression would have led to higher levels of the advantageous metabolite (16-cmc). Positive selection for mutations that increased the efficiency of this initially promiscuous activity (e.g., T175P) would ultimately lead to the extant Tabernaemontaneae CorS enzymes. Furthermore, as tabersonine and 16-cmc derive from the same substrate, optimization of CorS activity would have likely come at the expense of TS activity. The resulting adaptive conflict would have also favored duplication and functional specialization of two separate paralogs.

Whereas almost all Apocynaceae species investigated in this study appear to harbor only one or (at most) two genes encoding cyclases, three separate duplication events ultimately gave rise to four cyclase genes (CrHID1–CrHID4) in *C. roseus*. Unlike AncHID4, the ancestral cyclase from this organism (AncHID2, TS) exhibits no detectable CorS activity. Instead, 16-cmc would only appear as a minor product in subsequent ancestors (i.e., sometime between AncHID2 and AncHID1), presumably following gene duplication and neofunctionalization of the TS (56). Our results demonstrate that a single amino acid substitution (T175P) is sufficient to enable low-level production of 16-cmc by AncHID2 (Figure 5d). A number of additional mutations lead to only a minor increase in CorS activity, which is concomitant with a significant loss of the original TS activity. Thus, AncHID1 displays low-level CorS activity while retaining ancestral TS activity as a promiscuous function. Following duplication of this ancestral gene, CorS activity is enhanced (largely via a Y300F mutation) in one copy, while CS activity appears to emerge de novo in the other copy. Interestingly, the extant *C. roseus* CorS (CrHID2) and CS (CrHID1) enzymes both retain residual TS activity, but the latter has completely lost the CorS activity of its direct ancestor (**Table S3**). As the minor levels of catharanthine detected in reactions with the extant and ancestral CorS enzymes are most likely due to spontaneous cyclization of 16-cmc under basic pH conditions (**Figure S28**), it is plausible that CS activity was a de novo innovation in accordance with the classical neofunctionalization model (56). The fourth *C. roseus* cyclase (CrHID4) arose from a third duplication event, but it degraded into a pseudogene following accumulation of neutral and detrimental (including nonsense and frameshift) mutations (**Figure S3**).

When considering the evolution of secondary metabolic pathways, it is tempting to speculate that the activity of the pathway enzymes evolved in a sequential manner. However, the substrate for the cyclases (dehydrosecodine) is highly reactive and likely does not accumulate to an appreciable extent in planta. Thus, it is difficult to imagine what kinds of selective pressures would have enabled the biosynthetic enzymes for production of dehydrosecodine to evolve before cyclization activity emerged. Since we found that *N. benthamiana* is able to convert stemmadenine acetate to precondylocarpine acetate, it is clear that oxidases can act promiscuously to non-specifically catalyze this reaction. We also demonstrated that extant ADHs acting upstream in the MIA pathway can generate small amounts of dehydrosecodine from precondylocarpine acetate (Figures 6, **S25**, and **S26**). As previously noted, non-enzymatic cyclization of this reactive intermediate can also occur under the right conditions to produce tabersonine (**Figure S27**). Therefore, it is possible to imagine a scenario in which dehydrosecodine and perhaps even low levels of tabersonine could first emerge from an “underground metabolism” (57, 58) consisting of reactions catalyzed by several substrate promiscuous enzymes as well as potential non-enzymatic transformations. Depending on the relative selective advantage provided by these new underground metabolites, recruitment of a CXE with latent TS activity could have occurred either during or after the first appearance of dehydrosecodine. The enzymes involved in this nascent pathway to produce aspidosperma alkaloids would have likely co-evolved to specialize in their new functions, and novel MIA metabolic pathways acting downstream of tabersonine could have also started to arise. Such a scenario is largely consistent with an integrated metabolite-enzyme coevolution model of pathway evolution (59).

Here, we have used a combination of phylogenetics, ancestral sequence reconstruction, biochemistry, and partial in vivo pathway reconstitution to probe the evolution of aspidosperma and iboga alkaloid biosynthesis in the Apocynaceae family. Our results support a model involving recruitment of multiple promiscuous enzymes from different pathways to generate the first aspidosperma alkaloid— tabersonine. Following enlistment of an ancestral CXE-like protein with low-level TS activity, we show that a relatively small number of mutations in and around the substrate binding pocket would have been sufficient to significantly increase this initial cyclase activity. Subsequent gene duplication events followed by sub- and neofunctionalization led to independent evolution of CorS activity in two different lineages, leading in turn to the iboga and pseudoaspidosperma classes of MIAs. CS activity arose last following duplication and neofunctionalization of an ancestral CorS enzyme, which occurred exclusively in the plant *C. roseus*. Overall, our work demonstrates how evolution has co-opted select members of a ubiquitous protein superfamily to catalyze unusual reactions in plant specialized metabolism and provides useful insight into the evolution of novel function in a unique class of biosynthetic enzymes.

## Acknowledgments

We would like to thank Benke Hong for his assistance with isolation of stemmadenine acetate, Delia Ayled Serna Guerrero, Sarah Heinicke, and Maritta Kunert for their assistance with LC-MS data acquisition, and members of the Max Planck Institute for Chemical Ecology greenhouse team for providing and caring for the plants. We also thank the following for generously providing cDNA used for amplifying some of the genes investigated in this study: Lorenzo Caputi, Mohamed O. Kamileen, Carsten Schotte, Evangelos C. Tatsis, Francesco Trenti, and Kotaro Yamamoto. We gratefully acknowledge the European Research Council (788301) and the Max Planck Society for funding.

## Notes

### Competing Interest Statement

The authors have declared no competing interest.

