## Supporting Information for "Evolution and Diversification of Carboxylesterase-like [4+2] Cyclases in Aspidosperma and Iboga Alkaloid Biosynthesis"

### Table of Contents


1. **Chemicals, reagents, plants, and miscellaneous items ... S2**
2. **Identification and selection of class I CXE gene sequences ... S2–S3**
3. **Phylogenetic analysis and ancestral sequence reconstruction ... S3–S4**
4. **Cloning and mutagenesis ... S4–S5**
   - General considerations ... S4
   - RNA isolation and cDNA generation ... S4
   - Cloning of genes ... S4
   - Generation of extant and ancestral HID mutants ... S5
5. **Protein expression and purification ... S5**
6. **Production and purification of angryline for activity assays ... S6**
7. **In vitro activity assays ... S6–S9**
   - Angryline cyclization assays (qualitative) ... S6
   - Angryline cyclization assays (quantitative) ... S6–S8
   - Ester hydrolysis assays ... S8–S9
8. **Pathway reconstitution in *Nicotiana benthamiana* ... S9–S10**
9. **LC-MS methods ... S10**
10. **Supplemental figures ... S11–S53**
    - Figure S1 ... S11
    - Figure S2 ... S12–S17
    - Figure S3 ... S18
    - Figure S4 ... S18
    - Figure S5 ... S19–S20
    - Figure S6 ... S20
    - Figure S7 ... S21
    - Figure S8 ... S22–S24
    - Figure S9 ... S25
    - Figure S10 ... S26
    - Figure S11 ... S27
    - Figure S12 ... S28
    - Figure S13 ... S29
    - Figure S14 ... S30
    - Figure S15 ... S31–S32
    - Figure S16 ... S33
    - Figure S17 ... S34–S35
    - Figure S18 ... S35
    - Figure S19 ... S36–S37
    - Figure S20 ... S37
    - Figure S21 ... S38
    - Figure S22 ... S39
    - Figure S23 ... S40
    - Figure S24 ... S41
    - Figure S25 ... S42
    - Figure S26 ... S43
    - Figure S27 ... S44
    - Figure S28 ... S45–S46
    - Figure S29 ... S47–S48
    - Figure S30 ... S48
    - Figure S31 ... S49
    - Figure S32 ... S50
    - Figure S33 ... S51
    - Figure S34 ... S52
    - Figure S35 ... S53
11. **References ... S54**

### 1. Chemicals, reagents, plants, and miscellaneous items

(+)-Catharanthine was purchased from Abcam. (–)-Tabersonine, 4-nitrophenol, 4-nitrophenyl butyrate, 4-methylumbelliferone, 4-methylumbelliferyl butyrate, and acetosyringone were purchased from Sigma-Aldrich. Vindoline was purchased from Chemodex. Carbenicillin and gentamicin were purchased from Formedium. Rifampicin and spectinomycin were purchased from Fisher Scientific. Isopropyl-β-D-thiogalactopyranoside (IPTG) was purchased from Calbiochem. Media components for bacterial growth were purchased from Formedium and Fisher Scientific. Deionized water was obtained from a Milli-Q IQ 7000 ultrapure water purification system (Merck Millipore) equipped with a Millipak 0.22 µm filter. For buffers, pH was monitored using an Orion Star A111 pH meter (Thermo Scientific) calibrated according to the manufacturer's specifications. Optical density ($OD_{600}$) was measured using a Fisherbrand cell density meter model 40 (Fisher Scientific). Unless otherwise specified, plants were grown in a greenhouse at 20–26 °C (day) / 18–20 °C (night) and 30–80% relative humidity. Plants were fertilized once per week with 0.1% Ferty 3 (Planta Düngemittel GmbH) and watered periodically as needed.

### 2. Identification and selection of class I CXE gene sequences

Sequences corresponding to HID and HID-like cyclase homologs (plant class I CXEs) were obtained through tBLASTn analysis of in-house and publicly available genomes and transcriptomes of primarily Gentianales plant species using CrHID3 as a search query (see **Table S5** for a list of all databases searched). Open reading frames were extracted from the top-ranking hits (selection parameters: E-value < 1 x $10^{-50}$; query coverage > 60–70%), and translated sequences were aligned with those of known HIDs (GmHIDH, GeHIDM) and HID-like cyclases (CrHID1–CrHID4, TiHID1, TiHID2) using MAFFT[1] (default parameters). Identical or near-identical and partial (short/truncated) sequences were identified and removed from the analysis. If the genomic and transcriptomic sequences of a given gene were not identical, the genomic sequence was kept and the transcriptomic sequence was discarded. With a set of unique top-ranking candidate genes in hand, the sequences were realigned using MAFFT, and a phylogeny was inferred using FastTree[2] (default parameters). Sequences appearing nearest to the known cyclases were given abbreviated names (e.g., AhHID1–AhHID4 from *Amsonia hubrichtii*). The placement of the sequences within the phylogeny was used to determine whether or not to keep these sequences for further analysis. Those falling into clades located further away from the cyclases than Clade 5 or the HID/HID-like clade were removed.

Additional sequences were acquired from the OneKP database[3,4] via tBLASTn using CrHID3 as a search query. A total of 14 homologs from the following Gentianales species were added to the set of genes used to build the phylogeny: *Apocynum androsaemifolium*, *Asclepias curassavica*, *Galium boreale*, *Holarrhena pubescens*, *Psychotria ipecacuanha* (renamed to *Carapichea ipecacuanha*), *Psychotria marginata*, *Wrightia natalensis*. As a number of top-ranking homologs from *Ipomoea* and *Solanum* spp. (Solanales) were also identified, we included the genomes and transcriptomes of *Ipomoea nil*, *Ipomoea trifida*, *Ipomoea triloba*, and *Solanum pennellii* in the collection of BLAST databases we searched. Along with *Camptotheca acuminata* (Cornales), these were the only non-Gentianales species we searched by BLAST. Top-ranking hits from the NCBI nucleotide collection (nr/nt) were identical or near-identical to the HID homologs identified in the following species: *Amsonia hubrichtii* (AhHID1), *Catharanthus roseus* (CrHID1–CrHID4), *Coffea canephora* (CcHID1–CcHID4), *Ipomoea nil* (InHID1–InHID3), *Tabernaemontana elegans* (TeHID1 and TeHID4), *Tabernanthe iboga* (TiHID1 and TiHID2), *Vinca minor* (VmHID1). The following two genes from *C. roseus* (found in both the genome (as three exons) and the transcriptome) were not included in the phylogeny: cra_locus_21316_iso_1_len_1379_ver_3 (did not amplify in PCR reactions), CRO_T020110 (the experimental sequence differed considerably from the predicted sequence, including frame-shifting insertions and deletions resulting in a premature stop codon).

Additional class I CXEs that had been described in previous studies were also included in the phylogeny. These included a representative set of gibberellin receptors (GID1s) from a total of 19 monocot and eudicot species, CXEs from *Arabidopsis thaliana* (AtCXE1–AtCXE20), 2-hydroxyisoflavanone dehydratases (HIDs) from Fabaceae spp., acylsugar acylhydrolases (ASHs) from *Solanum* spp., tuliposide-converting enzymes (TCEs) from *Tulipa gesneriana*, and a number of other previously described CXEs and CXE-like proteins from various plants. The closest homolog of CrHID3 from *Escherichia coli* (EcCXE) was included to serve as an outgroup. In total, 257 gene sequences (not including EcCXE) from 77 plant species were selected for further analysis (see **Table S1** for a full list with additional information; see **Dataset S1** for all nucleotide sequences).

Although HID/HID-like cyclase homologs were identified in the transcriptomes of the following species, they were not included in the phylogeny due to their high similarity to those from related species (shown in parentheses): *Calotropis procera* (*C. gigantea*), *Catharanthus longifolius* (*C. roseus*), *Catharanthus ovalis* (*C. roseus*), *Cinchona ledgeriana* (*C. pubescens*), *Ipomoea triloba* (*I. trifida*). No close homologs were identified in the transcriptomes of the following species: *Gentiana macrophylla*, *Hoodia gordonii*. In some cases, the sequences of the HID/HID-like cyclase homologs or other class I CXEs contained a premature stop codon or ambiguous nucleotides. Stop codons were replaced with appropriate sense codons chosen based on an alignment of the target sequence with its closest homologs. Premature stop codons (occurring only once in each sequence) were replaced in the following gene sequences: AcuHID1, AhHID4, CrHID4, GsHID1b, InHID5, MsHID4, SgHID2, UgHID3. Ambiguous nucleotides were similarly replaced in the following gene sequences: PiHID3, PrMC3, RstHID1–RstHID5. See **Table S1** for additional details.

#### 3. Phylogenetic analysis and ancestral sequence reconstruction

The protein sequences of the 258 genes acquired as described above were aligned using MAFFT v7.450[1] (algorithm: L-INS-i; scoring matrix: BLOSUM62; gap open penalty: 1.53; offset value: 0.123) in Geneious Prime 2019.2.3. Whenever possible, sequence-verified cloned gene sequences were used. A maximum likelihood phylogeny was inferred using IQ-TREE v1.6.12[5] with ModelFinder[6] (best-fit substitution model: JTT+I+G4) together with SH-aLRT (x1000) and ultrafast bootstrap[7] (UFBoot2, x1000) supports. The tree was rooted at EcCXE, and the cladogram/phylogram was visualized with iTOL v6[8] (see **Figures 2** and **S2**).

For ancestral sequence reconstruction, a more focused phylogeny consisting of 145 genes from Clades 1–5 along with those from the ASH/ASH-like and HID/HID-like clades was constructed. All terminal stop codons were removed from the corresponding nucleotide sequences, and short/truncated genes (<900 nt: PiHID3, PmHID1, SlASH2), long genes (>1020 nt: AcuHID3, AsHID3, CgHID3), and genes aligning poorly with others internally (AcuHID1, AsHID1, CgHID1) or at the termini (InHID5, InHID6, ItHID3) were discarded. The remaining gene sequences were aligned (translation align) using MAFFT v7.450 (algorithm: L-INS-i; scoring matrix: BLOSUM62; gap open penalty: 1.53; offset value: 0.123) in Geneious Prime 2019.2.3, and a codon-based maximum likelihood phylogeny was inferred using IQ-TREE v1.6.12 with ModelFinder (best-fit substitution model: GY+F1X4+I+G4) together with SH-aLRT (x1000) and ultrafast bootstrap (UFBoot2, x1000) supports. The tree was rooted at the superclade containing the Clade 5 CXEs and the Fabaceae HIDs, and the phylogram was visualized with iTOL v6 and FigTree v1.4.4 (http://tree.bio.ed.ac.uk/software/figtree/) (see **Figures 4** and **S8**).

The codon-based alignment and focused phylogeny of 145 gene sequences were used for ancestral sequence reconstruction using the FastML server[9] and the Yang codon-substitution model. The most probable sequences (marginal reconstruction) at nine nodes corresponding to ancestral HID-like enzymes (AncHID1–AncHID9) were selected for further analysis. The presence of insertions or deletions in the reconstructed sequences was inferred manually by parsimony.[10] The ancestral genes were synthesized (Twist Bioscience) and cloned into the pOPINF vector for expression in *E. coli* (see “Cloning and mutagenesis”). See **Dataset S2** for the nucleotide sequences of AncHID1–AncHID9.

### 4. Cloning and mutagenesis

*General considerations*

DNA oligonucleotides for cloning and mutagenesis were purchased from Integrated DNA Technologies (IDT). Oligonucleotides ≤60 bp in length were ordered as 25 nmole DNA oligos whereas those >60 bp in length were ordered as 4 nmole Ultramer DNA oligos. All primers used for cloning and mutagenesis are listed in **Table S6**. PCR reactions were performed using the 2x Platinum SuperFi (II) PCR Master Mix (Invitrogen) and an Applied Biosystems SimpliAmp thermal cycler (Thermo Fisher Scientific). Restriction endonucleases were purchased from New England Biolabs. DNA concentrations were measured using an Implen NanoPhotometer N60 spectrophotometer. Sequencing services were provided by Genewiz (Azenta Life Sciences).

*RNA isolation and cDNA generation*

Leaf, stem, and root tissues were harvested from plants of interest and immediately flash frozen in liquid $N_2$. The combined tissues from each plant were ground to a fine powder under liquid $N_2$ using a mortar and pestle. Total RNA was extracted and purified from ~100 mg of frozen tissue powder from each plant according to the established CTAB method.[11,12] DNA was removed from the RNA samples using the TURBO DNA-*free* Kit (Invitrogen), and cDNA was synthesized using the SuperScript IV VILO Master Mix (Invitrogen) according to the manufacturer’s instructions. cDNA from several additional plants was kindly provided by our colleagues (see Acknowledgments).

*Cloning of genes*

All genes of interest were cloned into pOPINF for heterologous expression in *E. coli* or into a modified version of the binary vector p3Ω1[13] for heterologous expression in *Nicotiana benthamiana*. Genes were amplified by PCR using the primers listed in **Table S7**. All primers contained 15- to 20-bp In-Fusion compatible overhangs to enable subsequent cloning into the target vector. Amplicons obtained using cDNA as template were isolated via gel extraction followed by cleanup and purification using the NucleoSpin Gel and PCR Clean-up kit (Macherey-Nagel) according to the manufacturer’s instructions. Reactions performed using a synthetic gene (Twist Bioscience) as template were directly subjected to PCR cleanup using the same kit. When a plasmid was used as template, the completed PCR reaction was incubated with DpnI (20 U) at 37 °C for 1 h prior to purification using the same kit. Purified amplicons were cloned into pre-digested pOPINF (KpnI/HindIII) or p3Ω1 (BsaI) using the In-Fusion cloning kit (Takara) according to the manufacturer’s instructions. In-Fusion reactions were transformed into chemically competent Stellar or TOP10 *E. coli* cells prior to selection on LB-agar supplemented with carbenicillin (100 µg/mL, pOPINF) or spectinomycin (200 µg/mL, p3Ω1). Typically, 3–5 colonies were selected for overnight growth and plasmid isolation via miniprep using the Wizard Plus SV Minipreps DNA Purification System (Promega). Plasmid constructs were sequenced to verify the identity of the cloned gene and/or to ensure that no mutations were present in the open reading frame. The following primers were used for sequencing reactions: pOPINF (forward: TAATACGACTCACTATAGGG, reverse: TAGCCAGAAGTCAGATGCT); p3Ω1 (forward: GATGAAAAAGCCCTAAAATTGGAG, reverse: ATTATTCACAAATGAGAAACAGAATGG). When multiple isoforms of a gene cloned from cDNA were observed, the one exhibiting the fewest differences from the predicted sequence and/or the one present in highest abundance was selected for further investigation.

*Generation of extant and ancestral HID mutants*

HID mutants were acquired either by ordering fully synthetic genes (Twist Bioscience) or by using mutagenic primers and splicing 2–4 overlapping PCR fragments via In-Fusion. Full genes and gene fragments were amplified by PCR using the primers and templates listed in **Table S8**. When multiple PCR fragments were simultaneously cloned into pOPINF, primers were designed to contain 20-bp In-Fusion compatible overhangs. Otherwise, primers contained 15-bp overhangs. Upon PCR reaction completion, amplicons were purified and cloned into pOPINF (pOPINM for CrHID4 only) using the In-Fusion cloning kit (Takara) as described above (note that up to four fragments could be simultaneously spliced and cloned into the target vector—see **Table S8** for details). Competent cells were transformed, and plasmids were isolated and sequenced to ensure the correct identity of the mutated gene. The following primers were used for sequencing reactions involving pOPINM: forward: GAAATCATGCCGAACATC, reverse: TAGCCAGAAGTCAGATGCT. See **Dataset S2** for the nucleotide sequences of all HID mutants.

### 5. Protein expression and purification

The pOPINF or pOPINM plasmid containing the gene of interest was transformed into chemically competent SoluBL21 *E. coli* cells, and an individual colony was selected for overnight growth at 37 °C in 5 mL of 2x YT supplemented with carbenicillin (100 µg/mL). 100 mL of 2x YT (250 mL Erlenmeyer flask) supplemented with carbenicillin (100 µg/mL) was inoculated with 1 mL of the overnight seed culture and incubated at 37 °C (200 rpm). When the $OD_{600}$ reached 0.6–1.0 (~3.5 h post inoculation), the culture was cooled in an ice water bath for 5–10 min prior to addition of IPTG (200 µM). The culture was grown at 18 °C for 20–24 h before the cells were harvested by centrifugation (4000 rpm, 15 min, 4 °C) and stored at -80 °C until needed for protein purification.

The cells were thawed and resuspended in 10 mL of Buffer A1 (50 mM Tris, 50 mM glycine, 5% (v/v) glycerol, 500 mM NaCl, 20 mM imidazole, pH 8.0) supplemented with 0.5 mg/mL lysozyme (Sigma-Aldrich) prior to lysis via sonication (3 s *on* / 5 s *off*, 2 min total *on* time, 40% amplitude) using a Vibra-Cell Ultrasonic Processor (Sonics & Materials, Inc.). The cell lysate was centrifuged at 40,000 x g for 30 min (4 °C), and the clarified lysate was decanted into a 15 mL Falcon tube and incubated with 0.25 mL of pre-equilibrated His60 Ni Superflow Resin (Takara) at 4 °C for 1 h on a nutating shaker. The resin was pelleted via centrifugation (4000 rpm, 2 min, 4 °C) and washed with 3 x 10 mL of Buffer A1. After the last round of washing, protein was eluted by addition of 2 x 0.5 mL of Buffer B1 (50 mM Tris, 50 mM glycine, 5% (v/v) glycerol, 500 mM NaCl, 500 mM imidazole, pH 8.0) to the resin. Following the elution steps, the combined eluate was centrifuged at 16,900 x g for 2 min (4 °C) to remove any remaining resin. The protein was exchanged into Buffer A4 (20 mM HEPES, 150 mM NaCl, pH 7.5) using a PD-10 desalting column (Cytiva) and then concentrated using a 4 mL 10 kDa MWCO Millipore Amicon Ultra centrifugal filter (4000 rpm, 25 min, 4 °C). Further concentration using a 0.5 mL 10 kDa MWCO centrifugal filter (14,000 x g, 10 min, 4 °C) was performed if necessary. After dispensing into small aliquots, the concentrated protein was flash frozen in liquid $N_2$ and stored at -80 °C until used for assays. Protein concentration was determined using a NanoDrop One spectrophotometer (Thermo Scientific) and the molecular weight and $\varepsilon_{280}$ values calculated using the ExPASy ProtParam tool. Most proteins were purified on at least two separate occasions, and each batch was assessed for cyclase and carboxylesterase activity as described below (see “In vitro activity assays”).

The following proteins exhibited very low levels of soluble expression (<1 mg/L), precluding their in vitro analysis: MsHID2, VmHID3. None of the cloned genes from Clade 5 were tested for expression (CrHID7, TeHID7, TiHID8). AhHID4 contained a premature stop codon and was therefore also not tested for expression. The premature stop codon present in CrHID4 was removed (see **Figure S3**), but this protein only exhibited reasonable soluble expression levels with an N-terminal MBP tag.

### 6. Production and purification of angryline for activity assays

Angryline was generated from stemmadenine acetate as previously described with minor modifications.[14] Stemmadenine acetate was obtained as previously described.[15] CrPAS and CrDPAS were expressed in *N. benthamiana* and *E. coli*, respectively, and purified as previously described.[16,17] Reaction conditions for preparation of angryline were as follows: 2.5 mM stemmadenine acetate (9.7 mg, dissolved in methanol), 2 µM CrPAS, 100 µM FAD disodium salt hydrate (Sigma), 5 µM CrDPAS, 3 mM NADPH tetrasodium salt (Roche), 50 mM Tris (pH 8.5). First, stemmadenine acetate (methanol = 10% final reaction volume), CrPAS, FAD, and Tris buffer were combined in a 50 mL Falcon tube and incubated at 37 °C for 2 h (150 rpm). Then, CrDPAS and NADPH were added, and the reaction (total volume = 9.8 mL) was further incubated at 37 °C for 30 min (150 rpm). Upon completion (assessed by LC-MS), the reaction mixture was split into 0.5 mL aliquots, which were flash frozen in liquid $N_2$ and stored at -80 °C until purification by preparative HPLC.

All solvents used for HPLC purification were HPLC grade and purchased from Fisher Scientific. Just prior to HPLC purification, each 0.5 mL aliquot of the reaction mixture containing angryline was thawed, combined with 0.5 mL of 90:9:1 $CH_3OH$:$H_2O$:formic acid, and filtered through 0.22 µm nylon Costar Spin-X centrifuge tube filters (Corning) via centrifugation (16,000 x g, 1 min, room temperature). HPLC purification was performed on an Agilent 1260 Infinity II preparative-scale instrument with UV detection at 330 nm using a Phenomenex Kinetex XB-C18 column with the following specifications: dimensions, 250 x 10 mm; particle size, 5 µm; pore size, 100 Å. HPLC conditions were as follows: mobile phase (A = deionized water + 0.1% formic acid, B = acetonitrile + 0.1% formic acid); 10% to 30% B over 16 min, 30% to 90% B over 2 min, 90% B for 3 min, 90% to 10% B over 0.1 min, 10% B for 3.9 min; flow rate, 6.0 mL/min; injection volume, 1.0 mL. Fractions (retention time = 11.0–13.5 min) were combined and completely dried under reduced pressure. The residue was resuspended in methanol and transferred to a massed glass vial. The methanol was removed under reduced pressure, yielding angryline (formate salt) as a transparent light yellow amorphous solid (7.1 mg, 76% yield from stemmadenine acetate). The material was dissolved in 100 mM MES (pH 6.0) to a concentration of 30 mM and stored at -20 °C until used for enzymatic assays.

### 7. In vitro activity assays

*Angryline cyclization assays (qualitative)*

Analytical-scale enzymatic reactions for qualitative assessment of angryline cyclization activity were performed under the following conditions: 1–5 µM enzyme, 50 µM angryline, 50 mM Tris (pH 9.0). Reactions were performed in 1.5 mL Eppendorf tubes (safe-lock), and the total volume of each reaction was 100 µL. To set up the reactions, the following reagents were added to each tube in the order given: deionized water (35 µL), 100 mM Tris (pH 9.0, 50 µL), 10x enzyme in Buffer A4 (10 µL), 1 mM angryline (5 µL). Reactions were incubated at 37 °C for 30 min (800 rpm) prior to quenching by addition of 200 µL of 90:9:1 $CH_3OH$:$H_2O$:formic acid. The quenched reaction mixtures were centrifuged (17,000 x g, 10 min, 4 °C), and the supernatant was removed for LC-MS analysis. All reactions were performed and analyzed at least twice. Buffer A4 (10 µL) was substituted for enzyme in negative control reactions performed in parallel.

*Angryline cyclization assays (quantitative)*

Analytical-scale enzymatic reactions for quantitative assessment of angryline cyclization activity were performed as described above with some minor modifications. The final concentration of enzyme in the reactions ranged from 40 nM to 5 µM (see **Table S3**), and reactions were quenched with 90:9:1 $CH_3OH$:$H_2O$:formic acid containing 1 µM vindoline as internal standard. All reactions were performed and analyzed independently four times.

To enable quantification of substrate and product concentrations for determination of turnover number (TON), calibration curves were generated for angryline, 16-cmc, catharanthine, and tabersonine both prior to and following LC-MS analysis of each set of reactions in a given trial. Concentrations (µM) of standards analyzed were as follows: 1, 2, 4, 8, 16, 24, 32, 40. For angryline, 5 µL of 20x stocks prepared from a 1 mM solution in 100 mM MES (pH 6.0) was added to 95 µL of stock solvent A (35 µL deionized water + 50 µL 100 mM Tris (pH 9.0) + 10 µL Buffer A4) to prepare the dilution series. For catharanthine and tabersonine, 2 µL of 50x stocks prepared from a 2 mM solution in DMSO was added to 98 µL of stock solvent B (34.3 µL deionized water + 49 µL 100 mM Tris (pH 9.0) + 9.8 µL Buffer A4 + 4.9 µL 100 mM MES (pH 6.0)) to prepare the dilution series. As 16-cmc is not isolable as a stable compound, it was generated enzymatically under the following conditions: 10 µM TiHID2, 50 µM angryline, 50 mM Tris (pH 9.0). The reaction (total volume = 300 µL) was split into 3 x 100 µL aliquots and incubated at 37 °C for 1 h (800 rpm). Upon completion, aliquots of the reaction mixture were added to stock solvent B to prepare the dilution series (volumes were calculated based on the starting concentration of angryline). Each of the resulting 100 µL solutions was quenched by addition of 200 µL of 90:9:1 $CH_3OH$:$H_2O$:formic acid containing 1 µM vindoline as internal standard.

Extracted ion chromatograms were generated to quantify substrate/products (*m/z* = 337.1911 ± 0.01) and internal standard (*m/z* = 457.2333 ± 0.01). All relevant peaks were integrated, and raw peak areas were normalized by dividing by the peak area of the internal standard from the same sample. To generate the calibration curve for angryline, normalized peak area vs. concentration was fit to a standard linear regression with y-intercept = 0. For catharanthine and tabersonine, normalized peak area vs. concentration was fit to the following quadratic equation (*y* = normalized peak area, *x* = concentration):

$$y = \frac{a\left[(b + x + c) - \sqrt{(b + x + c)^2 - 4bx}\right]}{2b}$$

To generate the calibration curve for 16-cmc, the normalized peak areas for residual angryline, catharanthine, and tabersonine observed in each sample of the TiHID2 reaction dilution series were first converted to concentration using the above equations. These values were then subtracted from the theoretical starting concentration of angryline in each sample (i.e., 1–40 µM) to obtain the estimated 16-cmc concentration. Normalized 16-cmc peak area vs. concentration was fit to a standard linear regression with y-intercept = 0 to generate the 16-cmc calibration curve. See **Figure S29** for the calibration curves (combined across all trials) generated for angryline, 16-cmc, catharanthine, and tabersonine.

Concentrations of angryline, catharanthine, and tabersonine in each of the enzymatic reactions were determined from the corresponding normalized peak areas using the linear and quadratic equations obtained from the calibration curves as described above. As discussed below, concentrations of 16-cmc could only be *estimated* based on the calibration curves generated for this compound and angryline. A fundamental assumption made in generating the 16-cmc calibration curve is that all of the angryline added to the TiHID2 enzymatic reaction is converted only to 16-cmc, catharanthine, and tabersonine. However, angryline is highly unstable under basic conditions and readily degrades into non-quantifiable products over time. Consequently, the combined concentrations of the main cyclization products (16-cmc, catharanthine, tabersonine) and remaining starting material (angryline) will always be lower than the starting concentration of angryline in the reaction mixture. Indeed, it is apparent that a significant decrease in angryline is concomitant with only minor increases in 16-cmc and the other quantifiable cyclization products when the TiHID2 reaction is allowed to run for longer than 30 min (**Figure S28**). Since less angryline than anticipated is converted to 16-cmc, the concentration of 16-cmc determined from the corresponding calibration curve would be an overestimate. Moreover, because the complete mass balance of the reaction cannot be accounted for, the 16-cmc concentration cannot be accurately determined simply by subtracting the final concentrations of catharanthine, tabersonine, and residual angryline from the starting concentration of angryline. Doing so would similarly lead to an overestimate of the 16-cmc concentration. It would also result in gross inaccuracies when trying to quantify low levels of 16-cmc.

One possible solution to this problem would be to simply use the angryline calibration curve instead of the 16-cmc calibration curve to directly calculate the 16-cmc concentration. This strategy would work only if angryline and 16-cmc exhibit equal ionization efficiencies. The maximum area of the peak corresponding to 16-cmc in reactions of CorS with 50 μM angryline is consistently lower than that corresponding to the same amount of unreacted angryline. This observation may be a consequence of the partial degradation of angryline as well as of 16-cmc during the course of the reaction. However, it may also be due to a slightly lower ionization efficiency of 16-cmc relative to angryline under the conditions we employ for LC-MS analysis. In this case, using the angryline calibration curve for quantification of 16-cmc concentration would yield an underestimated value. As the 16-cmc concentration calculated using the 16-cmc or angryline calibration curves would be an overestimate or an underestimate of the true value, respectively, we ultimately concluded that a reasonable compromise was to take the average of these two quantities in estimating the concentration of 16-cmc in a given reaction mixture.

The concentrations of 16-cmc, catharanthine, and tabersonine in the negative control reaction lacking enzyme ([16-cmc] = 0 μM, [catharanthine] < 0.1 μM, [tabersonine] < 0.1 μM) were subtracted from those in each experimental reaction, and the resulting values were divided by the enzyme concentration to obtain TON (mol product/mol enzyme). The results of all angryline cyclization assays are presented in **Table S3**.

*Ester hydrolysis assays*

Rates of ester hydrolysis were evaluated using either an absorbance-based (4-nitrophenyl butyrate, 4-NPB) or a fluorescence-based (4-methylumbelliferyl butyrate, 4-MUB) assay performed in 96-well plate format and a CLARIOstar Plus microplate reader (BMG Labtech). All reactions were performed under the following conditions: 5 nM to 5 μM enzyme, 500 μM substrate, 50 mM Tris (pH 8.0). Reactions were performed in 96-well plates (absorbance measurements: CytoOne clear, flat/clear bottom (STARLAB Int.); fluorescence intensity measurements: Nunc A/S black, flat/opaque bottom (Thermo Fisher Scientific)), and the total volume of each reaction was 200 μL. To set up the reactions, a mixture of substrate in reaction buffer (75 μL deionized water + 100 μL 100 mM Tris (pH 8.0) + 5 μL 20 mM substrate (DMSO stock) per well) was added via multichannel pipette to 12 wells at a time and incubated at 37 °C for 5 min. Then, 20 μL of 10x enzyme in Buffer A4 was added to each well to initiate the reaction, and the plate was immediately placed in the plate reader (temperature maintained at 37 °C for the duration of the measurements). In the negative control reaction performed in parallel, 20 μL of Buffer A4 substituted for enzyme. For reactions with 4-NPB, absorbance at 405 nm was monitored every 15 s for a total of 10 min. For reactions with 4-MUB, fluorescence intensity ($\lambda_{ex}$ = 365 nm, $\lambda_{em}$ = 445 nm) was monitored every 15 s for a total of 10 min. All reactions were performed and analyzed independently four times.

To enable quantification of product concentrations for determination of turnover frequency (TOF ($min^{-1}$) = μM product/min per μM enzyme), calibration curves were generated for hydrolysis products 4-nitrophenol and 4-methylumbelliferone by monitoring absorbance or fluorescence intensity at the following concentrations (μM): 0 (DMSO), 5, 10, 20, 30, 40, 50, 60, 70, 80, 90, 100, 120, 180, 240, 320, 400, 500. To prepare the dilution series, 5 μL of 40x stocks prepared from 4 mM and 20 mM solutions in DMSO was added to 175 μL of stock solvent C (75 μL deionized water + 100 μL 100 mM Tris (pH 8.0)). The plate was incubated at 37 °C for 5 min before adding 20 μL of Buffer A4 to each well. Absorbance or fluorescence intensity was monitored over 10 min, and the average value over this time interval was determined for each concentration tested. Signal vs. concentration was fit to the quadratic equation used to generate the calibration curves for catharanthine and tabersonine (see above; $y$ = absorbance or fluorescence intensity, $x$ = concentration). See **Figure S29** for the calibration curves generated for 4-nitrophenol and 4-methylumbelliferone.

Concentrations of hydrolysis products in each of the enzymatic reactions were interpolated from the corresponding signal intensities using the calibration curves generated as described above. Initial rates of hydrolysis were determined by fitting product concentration vs. time to either a standard linear regression or the following logarithmic function ($y$ = concentration, $x$ = time):

$$y = y_0 + b \ln\left(1 + \frac{x}{x_0}\right)$$

This equation is an approximation of the integrated form of the Michaelis-Menten equation at initial time points.[18] The initial rate is the derivative of $y$ with respect to $x$ when $x = 0$, or $b/x_0$. When curve fitting was not possible due to substantial progression of the reaction at $x = 0$, the initial rate was estimated using the following equation:

$$estimated\ rate = \frac{y_0^{enzymatic} - y_0^{buffer}}{x_i}$$

Here, $y_0$ refers to the concentration of hydrolysis product at the first time point in either the enzymatic reaction or the negative control reaction (buffer), and $x_i$ refers to the time from true reaction initiation (addition of enzyme) to first read ($x_{i(abs)}$ = 1 min, $x_{i(fluor)}$ = 1.3 min). If the product concentration at the first time point ($y_0$) was calculated to be >500 µM, $y_0$ was set to 500 µM. If the estimated rate was larger than the rate determined from linear or nonlinear regression analysis and $y_0$ > 100 µM, the estimated rate was used as the final value. The spontaneous rate of hydrolysis (negative control reaction) was subtracted from all final rate values, and these corrected rates were divided by the enzyme concentration to obtain TOF. The results of all ester hydrolysis assays are presented in **Table S4**.

### 8. Pathway reconstitution in *Nicotiana benthamiana*

*Nicotiana benthamiana* plants used for transient gene expression and pathway reconstitution were grown on a standard soil mix in a greenhouse under a 14 h light / 10 h dark photoperiod at 23–26 °C (day) / 16–22 °C (night) and 40–70% relative humidity. Plants were grown for four weeks prior to transfer into a controlled environment growth chamber and infiltration with *Agrobacterium tumefaciens* strains. In the growth chamber, plants were grown under a 16 h light / 8 h dark photoperiod at 21 °C and 35–60% relative humidity. Plants were watered periodically as needed.

Genes to be heterologously expressed in *N. benthamiana* were cloned into p3Ω1 as described above (see "Cloning and mutagenesis"). Plasmids were transformed into chemically competent *A. tumefaciens* GV3101 cells using the freeze-thaw method[19] prior to selection on LB-agar supplemented with rifampicin (50 µg/mL) + gentamicin (50 µg/mL) + spectinomycin (200 µg/mL). Positive transformants (verified by colony PCR using the p3Ω1 sequencing primers) were inoculated into 10 mL of LB supplemented with rifampicin (50 µg/mL) + gentamicin (50 µg/mL) + spectinomycin (200 µg/mL) and grown at 28 °C for 2 days. Subsequently, the cultures were centrifuged (4000 rpm, 10 min), the supernatant was removed, and the cells were washed with 10 mL of infiltration buffer (50 mM MES hydrate, 2 mM $Na_3PO_4$•$12H_2O$, 10 mM $MgCl_2$, 0.5% (w/v) D-(+)-glucose, 100 µM acetosyringone). The cells were centrifuged again (4000 rpm, 10 min) and resuspended in 10 mL of fresh infiltration buffer following removal of the supernatant. $OD_{600}$ was measured for each sample (diluted 10x), and individual strains were combined and further diluted such that the final $OD_{600}$ = 0.3 for each strain in a total volume of 4 mL. Following incubation at room temperature for ≥1 h, strain mixtures were infiltrated into four-week-old *N. benthamiana* plants. Infiltrations were performed on the abaxial side of leaves (2–4 per plant) using a needleless 1 mL syringe. After 3 days (~67–69 h post infiltration), stemmadenine acetate (50 µM in deionized water + 1% DMSO) was infiltrated into the leaves, and the plants were left to incubate for an additional 2 days (~48–49 h). Tissue was harvested by cutting 4 disks (1 cm diameter) from each leaf, placing them inside a 2 mL Eppendorf tube (safe-lock) containing a tungsten carbide bead (3 mm diameter, Qiagen), and immediately flash freezing in liquid $N_2$. Frozen tissue was stored at -80 °C until extracted for analysis.

For metabolite extraction, the frozen leaf tissue was disrupted using a TissueLyser II instrument (Qiagen) set at a frequency of 20 $s^{-1}$ for a total of 2 min. Immediately thereafter, 400 µL of LC-MS grade methanol was added to each sample, and the tubes were vortexed, sonicated for 15 min, vortexed again, and centrifuged (17,000 x g, 5 min, room temperature) to remove bulk tissue material. The supernatant was removed, incubated at 4 °C for 1–2 h, and centrifuged again (16,900 x g, 10 min, 4 °C) prior to filtration through a 0.2 µm PTFE syringe filter. The samples were then analyzed by LC-MS as described below. The entire procedure with all plasmid constructs and *A. tumefaciens* strains was performed on three separate occasions (results for trials 1–3 are shown in **Figures 6**, **S25**, and **S26**).

### 9. LC-MS methods

All solvents used for LC-MS analysis were LC-MS grade and purchased from Fisher Scientific. LC-MS analysis of in vitro enzymatic reactions was performed on a Thermo Scientific UltiMate 3000 UHPLC system coupled to a Bruker Impact II UHR-Q-TOF mass spectrometer equipped with an electrospray ionization (ESI) source using a Waters ACQUITY UPLC BEH C18 column (5 x 2.1 mm VanGuard pre-column) with the following specifications: dimensions, 50 x 2.1 mm; particle technology, BEH; particle size, 1.7 µm; pore size, 130 Å. HPLC conditions were as follows: mobile phase (A = 5 mM ammonium acetate (pH 6.0), B = acetonitrile); 20% B for 1 min (diverted to waste), 20% to 30% B over 2 min, 30% to 90% B over 3 min, 90% to 100% B over 0.1 min, 100% B for 1.4 min, 100% to 20% B over 0.1 min, 20% B for 1.4 min; flow rate, 0.6 mL/min; column oven temperature, 40 °C; injection volume, 1 µL. LC-MS traces for all in vitro enzymatic reactions are shown in **Figures 3**, **S3**, **S4**, **S6**, **S7**, and **S30–S34**.

LC-MS analysis of *N. benthamiana* extracts was performed on the same instruments using a Phenomenex Kinetex XB-C18 column (2 x 2.1 mm SecurityGuard ULTRA pre-column) with the following specifications: dimensions, 100 x 2.1 mm; particle size, 2.6 µm; pore size, 100 Å. HPLC conditions were as follows: mobile phase (A = deionized water + 0.1% formic acid, B = acetonitrile + 0.1% formic acid); 10% B for 1 min (diverted to waste), 10% to 30% B over 5 min, 30% to 100% B over 0.1 min, 100% B for 1.4 min, 100% to 10% B over 0.1 min, 10% B for 2.4 min; flow rate, 0.6 mL/min; column oven temperature, 40 °C; injection volume, 2 µL.

For MS data acquisition, ionization was performed via pneumatic-assisted ESI in positive mode with a capillary voltage of 3,500 V and an end plate offset of 500 V. The nebulizer pressure was set to 2.5 bar, and nitrogen (temperature, 250 °C; flow rate, 11 L/min) was used as the drying gas. Data were recorded at a rate of 12 Hz in the range *m/z* = 80–1000 with data-dependent tandem MS (MS2) and an active exclusion window of 0.2 min. For MS2, fragmentation was triggered on an absolute threshold of 400 counts and limited to a total cycle time range of 0.5 s. Collision energy (20–50 eV) was determined automatically using the stepping option model. MS2 spectra of (–)-16-cmc, angryline, (+)-catharanthine, and (–)-tabersonine are shown in **Figure S35**. For calibration of recorded MS spectra, a sodium formate-isopropanol solution was directly injected into the source at 0.18 mL/h using an automated syringe pump at the beginning of each sample run. Bruker Compass DataAnalysis (Version 5.3) software was used for subsequent data analysis.

### 10. Supplemental figures

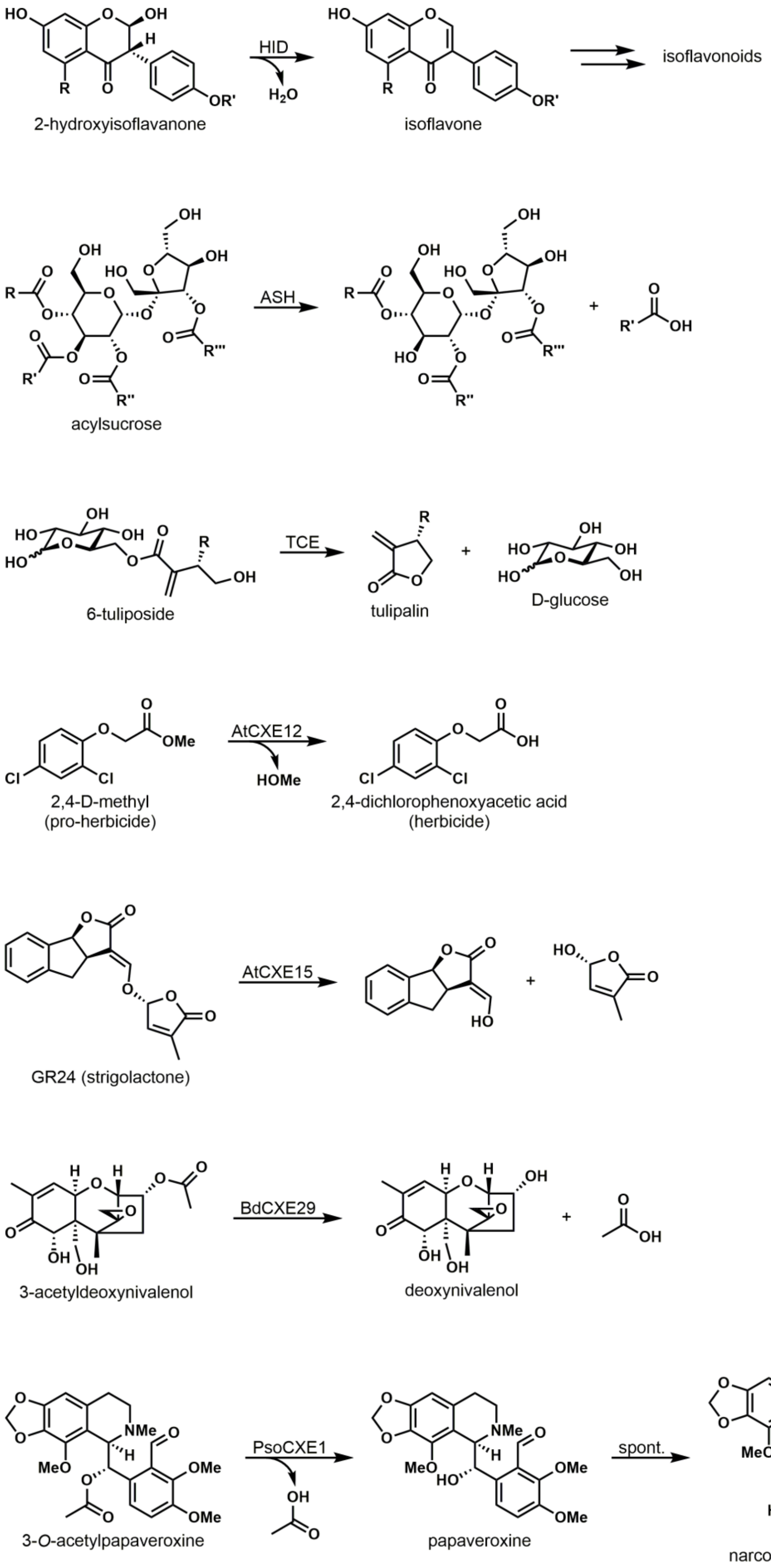


**Figure S1.** Selected examples of physiologically relevant reactions catalyzed by plant class I carboxylesterases (CXEs). Enzyme abbreviations: HID (2-hydroxyisoflavanone dehydratase), ASH (acylsugar acylhydrolase), TCE (tuliposide-converting enzyme), AtCXE (*Arabidopsis thaliana* CXE), BdCXE (*Brachypodium distachyon* CXE), PsoCXE (*Papaver somniferum* CXE).

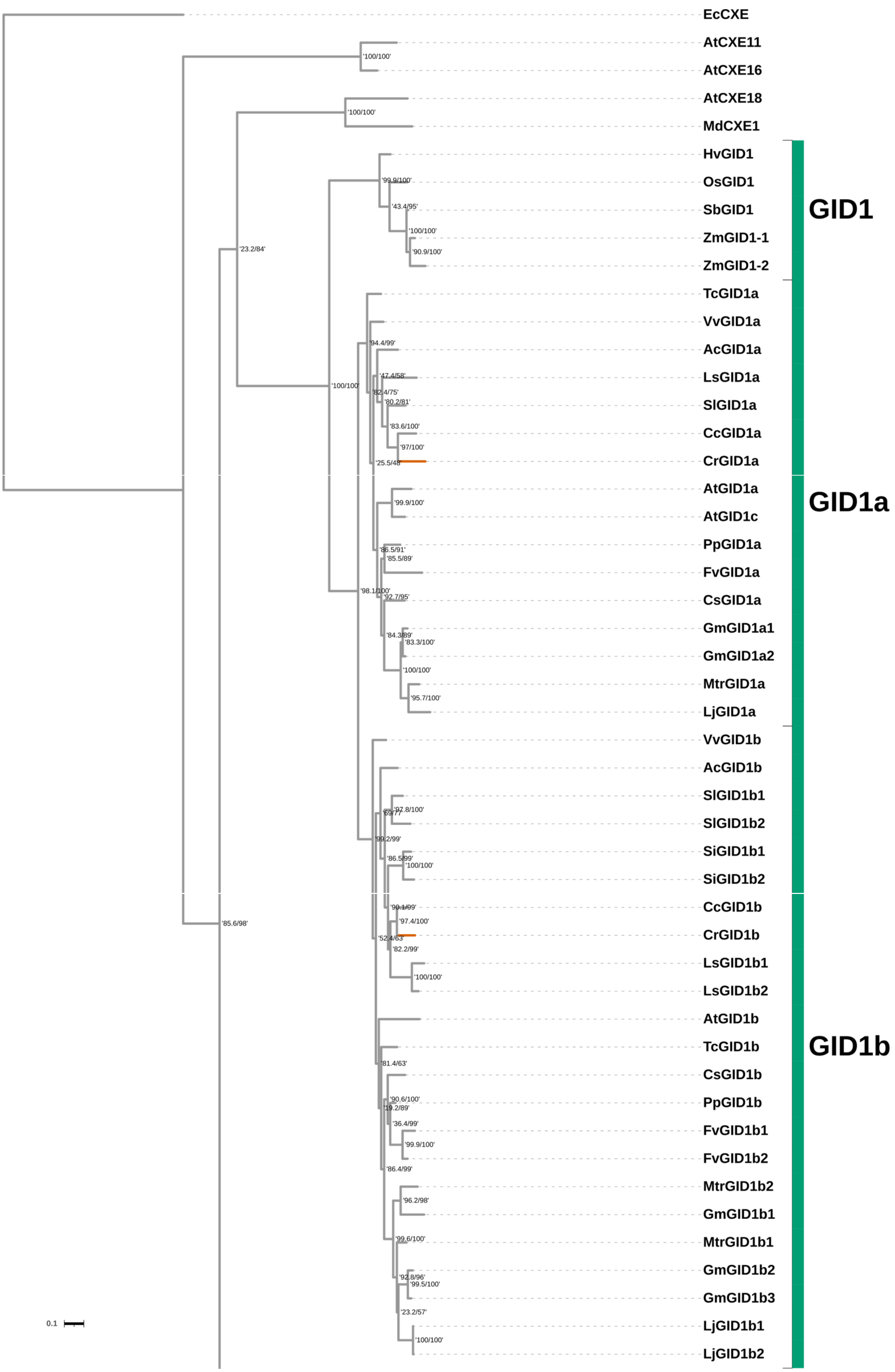


**Figure S2.** Continued on next page…

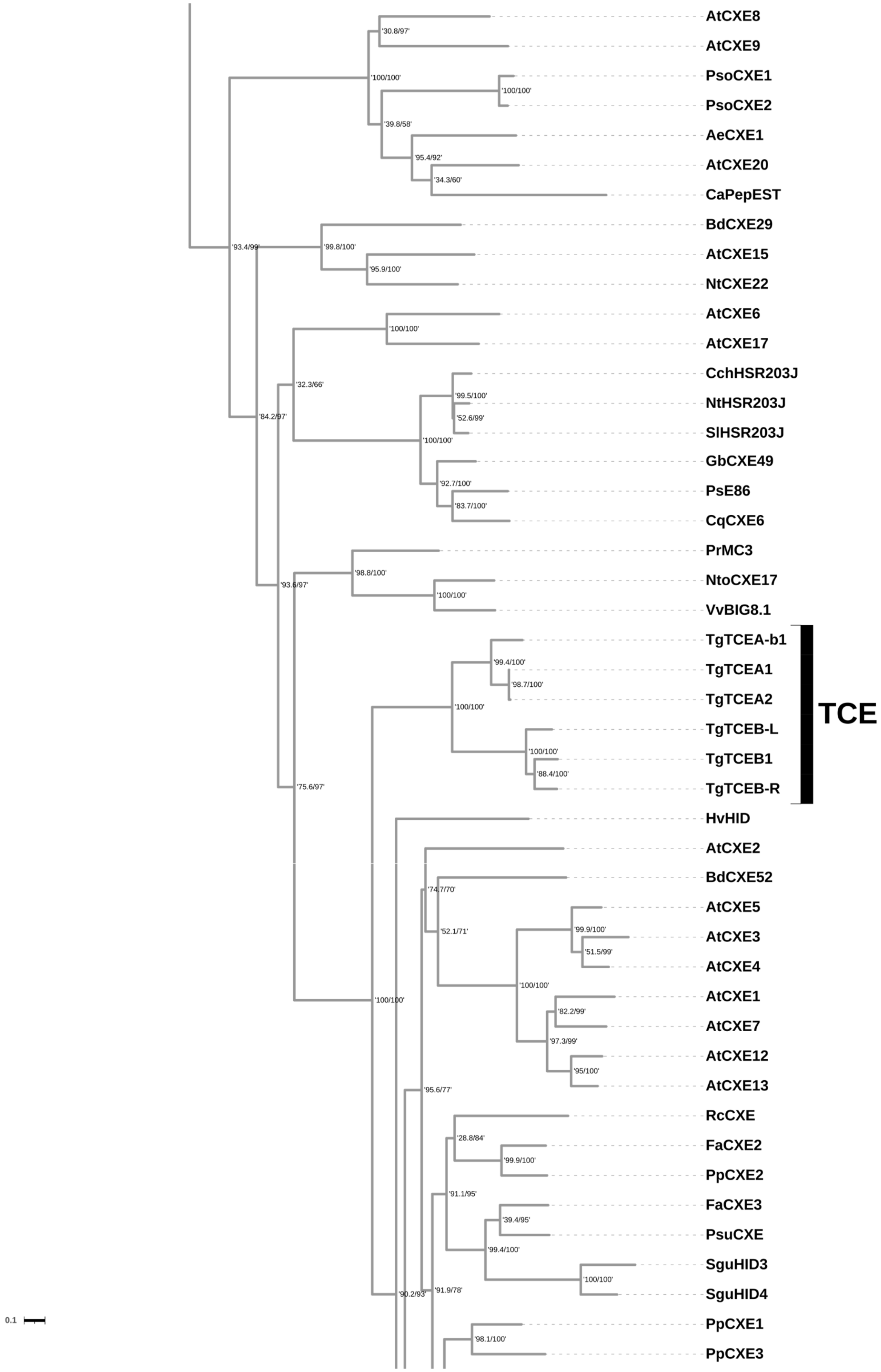


**Figure S2.** Continued on next page…

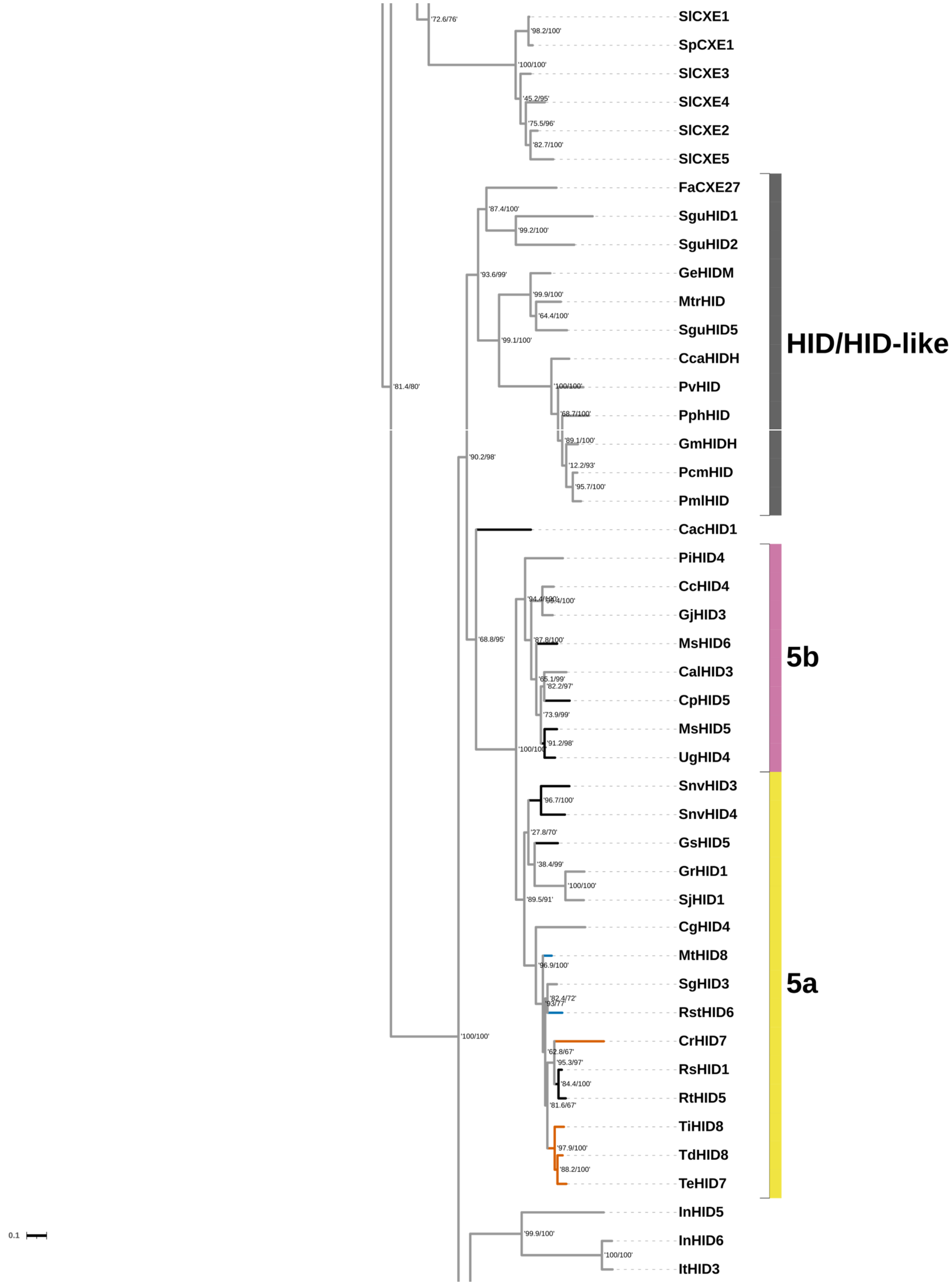


**Figure S2.** Continued on next page…

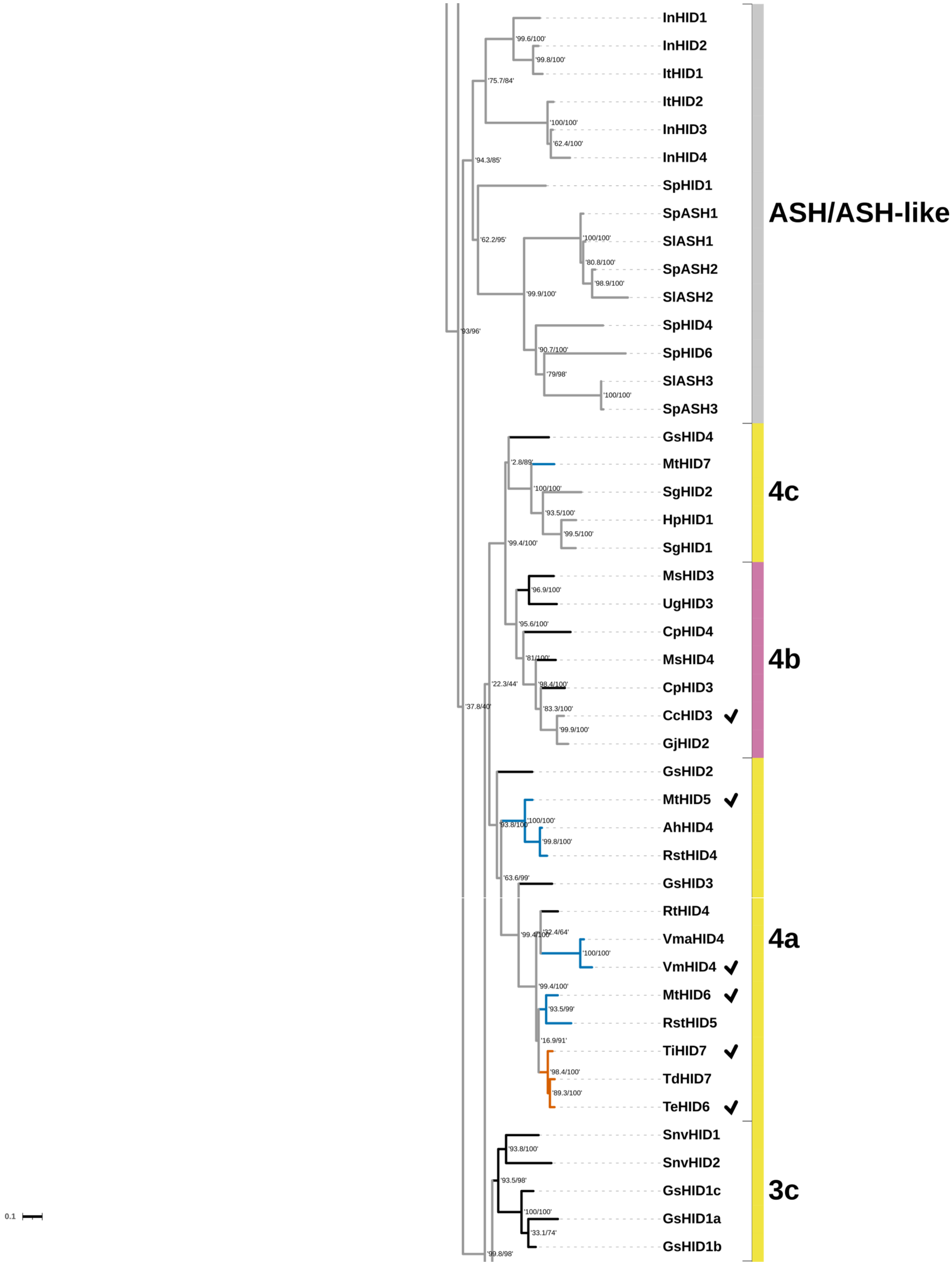


**Figure S2.** Continued on next page…

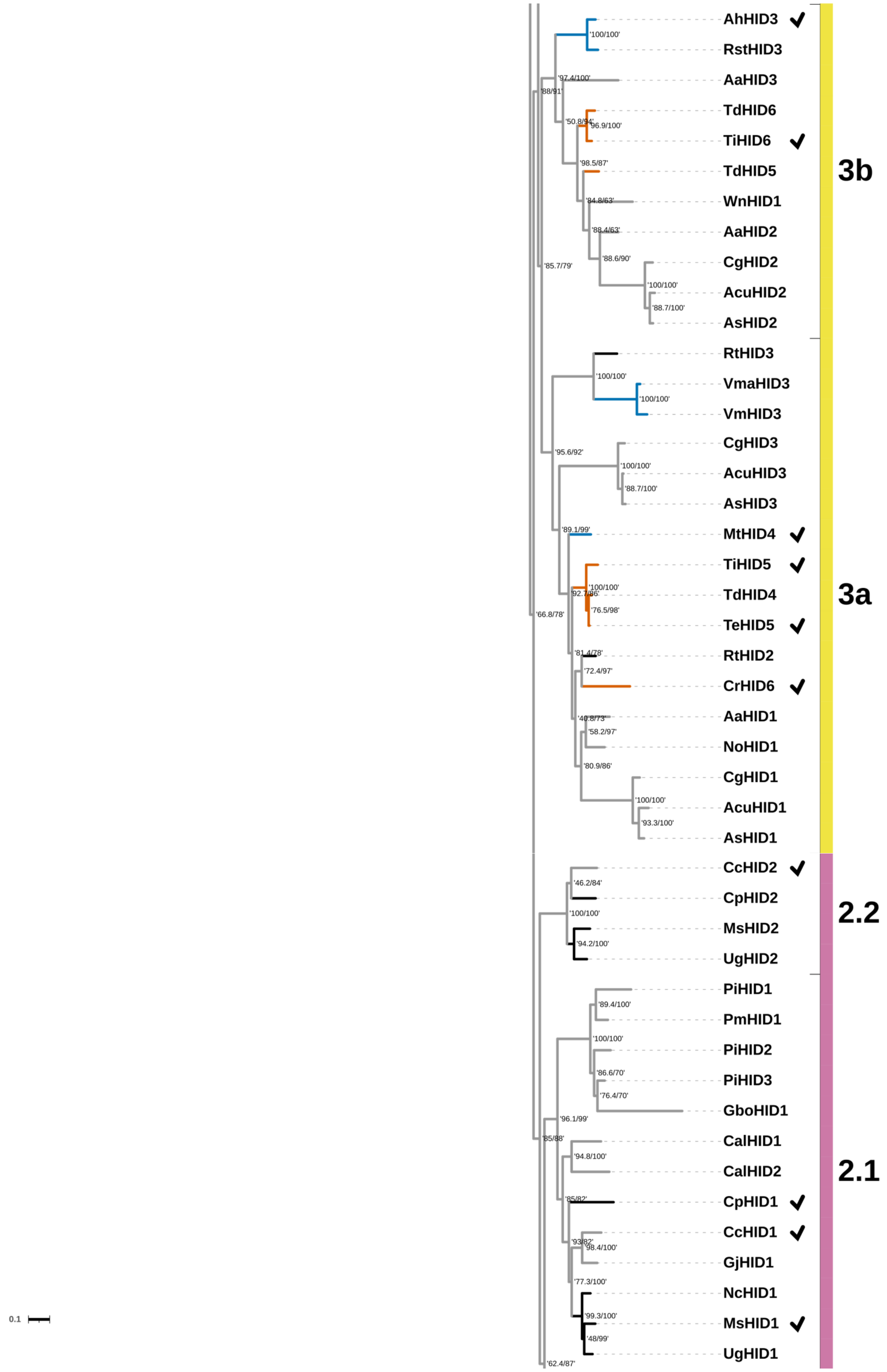


**Figure S2.** Continued on next page…

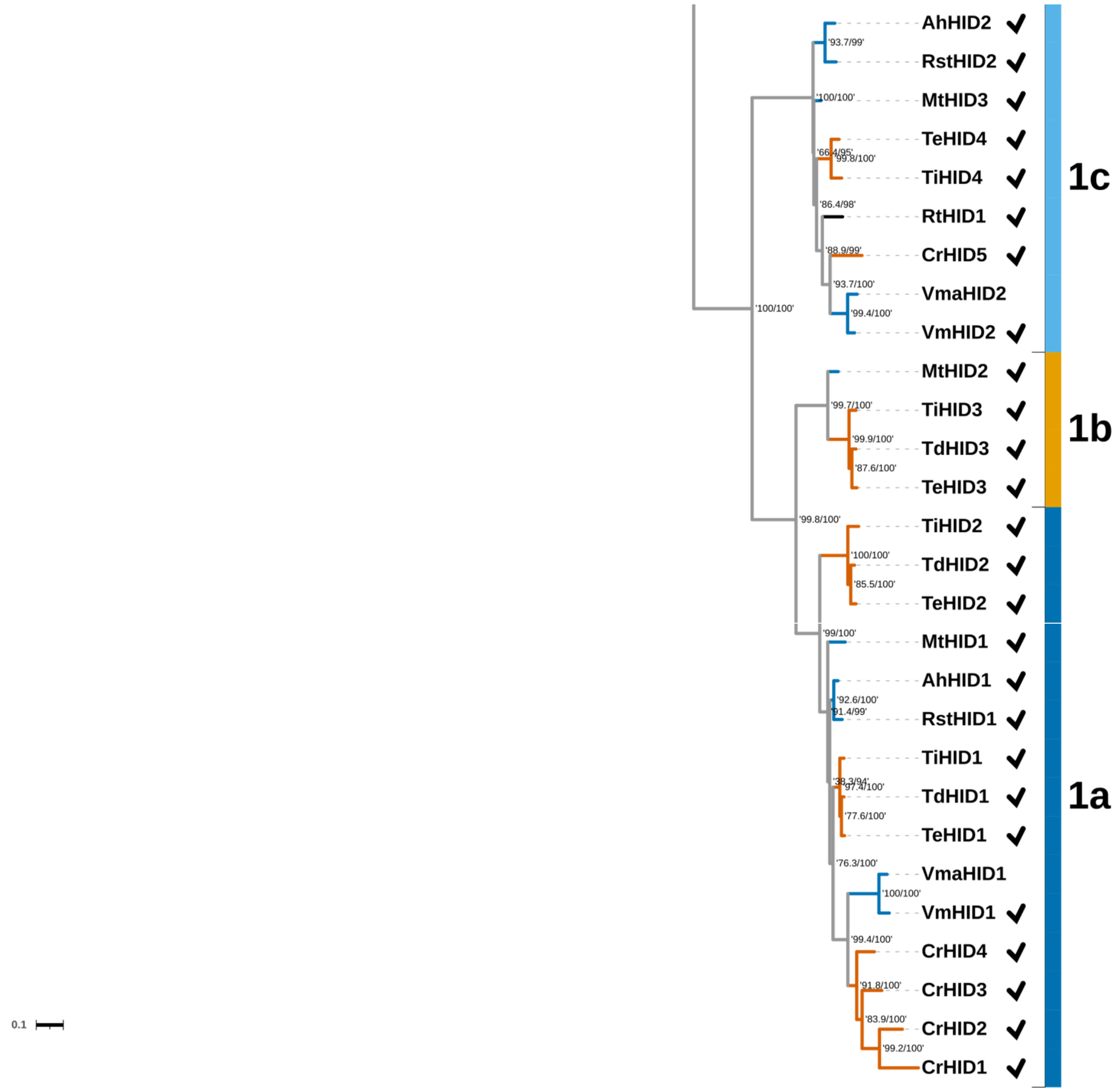


**Figure S2.** Maximum likelihood phylogenetic tree of 258 class I CXE and CXE-like proteins from plants. Values at branch nodes indicate ‘SH-aLRT support (%) / ultrafast bootstrap support (%)’. Clades and branches are labeled/colored as in **Figure 2**. Checkmarks appear next to all enzymes experimentally tested in the present work. Scale bar length represents 0.1 amino acid substitutions per site (see bottom left corner of each page).

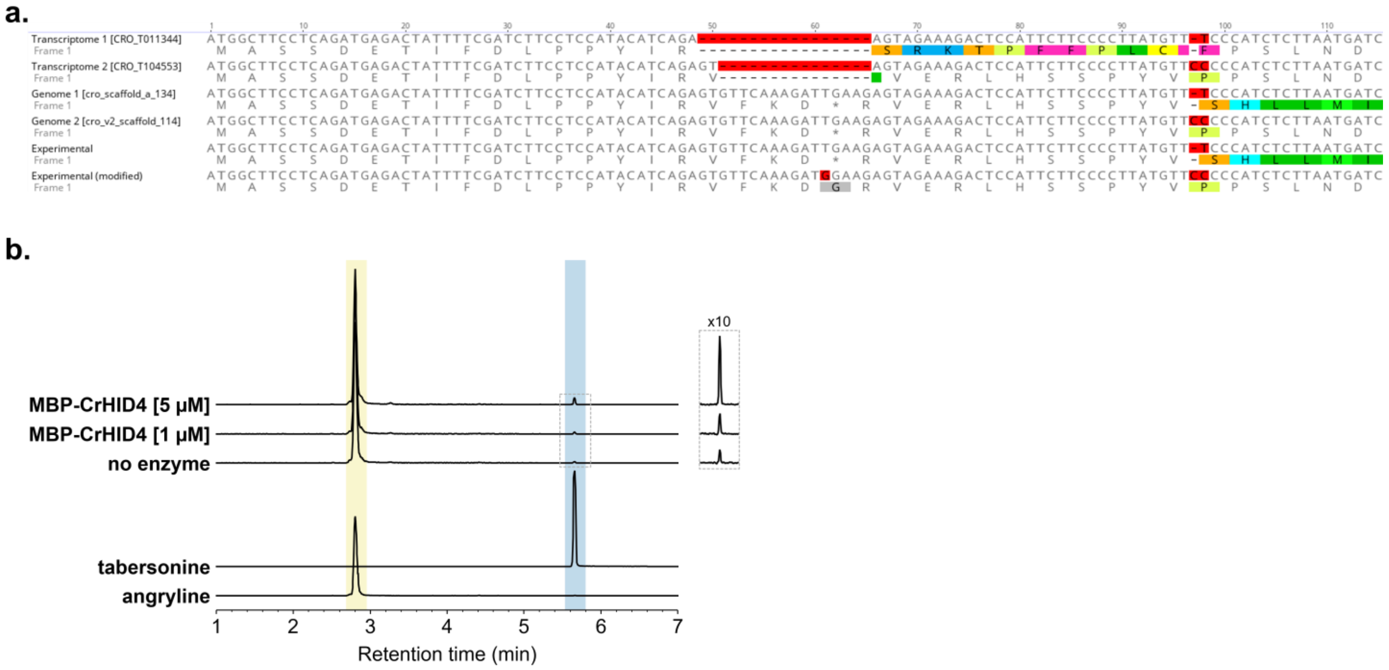


**Figure S3.** Characterization of CrHID4. (**a**) Alignment of sequences corresponding to CrHID4. The predicted sequence from transcriptome 1 (CRO_T011344) is identical to HL4 (MF770515.1) previously reported.[20,21] The experimental sequence obtained in this work is identical to the predicted sequence from genome 1 (cro_scaffold_a_134). As this sequence contains an internal stop codon (TGA at position 61 of the alignment) and a deletion resulting in a frameshift (position 97), the experimental sequence was modified to remove the stop codon (TGA → GGA; this codon is GGA and/or codes for Gly in all extant cyclases) and insert the missing nucleotide (C at position 97 according to the transcriptome 2 and genome 2 sequences). All nucleotide sequences from position 99 to the end are identical. (**b**) LC-MS traces (EIC 337.1911 ± 0.01) of reactions between angryline and MBP-tagged CrHID4 at 1 µM or 5 µM enzyme. Overexpression of the modified experimental CrHID4 gene sequence yields very low levels of enzyme. Attaching an MBP tag at the N-terminus improves expression, but the enzyme exhibits very low TS activity.

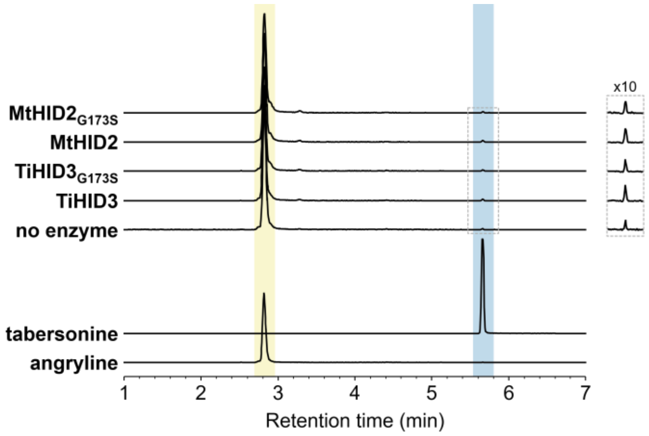


**Figure S4.** LC-MS traces (EIC 337.1911 ± 0.01) of reactions between angryline and TiHID3, TiHID3$_{G173S}$, MtHID2, and MtHID2$_{G173S}$ (all at 5 µM enzyme). Tabersonine is not generated above background levels in these reactions.

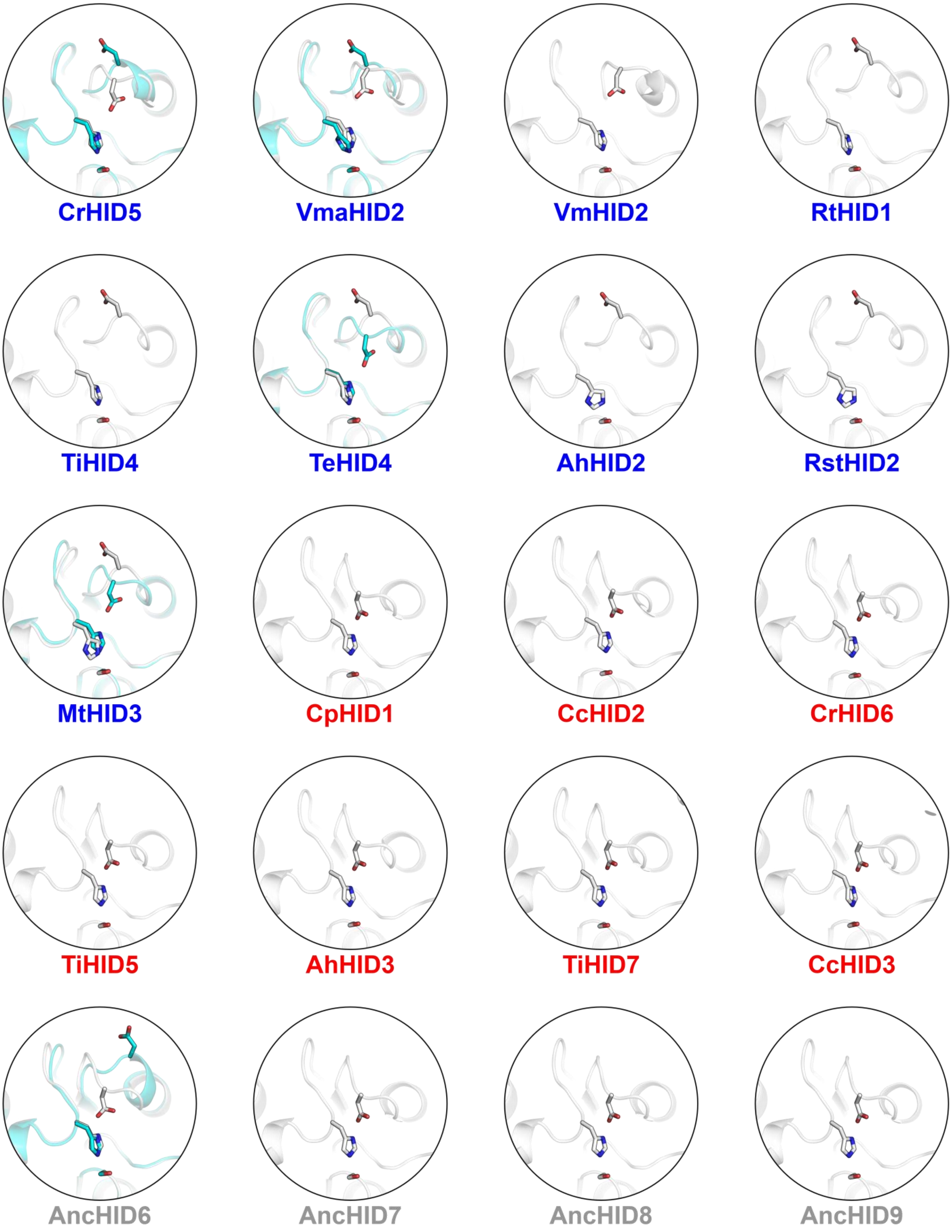


**Figure S5.** AlphaFold models of Clade 1c CXE-like proteins (labeled in blue), representative extant CXEs (labeled in red), and ancestral enzymes AncHID6–AncHID9 (labeled in gray) with catalytic triad residues (Ser-His-Asp) shown as sticks. For four of the CXE-like proteins and AncHID6, the side chain of the catalytic Asp is positioned either inward near the catalytic His or outside of the active site. Top-ranking models (rank 1) are depicted in white whereas lower-ranking models showing an alternative position of the catalytic Asp (rank 2, CrHID5; rank 3, VmaHID2; rank 5, TeHID4; rank 4, MtHID3; rank 2, AncHID6) are depicted in cyan. Note that VmHID2 is the only Clade 1c CXE-like protein for which all five AlphaFold models (rank 1–rank 5) predict the catalytic Asp to be positioned inward near the catalytic His. Two, four, one, one, and one model(s) predict this to be the case for CrHID5, VmaHID2, TeHID4, MtHID3, and AncHID6, respectively. For the remaining four CXE-like proteins, all five AlphaFold models predict the catalytic Asp to be positioned outside of the active site. This residue is predicted to be positioned inward near the catalytic His in all models of the extant as well as ancestral (AncHID7–AncHID9) CXEs. All AlphaFold models shown here and in subsequent figures were predicted using ColabFold v1.5.2.[22]

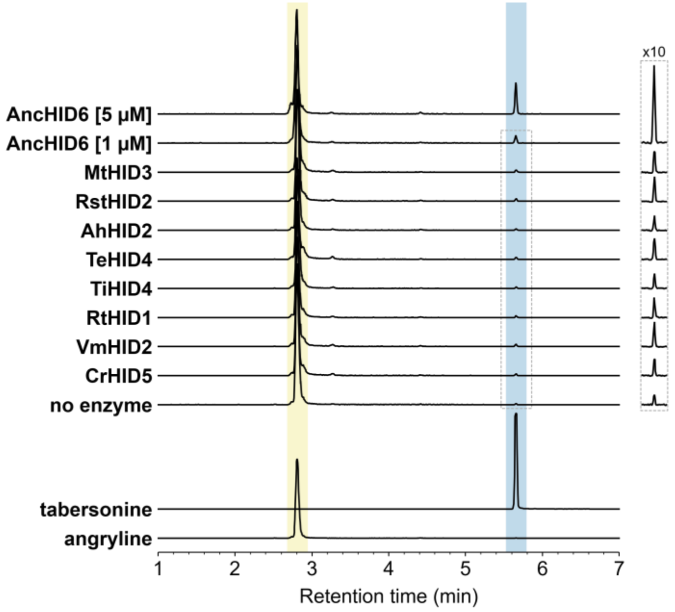


**Figure S6.** LC-MS traces (EIC 337.1911 ± 0.01) of reactions between angryline and Clade 1c enzymes (all at 5 μM except for AhHID2 (2.5 μM)) as well as AncHID6 (at 1 μM or 5 μM as indicated). Among these enzymes, only AncHID6 is capable of generating tabersonine at low levels.

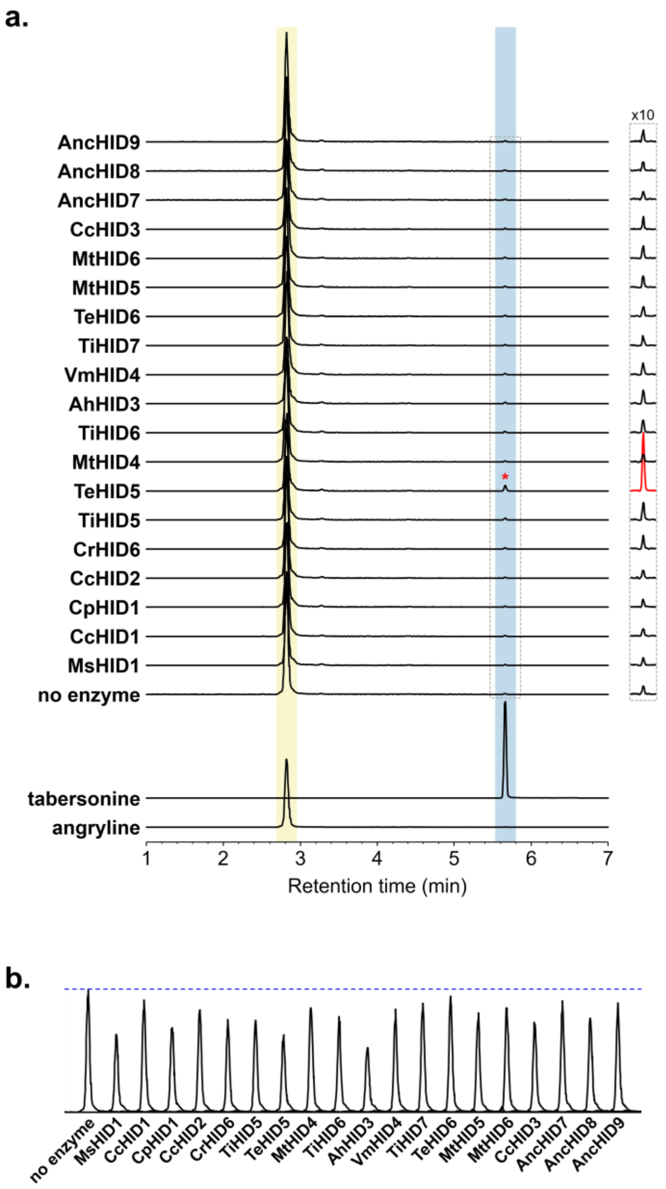


**Figure S7.** LC-MS traces (EIC 337.1911 ± 0.01) of reactions between angryline and 16 extant CXEs from Clades 2–4 (all at 5 μM except for CcHID2, TeHID6, and MtHID6 (2.5 μM)) as well as AncHID7, AncHID8, and AncHID9 (all at 5 μM). (**a**) Tabersonine is not generated above background levels in these reactions. The low level of tabersonine seen in the reaction with TeHID5 (red asterisk) is likely due to a small amount of contaminating TS, as this result was not reproducible. (**b**) Peaks (EIC 337.1911 ± 0.01) corresponding to remaining starting material (angryline) in each reaction. Moderate levels of angryline depletion are observed in reactions with some CXEs, suggesting accelerated angryline degradation in the presence of these enzymes.

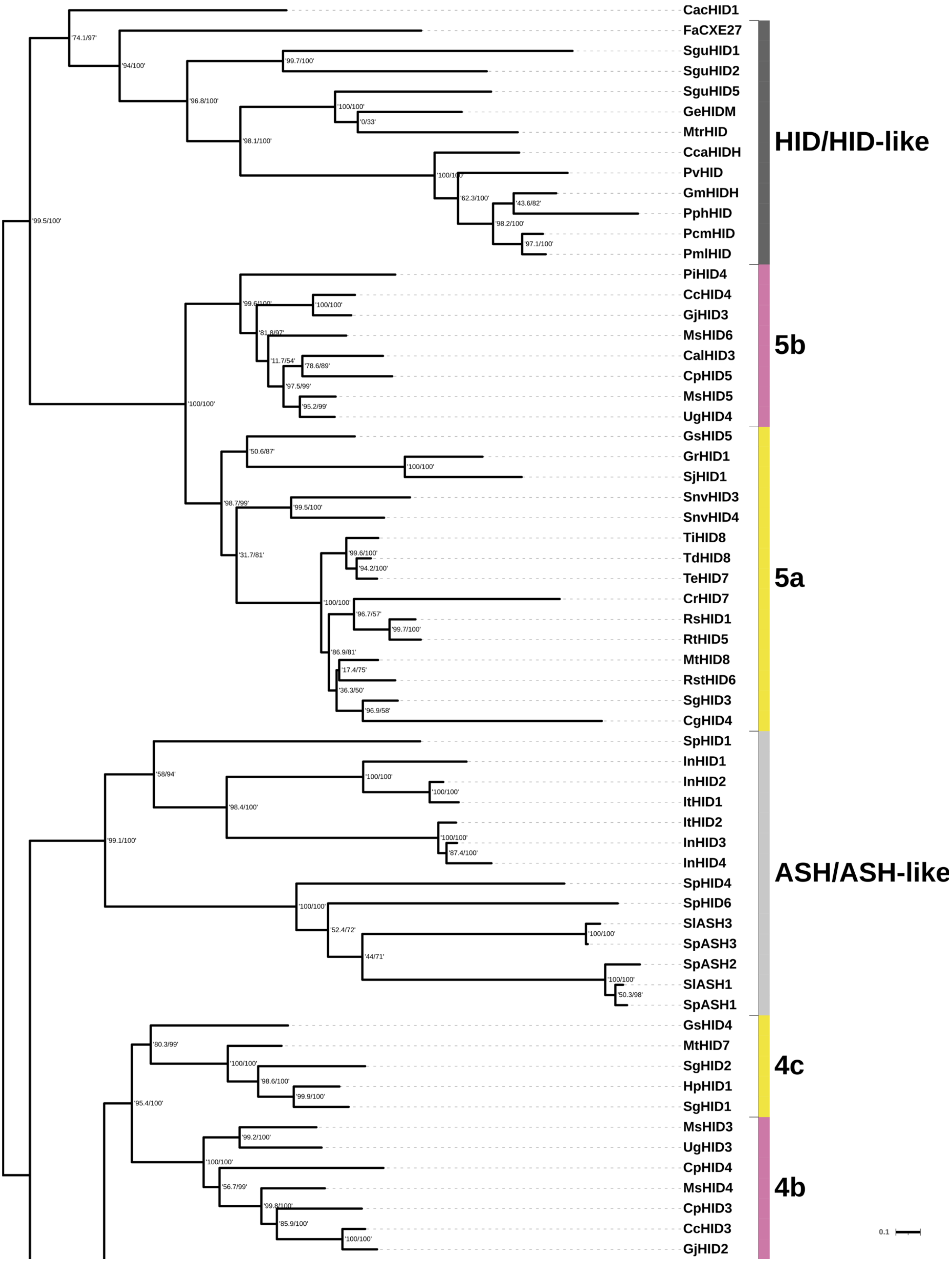


**Figure S8.** Continued on next page…

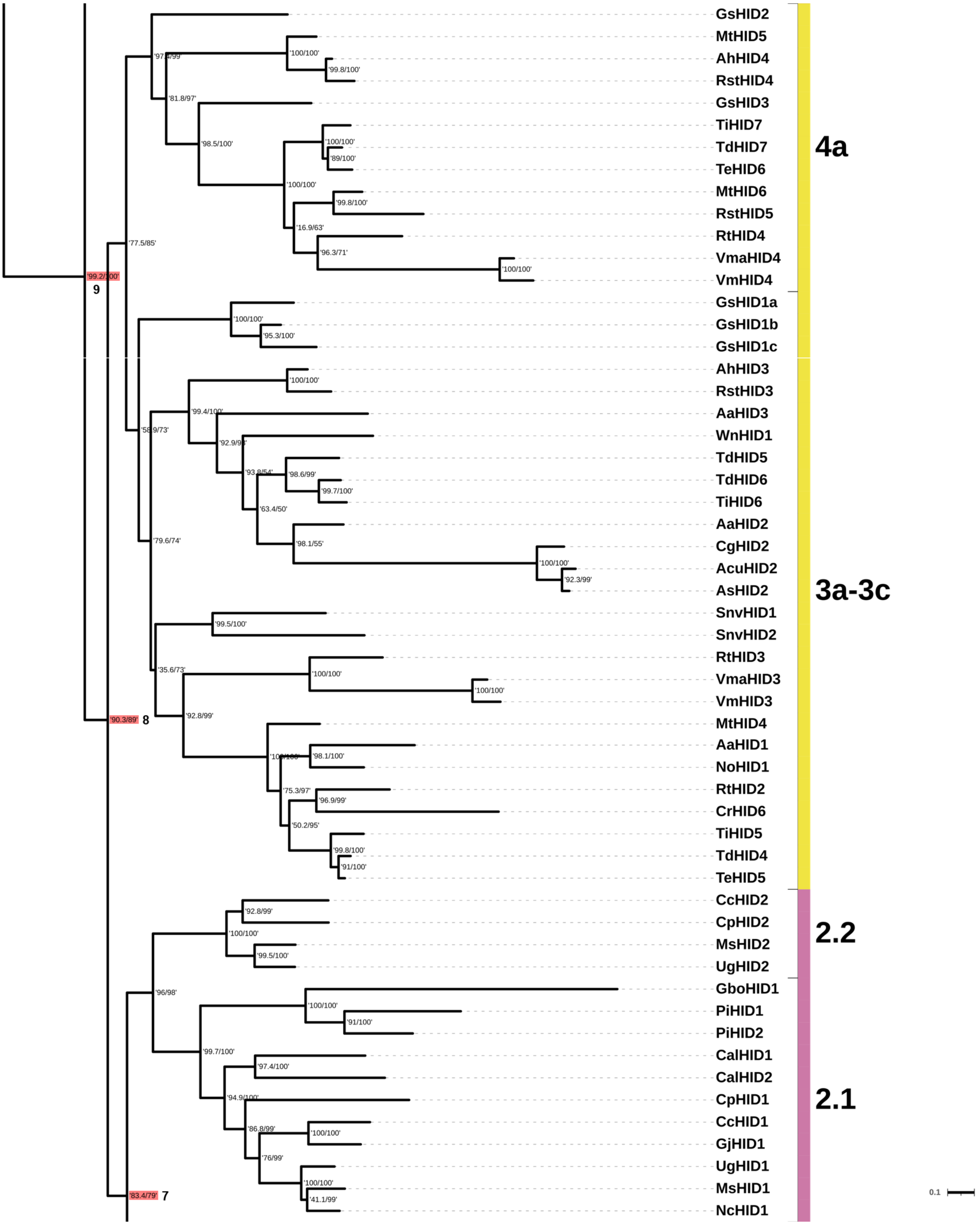


**Figure S8.** Continued on next page…

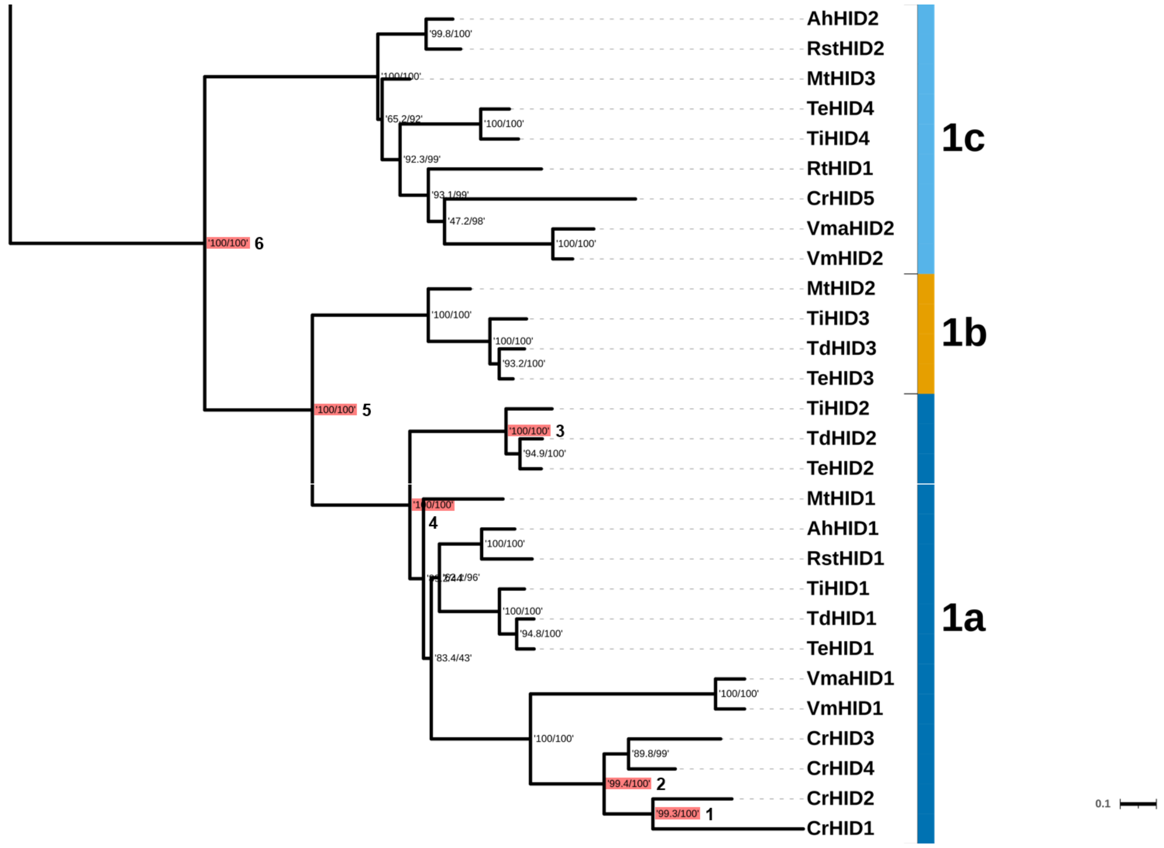


**Figure S8.** Maximum likelihood phylogenetic tree of 145 class I CXE and CXE-like proteins from plants. This more focused phylogeny was used for ancestral sequence reconstruction analysis. Values at branch nodes indicate 'SH-aLRT support (%) / ultrafast bootstrap support (%)'. Clades are labeled and colored as in **Figures 2** and **4**. Reconstructed ancestral nodes are highlighted in red and labeled 1–9. Scale bar length represents 0.1 nucleotide substitutions per codon site (see bottom right corner of each page).

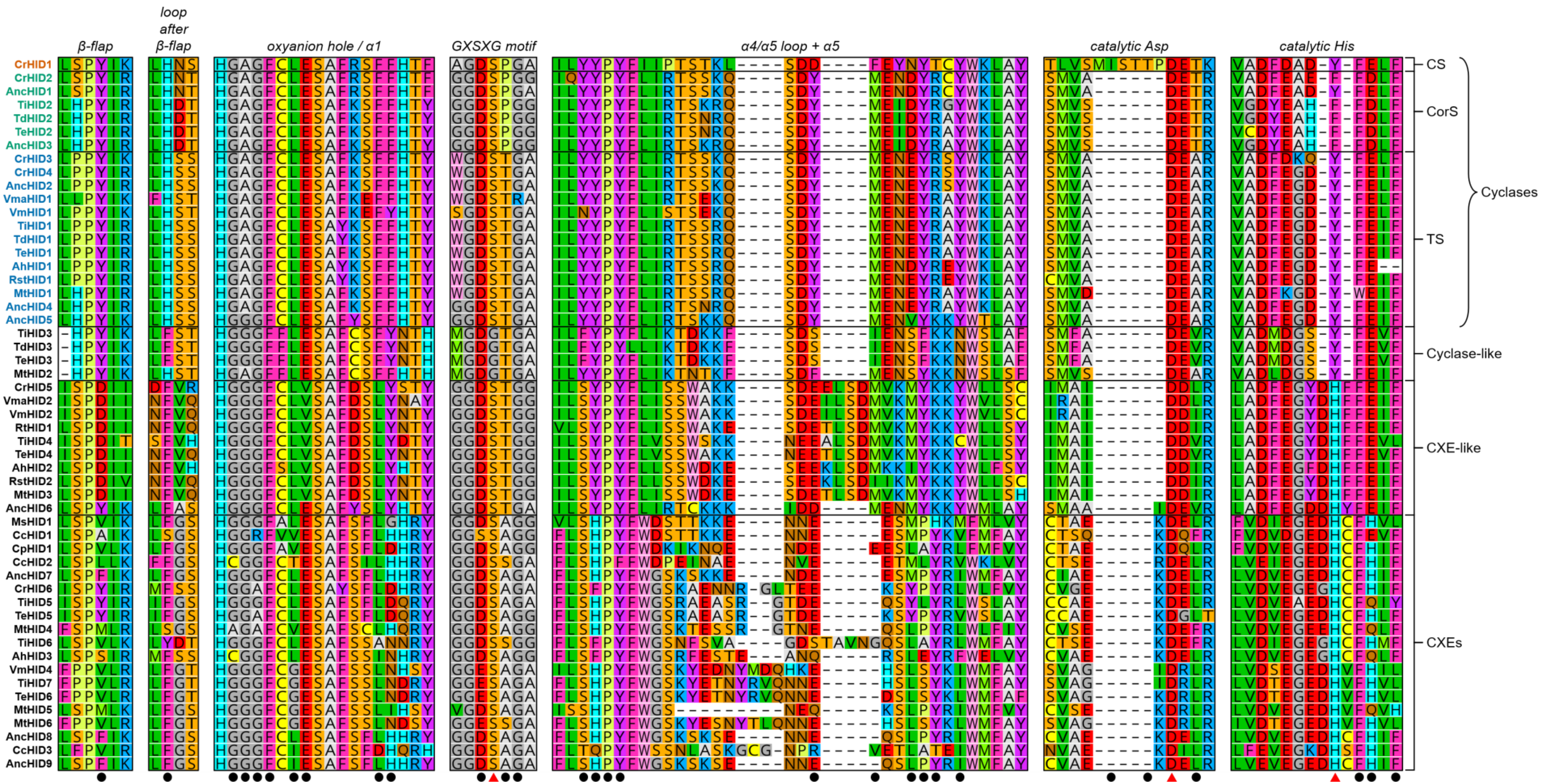


**Figure S9.** Multiple sequence alignment (MSA) of selected extant and ancestral HID-like enzymes. Seven contiguous regions encompass all amino acid residues located within 6 Å of the substrate binding pocket (black circles). Red triangles are located below the three residues making up the catalytic triad. The names of the individual cyclases are colored according to the identity of their major cyclization product (red = catharanthine, green = 16-cmc, blue = tabersonine). All proteins from Clade 1, all experimentally tested CXEs from Clades 2–4, and all AncHIDs are shown. Except for VmaHID1 and VmaHID2, all proteins included in the alignment were experimentally tested.

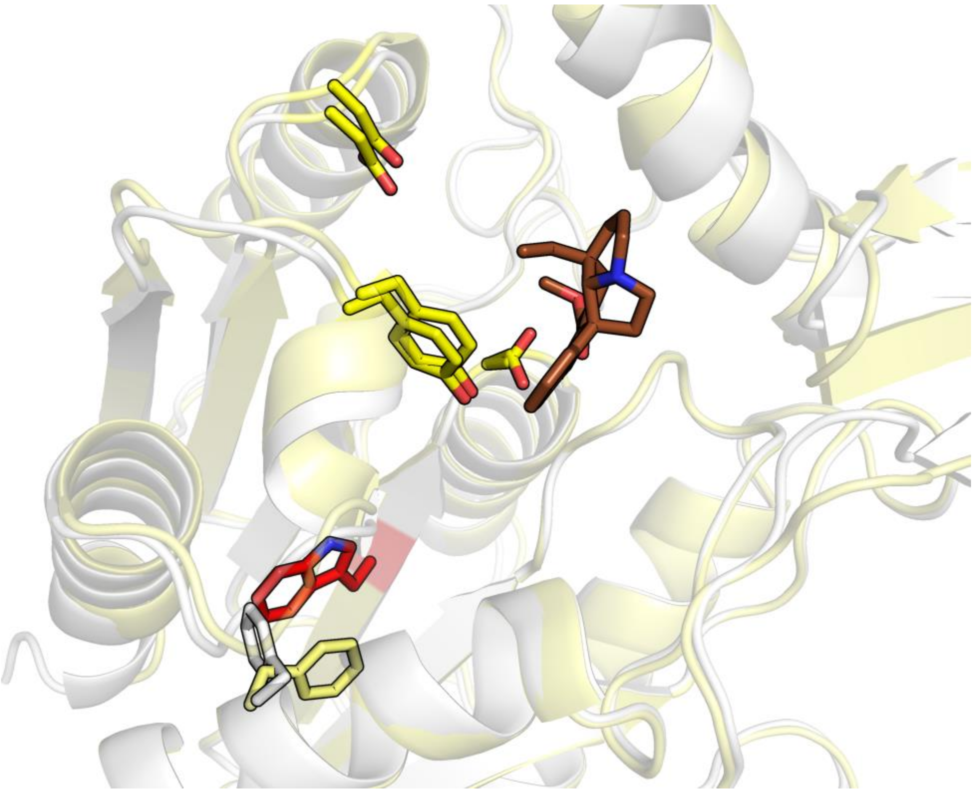

**Figure S10.** Superposition of CrHID3 (experimental structure, **PDB 6RS4**) and AncHID4 (AlphaFold model[22]). The protein backbone chain is shown in light yellow (CrHID3) or white (AncHID4), docked product (tabersonine) is shown in brown, catalytic triad residues are shown in yellow, and $CrHID3_{W167}$/$AncHID4_{G171}$ are shown in red. Although the latter two residues occupy an equivalent position outside of the substrate binding pocket, the $AncHID4_{G171W}$ mutant exhibits higher activity than the wild-type ancestor. The newly introduced Trp residue may impact protein dynamics by interacting with alpha helix 1 via pi stacking with the adjacent Phe ($CrHID3_{F99}$ shown in light yellow; $AncHID4_{F103}$ shown in white).

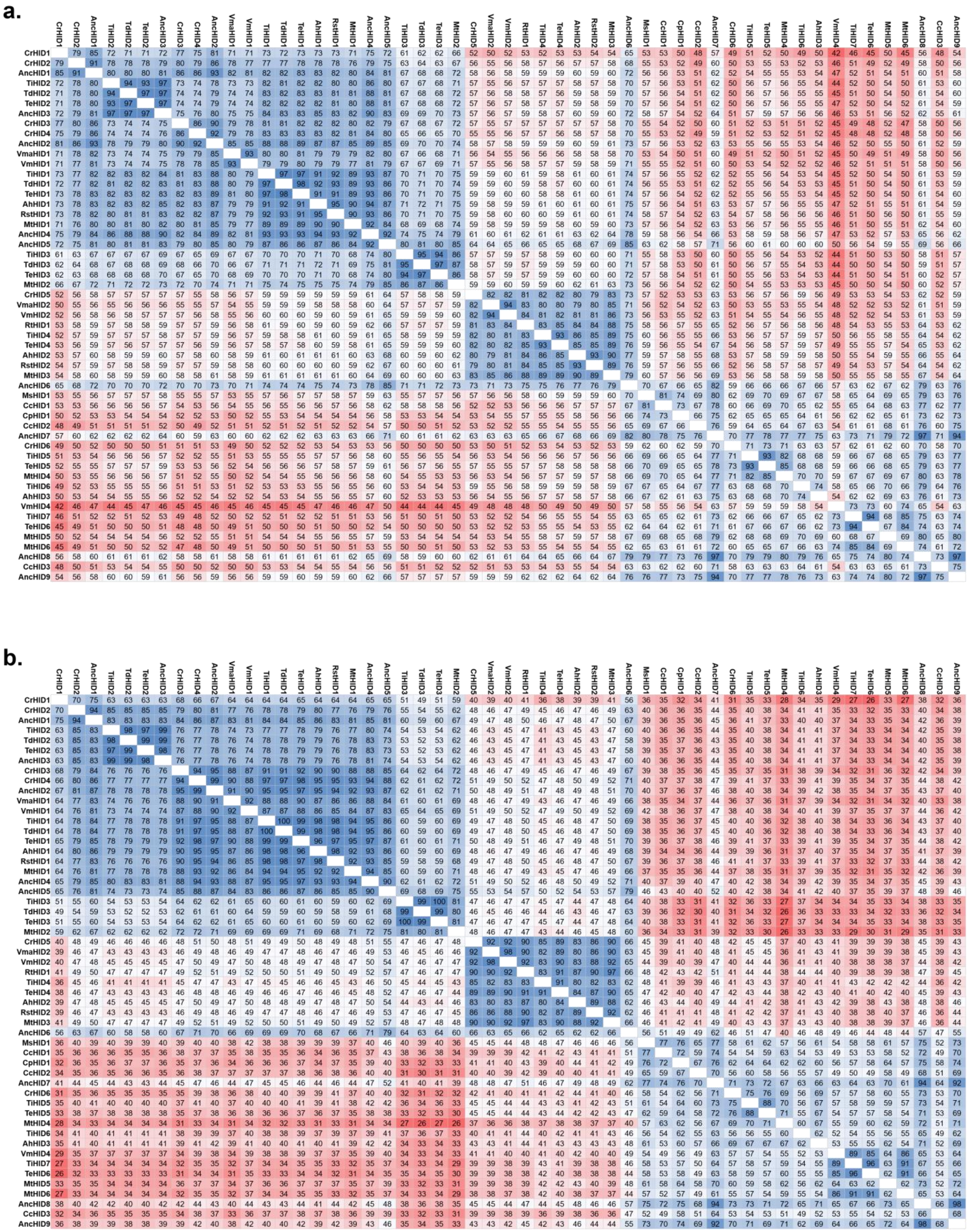


**Figure S11.** Sequence identities of selected extant and ancestral HID-like enzymes. (**a**) Sequence identities (% ID) based on an alignment of full protein sequences. (**b**) Sequence identities (% ID) based on an alignment of just the seven key regions shown in **Figure S9**.

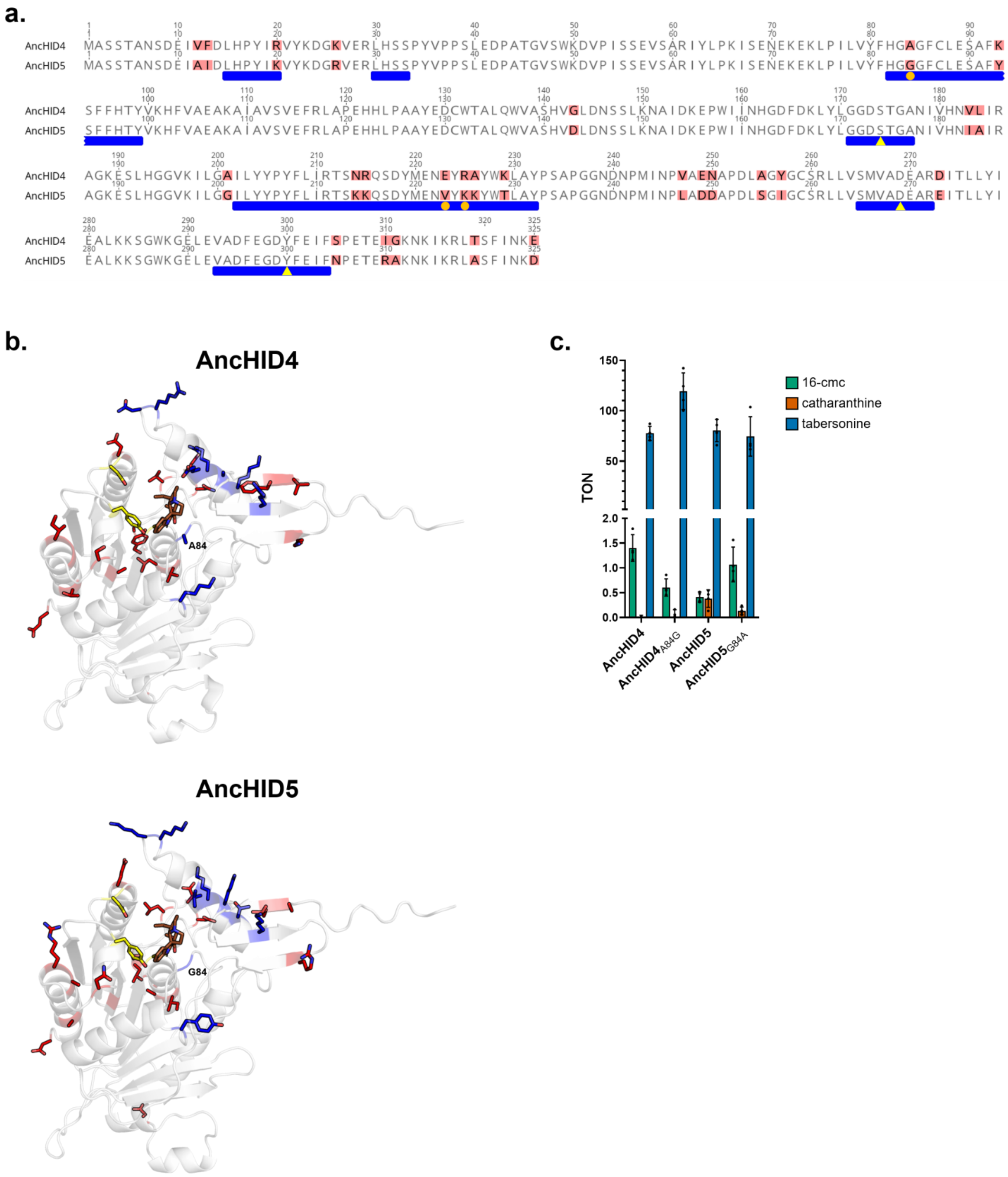


**Figure S12.** (**a**) Pairwise sequence alignment of AncHID4 and AncHID5. All differences from the consensus sequence are highlighted in red. Blue bars appear below the seven contiguous regions encompassing all amino acid residues located in proximity to the substrate binding pocket. Orange circles indicate differences in residues located within 6 Å of bound substrate/product. Yellow triangles indicate the residues comprising the catalytic triad. (**b**) Comparison of AncHID4 and AncHID5 (both AlphaFold models[22]). Docked product (tabersonine) is shown in brown, catalytic triad residues are shown in yellow, and the 27 residues differing between the two cyclases are shown in blue and red. Residues highlighted in blue (x9) represent all differences in the seven key regions surrounding the substrate binding pocket. Residue 84 is labeled in each model. (**c**) Angryline cyclization activity of wild-type and mutant cyclases expressed as turnover number (TON = mol product/mol enzyme). Colored bar height indicates the mean of four independent experiments. The results of individual experiments are represented by dots (error bars = SD). Reactions were performed at 37 °C for 30 min in 50 mM Tris (pH 9.0) with 50 μM angryline and 200 nM enzyme.

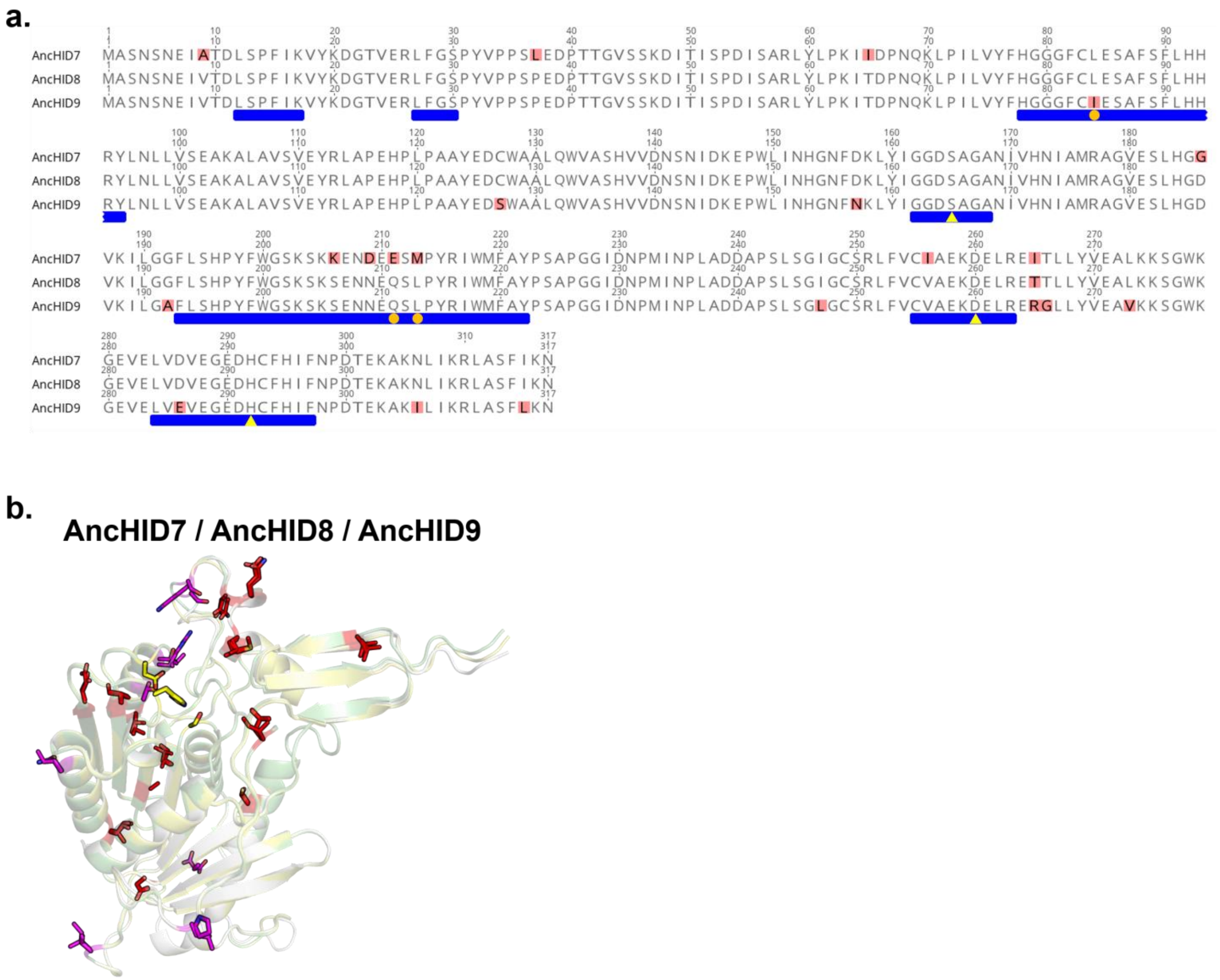


**Figure S13.** (**a**) Multiple sequence alignment of AncHID7, AncHID8, and AncHID9. All differences from the consensus sequence are highlighted in red. Blue bars appear below the seven contiguous regions encompassing all amino acid residues located in proximity to the substrate binding pocket. Orange circles indicate differences in residues located within 6 Å of bound substrate/product. Yellow triangles indicate the residues comprising the catalytic triad. (**b**) Superposition of AlphaFold models[22] of AncHID7 (white), AncHID8 (pale yellow), and AncHID9 (pale green). Catalytic triad residues are shown in yellow, and the 20 residues differing between the three enzymes are shown in red/magenta. Residues highlighted in magenta (x7) represent all non-conservative differences, none of which are located in the substrate binding pocket. Conservative differences (highlighted in red) include: A/G, C/S, D/E, D/N, E/Q, L/I, L/M, V/A, V/I, V/L.

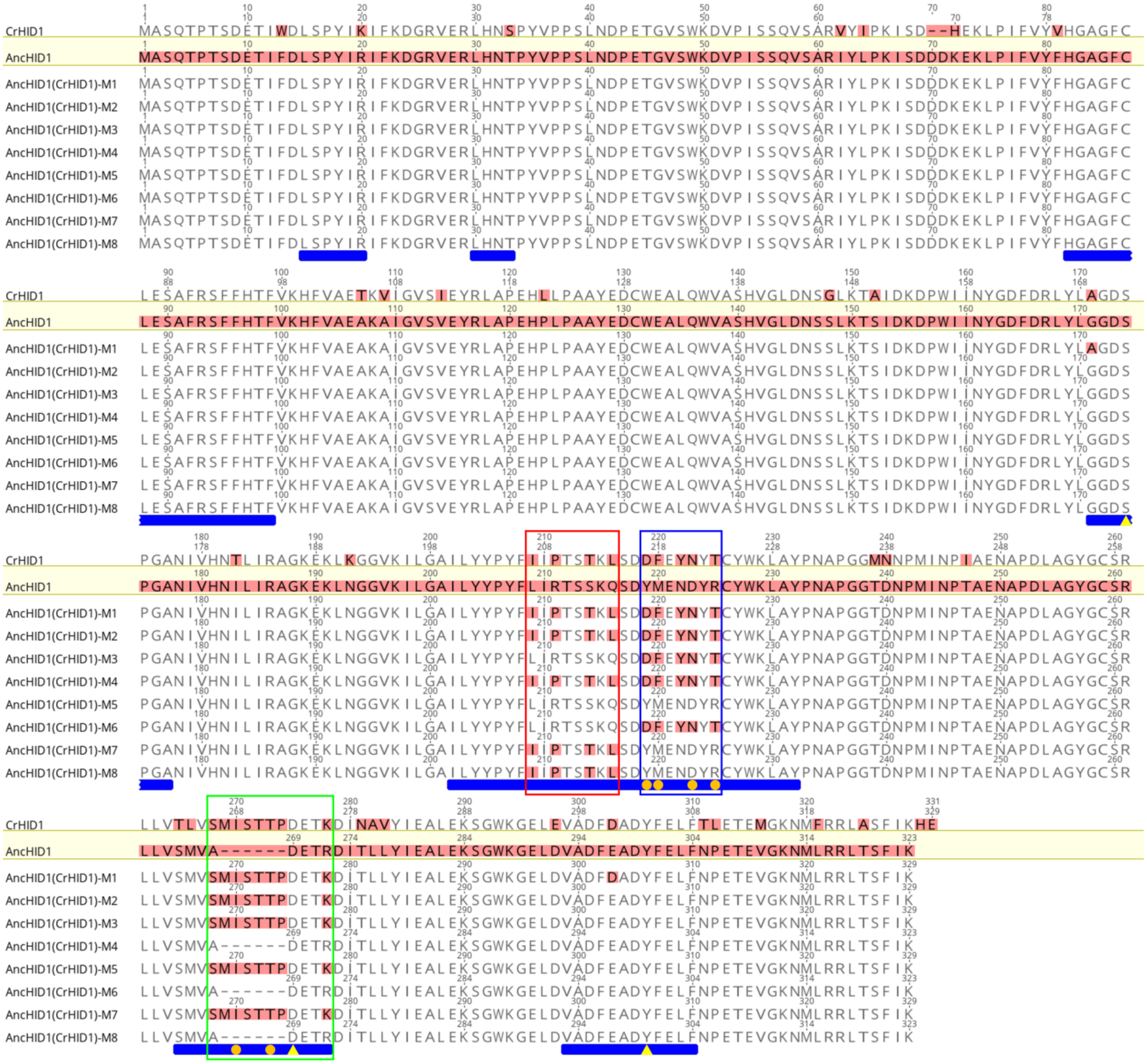


**Figure S14.** Multiple sequence alignment of CrHID1, AncHID1, and AncHID1 mutants. The template sequence (AncHID1) is enclosed in a yellow box, and all amino acid residues are highlighted in red. Among the other sequences, all differences from the template are highlighted in red. Blue bars appear below the seven contiguous regions encompassing all amino acid residues located in proximity to the substrate binding pocket. Orange circles indicate differences in residues located within 6 Å of bound substrate/product. Yellow triangles indicate the residues comprising the catalytic triad. Three regions individually probed in mutagenesis experiments are boxed in red (α4/α5 loop), blue (α5), and green (extended loop/catalytic Asp). See **Figure S15** for a comparison of the template (AncHID1) and target (CrHID1) structures as well as activity assay results.

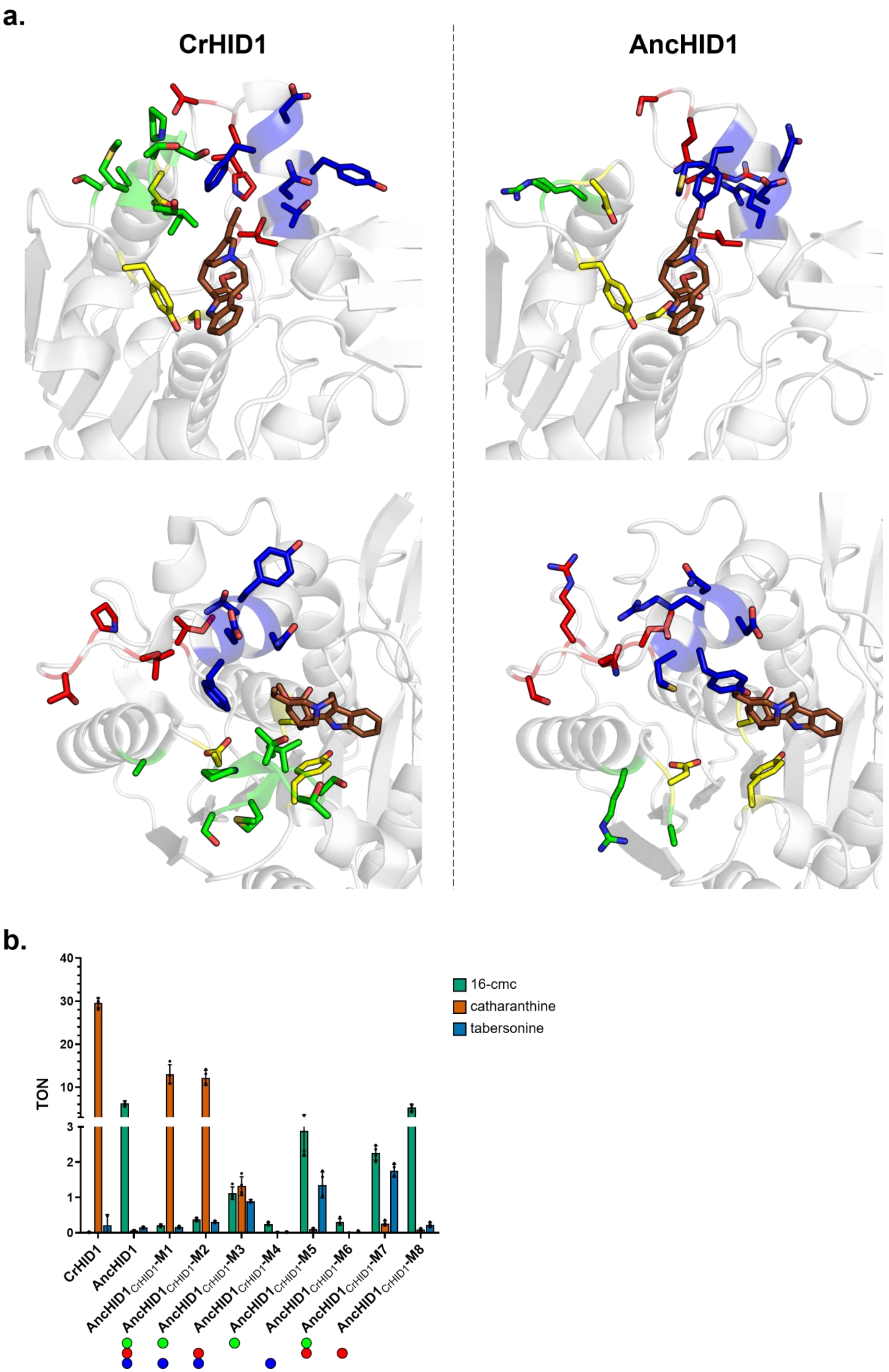


**Figure S15.** See next page for caption.

**Figure S15.** (**a**) Comparison of CrHID1 (experimental structure, **PDB 6RT8**) and AncHID1 (AlphaFold model[22]). Bound intermediate/docked product (16-cmc) is shown in brown, catalytic triad residues are shown in yellow, and residues located in the α4/α5 loop, α5, and extended loop/catalytic Asp regions that differ between the two cyclases are shown in red (α4/α5 loop), blue (α5), and green (extended loop/catalytic Asp). Two different views are shown for each structure. (**b**) Angryline cyclization activity of wild-type and mutant cyclases expressed as turnover number (TON = mol product/mol enzyme). Colored bar height indicates the mean of four independent experiments. The results of individual experiments are represented by dots (error bars = SD). Reactions were performed at 37 °C for 30 min in 50 mM Tris (pH 9.0) with 50 µM angryline and 1 µM enzyme. To facilitate interpretation of the results, colored circles corresponding to regions (see part **a** for color designations) replaced in each AncHID1$_{CrHID1}$ mutant appear below the mutant names (M2–M8) in the bar chart.

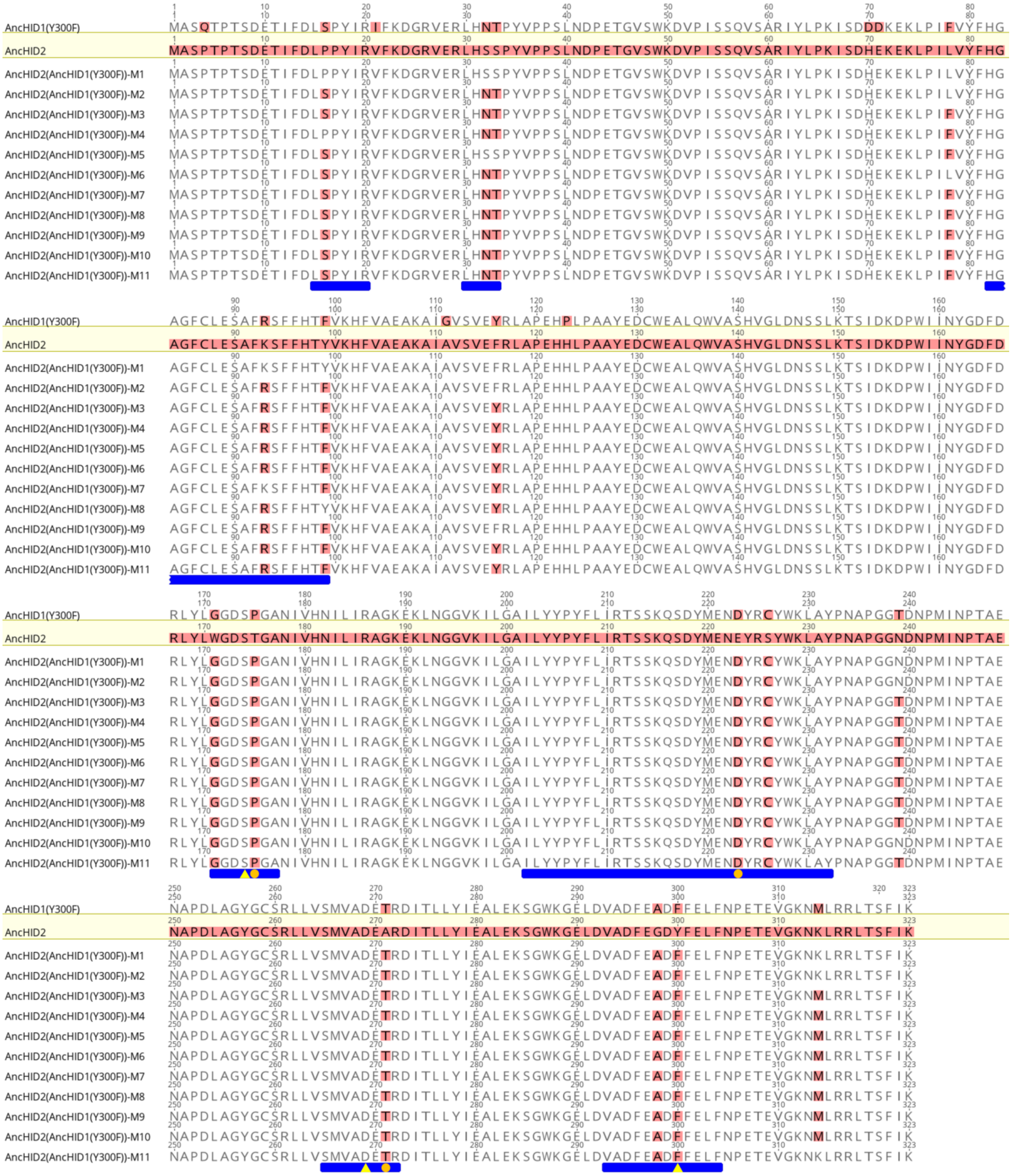


**Figure S16.** Multiple sequence alignment of AncHID1$_{Y300F}$, AncHID2, and AncHID2 mutants. The template sequence (AncHID2) is enclosed in a yellow box, and all amino acid residues are highlighted in red. Among the other sequences, all differences from the template are highlighted in red. Blue bars appear below the seven contiguous regions encompassing all amino acid residues located in proximity to the substrate binding pocket. Orange circles indicate differences in residues located within 6 Å of bound substrate/product. Yellow triangles indicate the residues comprising the catalytic triad. See **Figure S17** for a comparison of the template (AncHID2) and target (AncHID1$_{Y300F}$) structures as well as activity assay results.

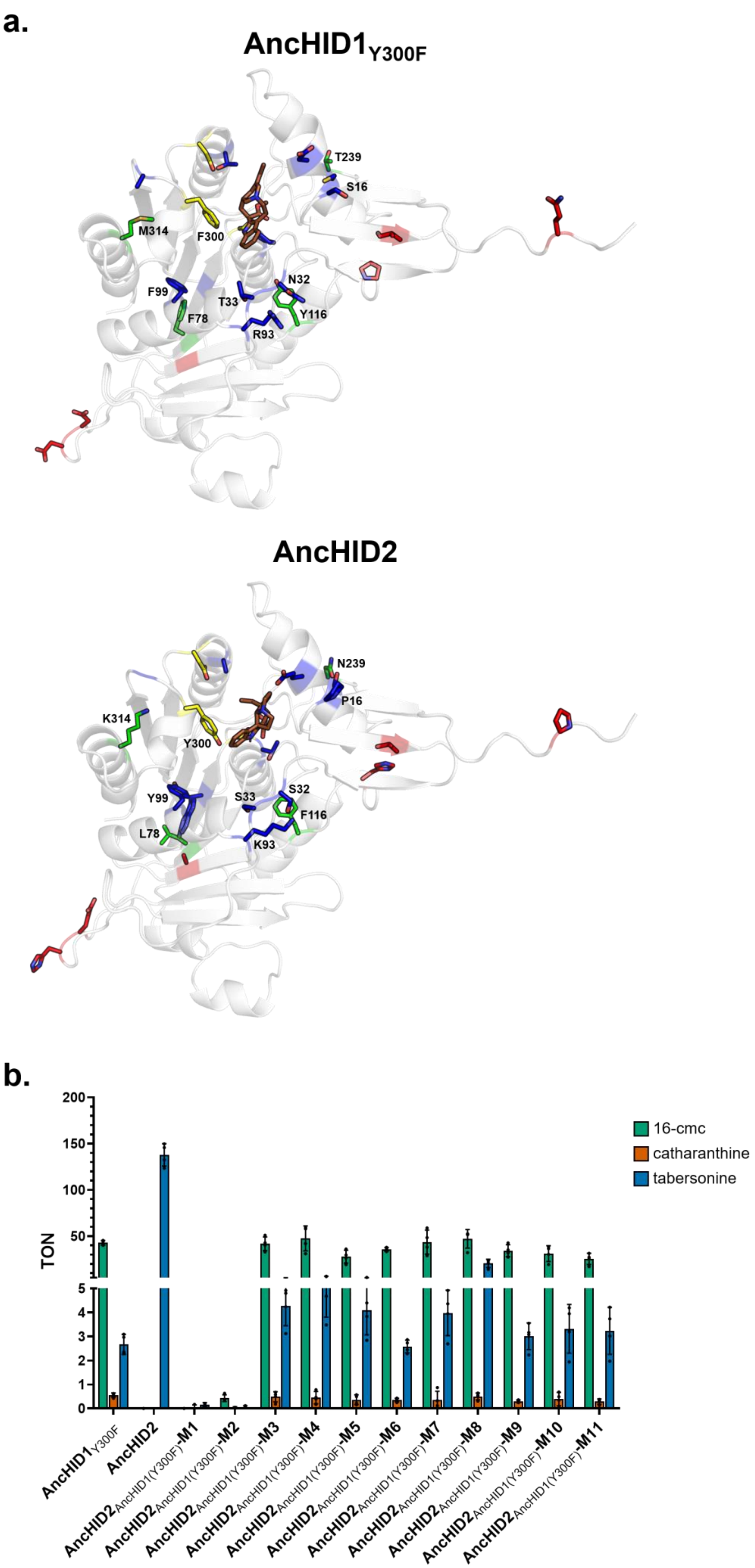


**Figure S17.** See next page for caption.

**Figure S17.** (**a**) Comparison of AncHID1$_{Y300F}$ and AncHID2 (both AlphaFold models[22]). Docked product (16-cmc for AncHID1$_{Y300F}$, tabersonine for AncHID2) is shown in brown, catalytic triad residues are shown in yellow, and the 21 residues differing between the two cyclases (not including catalytic triad residue F300/Y300) are shown in blue, green, and red. Residues highlighted in blue (x11) represent all differences (not including F300/Y300) in the seven key regions surrounding the substrate binding pocket. AncHID2$_{AncHID1(Y300F)}$-M2 contains mutations at all of these sites (as well as the Y300F mutation). Residues highlighted in green (x4) represent additional differences not found in the seven key regions. AncHID2$_{AncHID1(Y300F)}$-M3 contains mutations at all of these sites in addition to those highlighted in blue (and the Y300F mutation). This mutant differs from AncHID1$_{Y300F}$ at only six sites (highlighted in red), which do not appear to play an important role in cyclase activity. Individual residues probed in mutants M4–M11 are labeled. (**b**) Angryline cyclization activity of wild-type and mutant cyclases expressed as turnover number (TON = mol product/mol enzyme). Colored bar height indicates the mean of four independent experiments. The results of individual experiments are represented by dots (error bars = SD). Reactions were performed at 37 °C for 30 min in 50 mM Tris (pH 9.0) with 50 μM angryline and 200 nM enzyme.

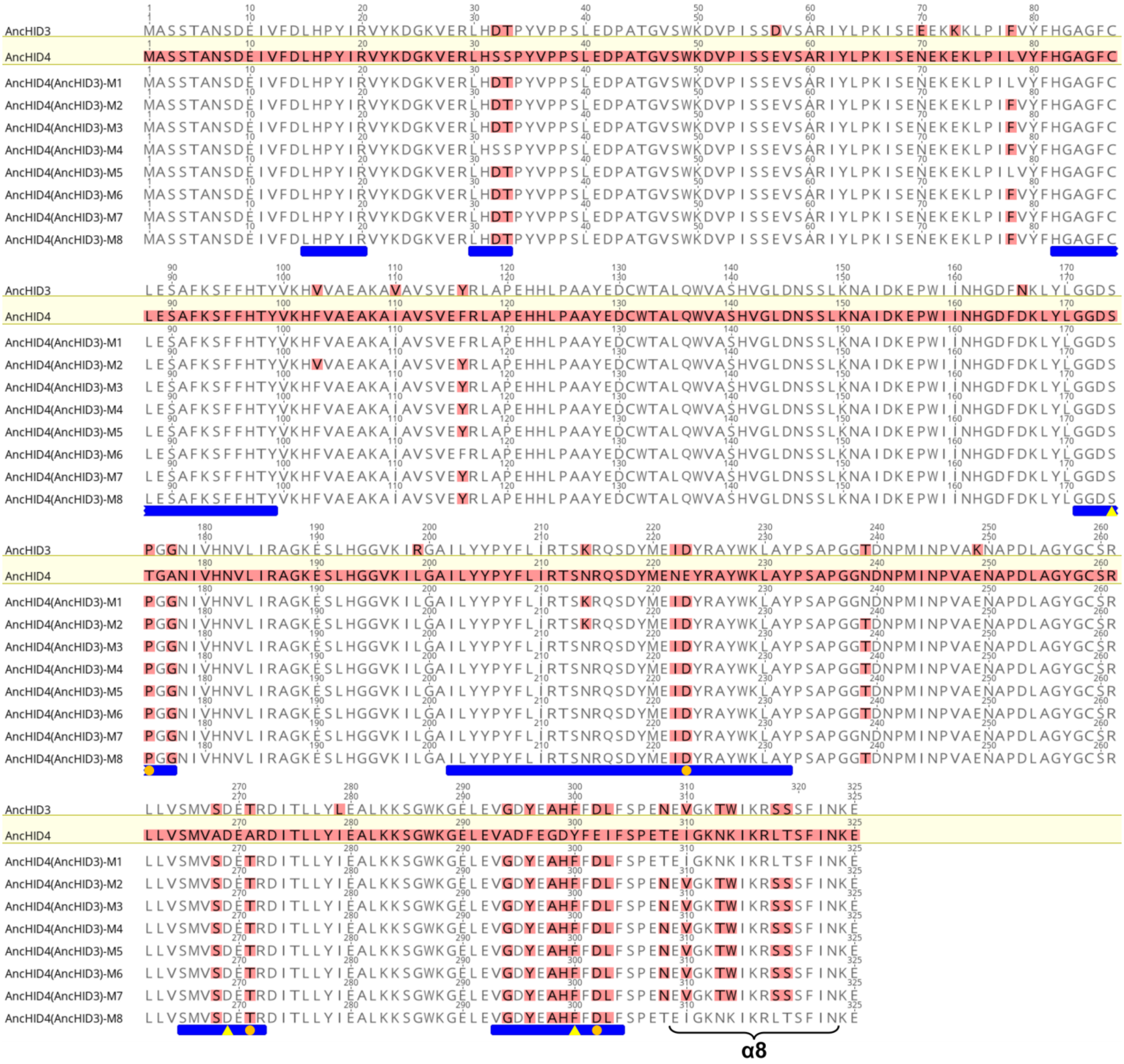

**Figure S18.** Multiple sequence alignment of AncHID3, AncHID4, and AncHID4 mutants. The template sequence (AncHID4) is enclosed in a yellow box, and all amino acid residues are highlighted in red. Among the other sequences, all differences from the template are highlighted in red. Blue bars appear below the seven contiguous regions encompassing all amino acid residues located in proximity to the substrate binding pocket. Orange circles indicate differences in residues located within 6 Å of bound substrate/product. Yellow triangles indicate the residues comprising the catalytic triad. Alpha helix 8 (α8) is labeled accordingly. See **Figure S19** for a comparison of the template (AncHID4) and target (AncHID3) structures as well as activity assay results.

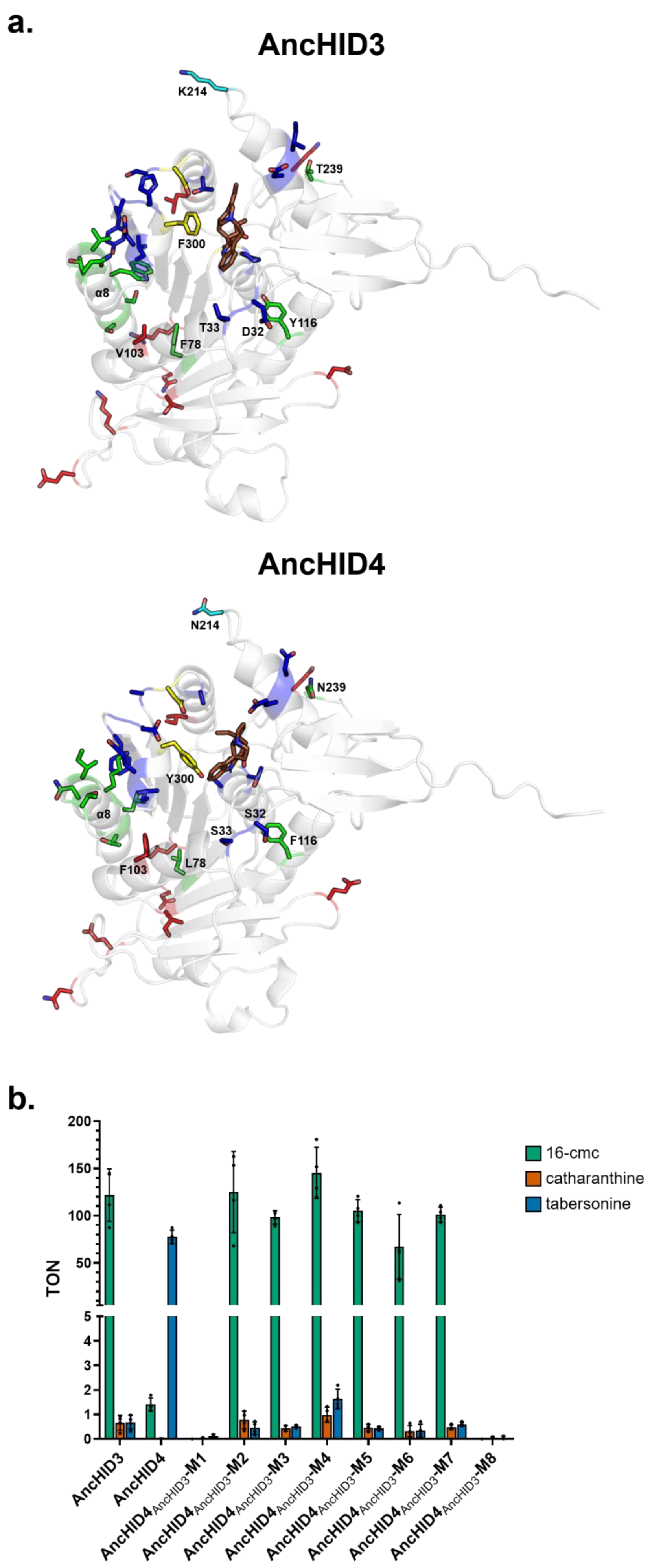


**Figure S19.** See next page for caption.

**Figure S19.** (**a**) Comparison of AncHID3 and AncHID4 (both AlphaFold models[22]). Docked product (16-cmc for AncHID3, tabersonine for AncHID4) is shown in brown, catalytic triad residues are shown in yellow, and the 33 residues differing between the two cyclases (not including catalytic triad residue F300/Y300) are shown in blue/cyan, green, and red. Residues highlighted in blue/cyan (x15) represent all differences (not including F300/Y300) in the seven key regions surrounding the substrate binding pocket. AncHID4$_{AncHID3}$-M1 contains mutations at all of these sites (as well as the Y300F mutation). Residues highlighted in green (x9) represent additional differences not found in the seven key regions. AncHID4$_{AncHID3}$-M2 contains mutations at all of these sites in addition to those highlighted in blue/cyan (plus F103V and Y300F). AncHID4$_{AncHID3}$-M3 lacks the F103V (red) and N214K (cyan) mutations and thus contains changes at 24 positions in total (all highlighted in blue and green except for F300/Y300). This mutant differs from AncHID3 at only 10 sites (highlighted in red and cyan), which do not appear to play an important role in cyclase activity. Individual residues probed in mutants M4–M8 are labeled. Alpha helix 8 (α8) located near the C-terminus contains six differences (highlighted in green), which appear to be critical for enzyme stability and cyclase activity. (**b**) Angryline cyclization activity of wild-type and mutant cyclases expressed as turnover number (TON = mol product/mol enzyme). Colored bar height indicates the mean of four independent experiments. The results of individual experiments are represented by dots (error bars = SD). Reactions were performed at 37 °C for 30 min in 50 mM Tris (pH 9.0) with 50 μM angryline and 200 nM enzyme.

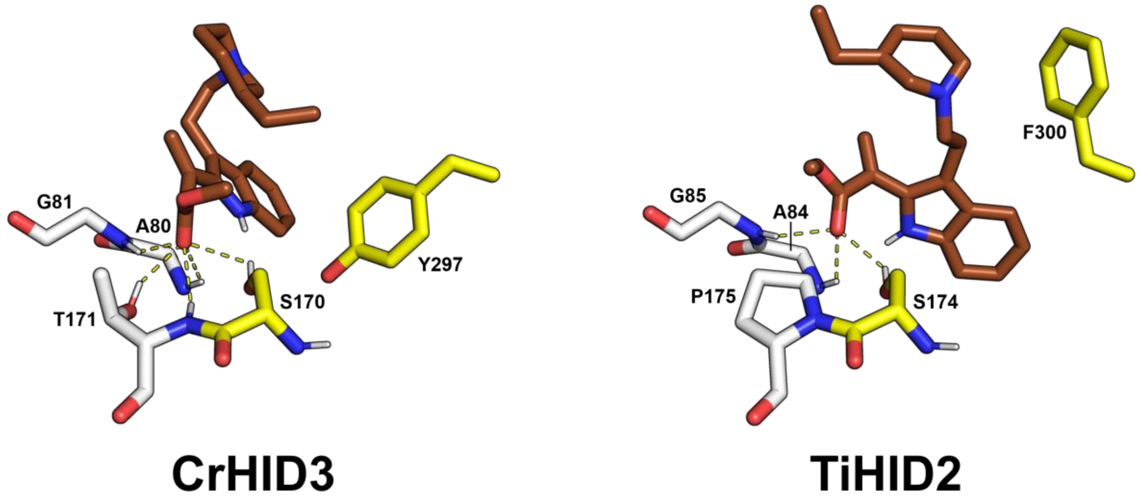


**Figure S20.** Experimental structures of CrHID3 (TS, **PDB 6RS4**) and TiHID2 (CorS, **PDB 6RJ8**) with docked substrate (dehydrosecodine) shown in brown, catalytic triad residues (Ser and Tyr/Phe) shown in yellow, oxyanion hole residues shown in white, and potential H-bonds between enzyme and substrate shown as yellow dashes. While CrHID3$_{T171}$ (equivalent to AncHID2$_{T175}$ and AncHID4$_{T175}$) possesses main chain NH and side chain OH groups that can act as H-bond donors to the carbonyl of the methyl acrylate portion of the substrate, TiHID2$_{P175}$ (equivalent to AncHID1$_{P175}$ and AncHID3$_{P175}$) is unique in lacking both of these potential H-bond donors. Thus, with the carbonyl oxygen only able to interact with the oxyanion hole of TiHID2 in a relatively superficial manner, the substrate may be more prone to orient itself such that the indole is rotated away from the oxyanion hole into a plane that is orthogonal to that occupied by the methyl acrylate. Such an orientation could further be promoted by repositioning of TiHID2$_{F300}$ (equivalent to AncHID1$_{Y300}$ and AncHID3$_{F300}$) away from the binding pocket when substrate is bound. In AncHID2 and AncHID4, the T175P mutation could cause the oxyanion hole to less effectively accommodate and interact with the carbonyl oxygen of dehydrosecodine. Repositioning of nearby residues (e.g., AncHID2$_{Y300}$ and AncHID4$_{Y300}$, equivalent to CrHID3$_{Y297}$) could accompany this change, further enabling the methyl acrylate and indole portions of dehydrosecodine to occupy orthogonal planes and shifting the product specificity away from tabersonine and toward 16-cmc.

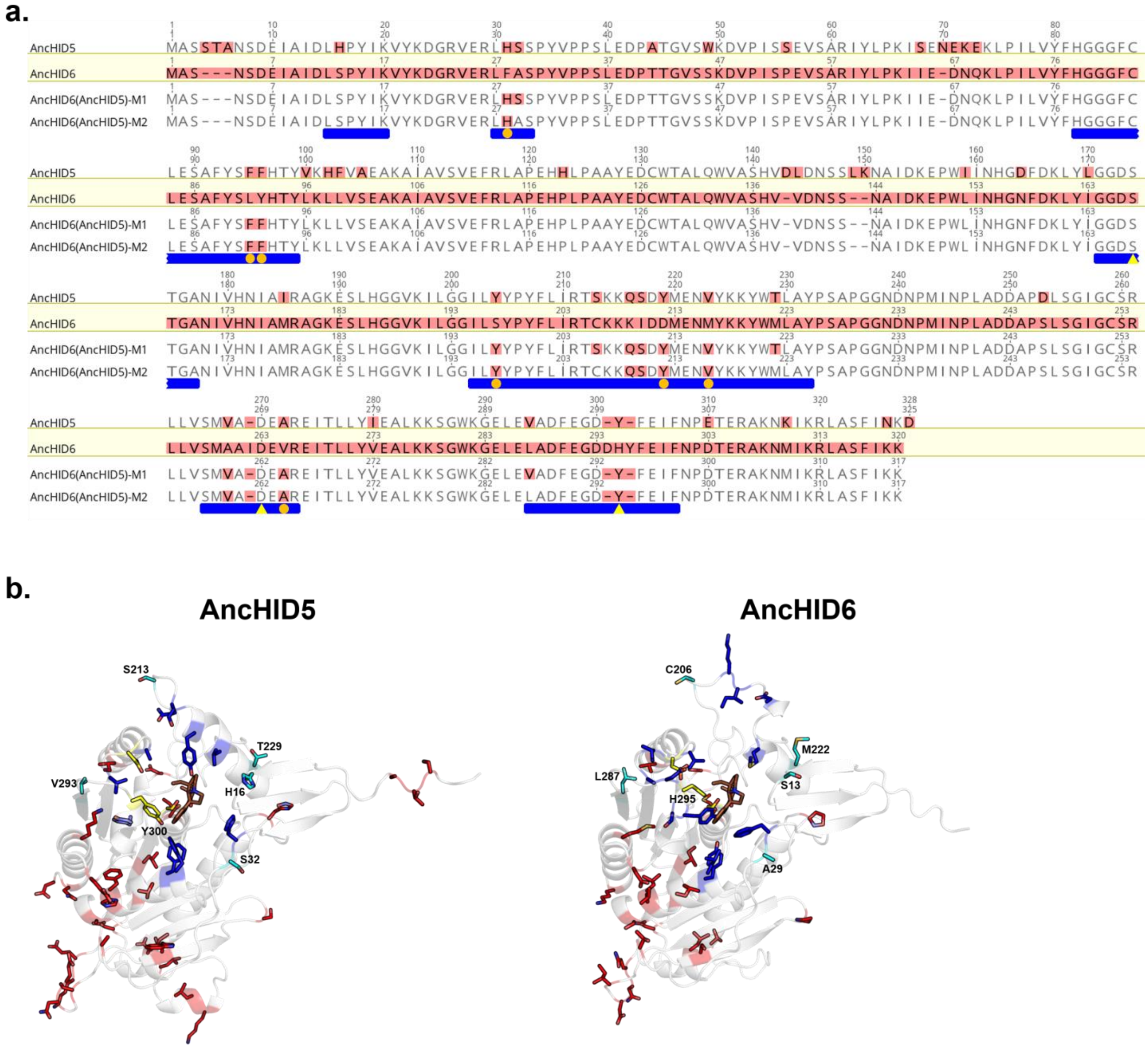


**Figure S21.** (**a**) Multiple sequence alignment of AncHID5, AncHID6, and AncHID6 mutants. The template sequence (AncHID6) is enclosed in a yellow box, and all amino acid residues are highlighted in red. Among the other sequences, all differences from the template are highlighted in red. Blue bars appear below the seven contiguous regions encompassing all amino acid residues located in proximity to the substrate binding pocket. Orange circles indicate differences in residues located within 6 Å of bound substrate/product. Yellow triangles indicate the residues comprising the catalytic triad. (**b**) Comparison of AncHID5 and AncHID6 (both AlphaFold models[22]). Docked product (tabersonine) is shown in brown, catalytic triad residues are shown in yellow, and the 48 residues differing between the two enzymes (not including catalytic triad residue Y300/H295) are shown in blue/cyan and red. Residues highlighted in blue/cyan (x18) represent all differences (not including Y300/H295) in the seven key regions surrounding the substrate binding pocket. $AncHID6_{AncHID5}$-M1 contains mutations at all of these sites (as well as the H295Y mutation) with the exception of H16/S13 (highlighted in cyan and labeled). $AncHID6_{AncHID5}$-M2 lacks mutations at four additional sites (highlighted in cyan and labeled) and thus differs from AncHID6 at a total of 14 positions (all highlighted in blue except for Y300/H295). These 14 changes are sufficient to recapitulate most of the cyclase activity observed with AncHID5 (see **Figure 5e**).

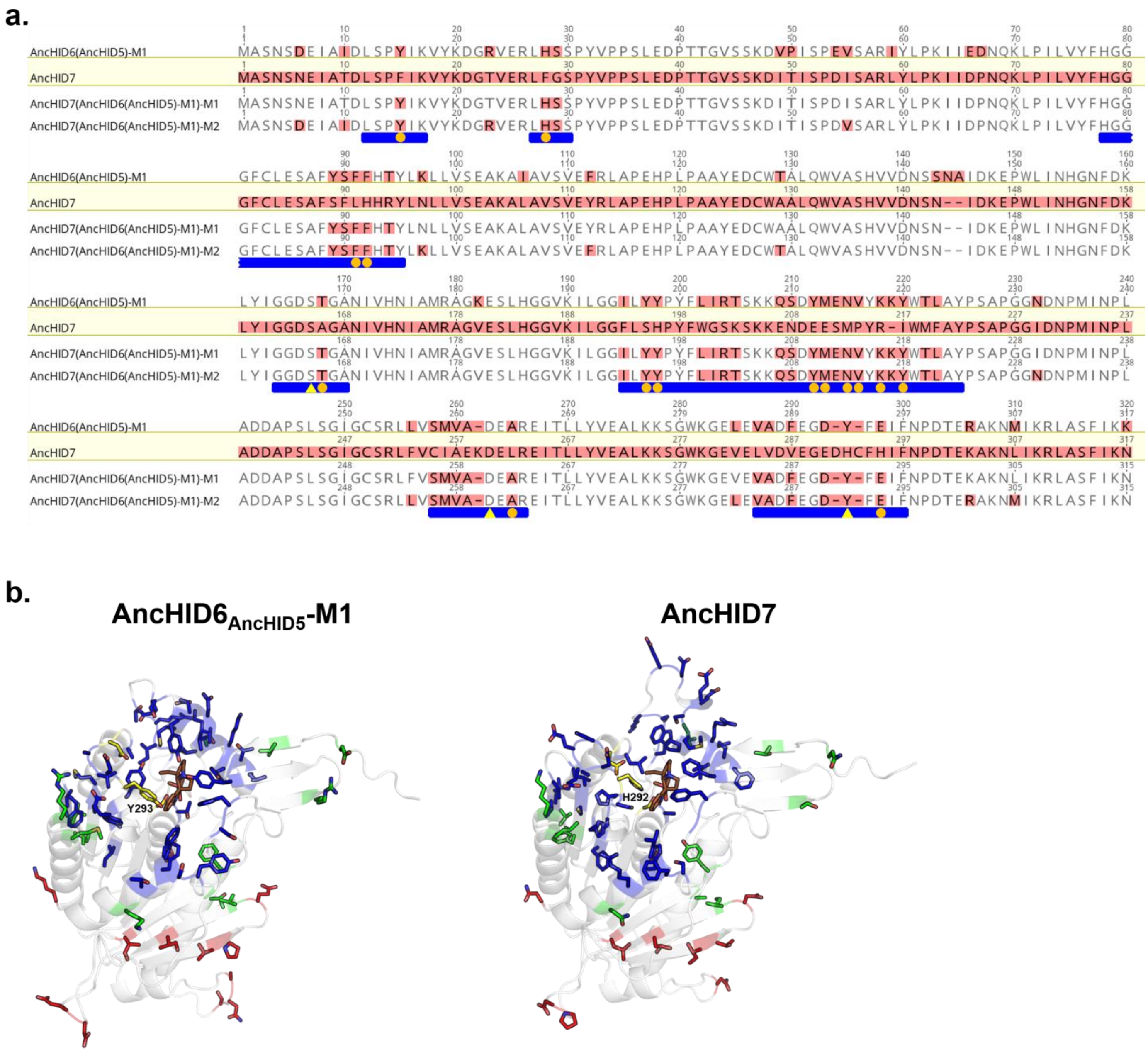


**Figure S22.** (**a**) Multiple sequence alignment of AncHID6$_{AncHID5}$-M1, AncHID7, and AncHID7 mutants. The template sequence (AncHID7) is enclosed in a yellow box, and all amino acid residues are highlighted in red. Among the other sequences, all differences from the template are highlighted in red. Blue bars appear below the seven contiguous regions encompassing all amino acid residues located in proximity to the substrate binding pocket. Orange circles indicate differences in residues located within 6 Å of bound substrate/product. Yellow triangles indicate the residues comprising the catalytic triad. (**b**) Comparison of AncHID6$_{AncHID5}$-M1 and AncHID7 (both AlphaFold models[22]). Docked product (tabersonine) is shown in brown, catalytic triad residues are shown in yellow, and the 65 residues differing between the two enzymes (not including catalytic triad residue Y293/H292) are shown in blue, green, and red. Residues highlighted in blue (x41) represent all differences (not including Y293/H292) in the seven key regions surrounding the substrate binding pocket. AncHID7$_{AncHID6(AncHID5)-M1}$-M1 contains mutations at all of these sites (as well as the H292Y mutation). Residues highlighted in green (x12) represent additional differences not found in the seven key regions. AncHID7$_{AncHID6(AncHID5)-M1}$-M2 contains mutations at all of these sites in addition to those highlighted in blue (plus H292Y). This mutant differs from the target AncHID6$_{AncHID5}$-M1 at only 12 sites (highlighted in red). See **Figure 5e** for results of activity assays with angryline.

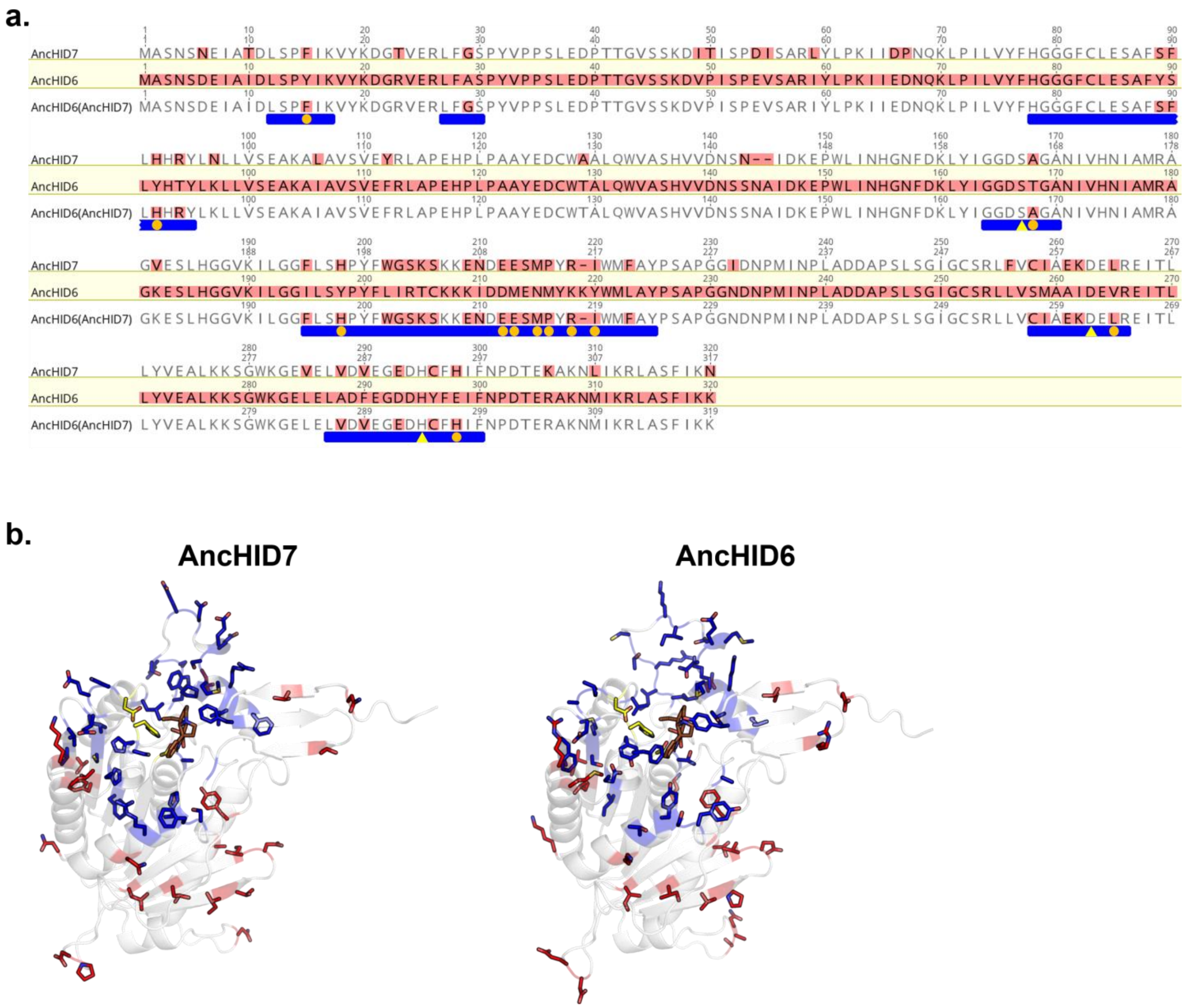


**Figure S23.** (**a**) Multiple sequence alignment of AncHID7, AncHID6, and the $AncHID6_{AncHID7}$ mutant. The template sequence (AncHID6) is enclosed in a yellow box, and all amino acid residues are highlighted in red. Among the other sequences, all differences from the template are highlighted in red. Blue bars appear below the seven contiguous regions encompassing all amino acid residues located in proximity to the substrate binding pocket. Orange circles indicate differences in residues located within 6 Å of bound substrate/product. Yellow triangles indicate the residues comprising the catalytic triad. (**b**) Comparison of AncHID7 and AncHID6 (both AlphaFold models[22]). Docked tabersonine is shown in brown, catalytic triad residues are shown in yellow, and the 59 residues differing between the two enzymes are shown in blue and red. Residues highlighted in blue (x35) represent all differences in the seven key regions surrounding the substrate binding pocket. $AncHID6_{AncHID7}$ contains mutations at all of these sites and exhibits high-level esterase activity (see **Table S4**).

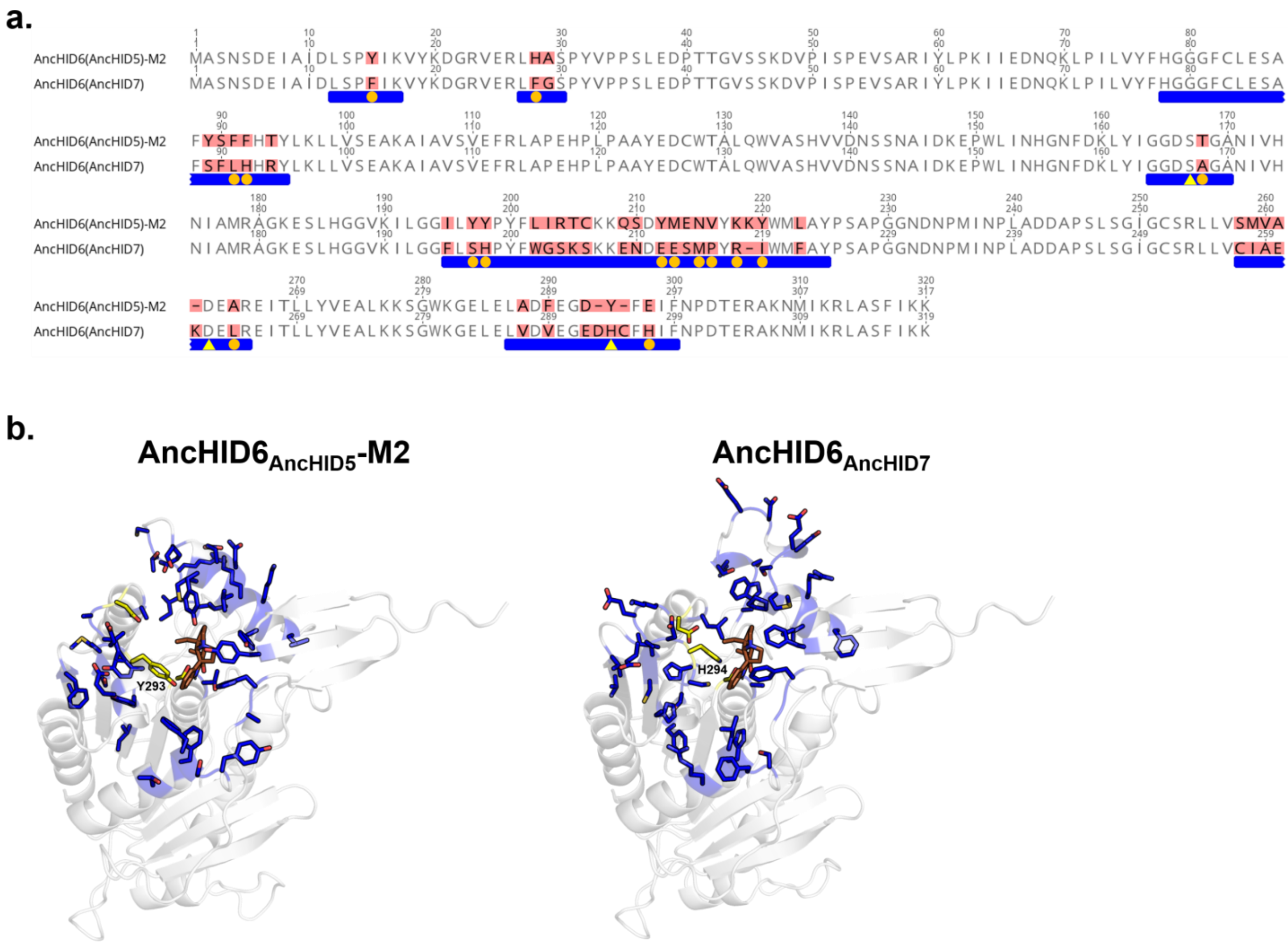


**Figure S24.** (**a**) Pairwise sequence alignment of AncHID6$_{AncHID5}$-M2 (cyclase with TS activity) and AncHID6$_{AncHID7}$ (CXE with esterase activity). All differences from the consensus sequence are highlighted in red. Blue bars appear below the seven contiguous regions encompassing all amino acid residues located in proximity to the substrate binding pocket. Orange circles indicate differences in residues located within 6 Å of bound substrate/product. Yellow triangles indicate the residues comprising the catalytic triad. (**b**) Comparison of AncHID6$_{AncHID5}$-M2 and AncHID6$_{AncHID7}$ (both AlphaFold models[22]). Docked tabersonine is shown in brown, catalytic triad residues are shown in yellow, and the 40 residues differing between the two enzymes (not including catalytic triad residue Y293/H294) are shown in blue. These residues (plus Y293/H294) also represent all differences in the seven key regions surrounding the substrate binding pocket. Thus, changing a total of 41 residues in these regions near the binding pocket is sufficient to generate a cyclase (TS) from a CXE, and vice versa.

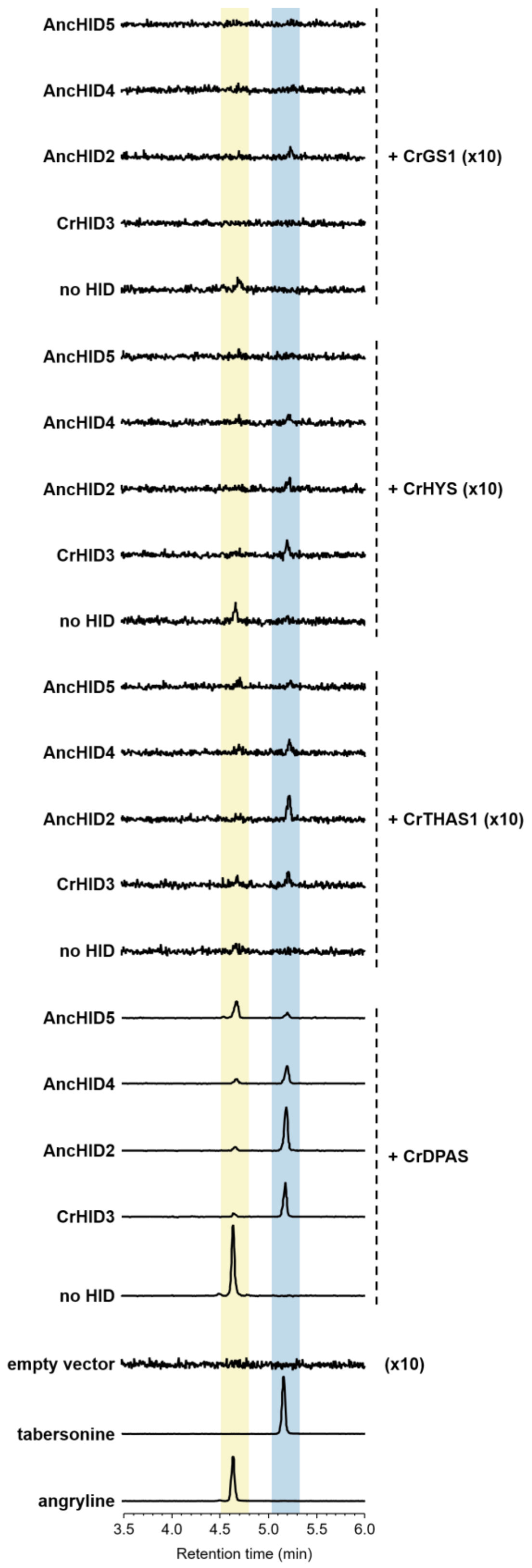


**Figure S25.** Reconstitution of dehydrosecodine (angryline) and tabersonine biosynthesis from stemmadenine acetate in *Nicotiana benthamiana* (trial 2). LC-MS chromatograms (EIC 337.1911 ± 0.01) of leaf extracts following infiltration of CrDPAS, CrTHAS1, CrHYS, or CrGS1 alone (no HID) or with extant (CrHID3) or ancestral (AncHID2, AncHID4, AncHID5) TS are shown. Signal intensity is magnified 10x for the empty vector, CrTHAS1, CrHYS, and CrGS1 samples.

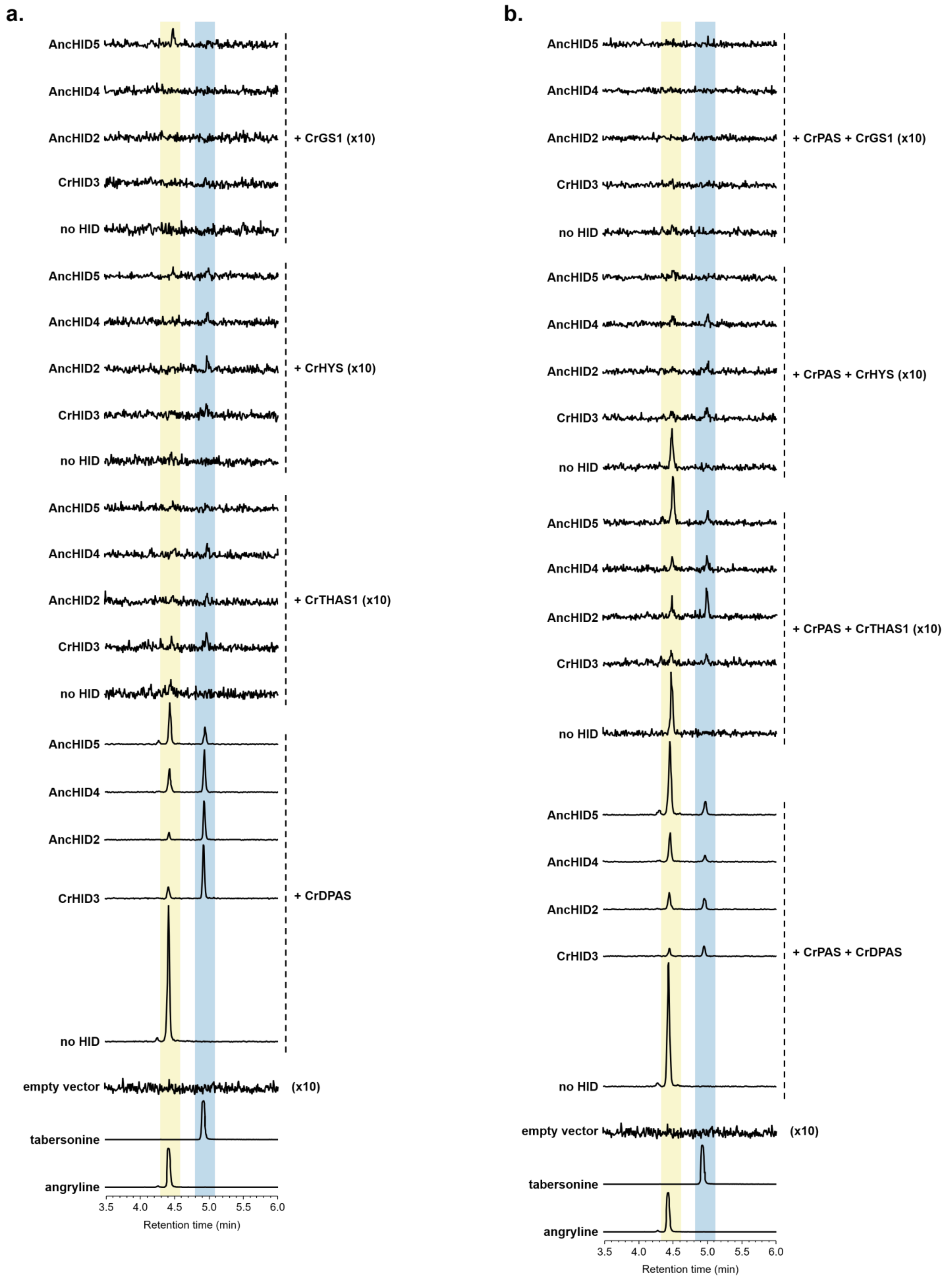


**Figure S26.** Reconstitution of dehydrosecodine (angryline) and tabersonine biosynthesis from stemmadenine acetate in *Nicotiana benthamiana* (trial 3). LC-MS chromatograms (EIC 337.1911 ± 0.01) of leaf extracts following infiltration of CrDPAS, CrTHAS1, CrHYS, or CrGS1 alone (no HID) or with extant (CrHID3) or ancestral (AncHID2, AncHID4, AncHID5) TS are shown. Signal intensity is magnified 10x for the empty vector, CrTHAS1, CrHYS, and CrGS1 samples. (**a**) Without co-infiltration of CrPAS. (**b**) With co-infiltration of CrPAS.

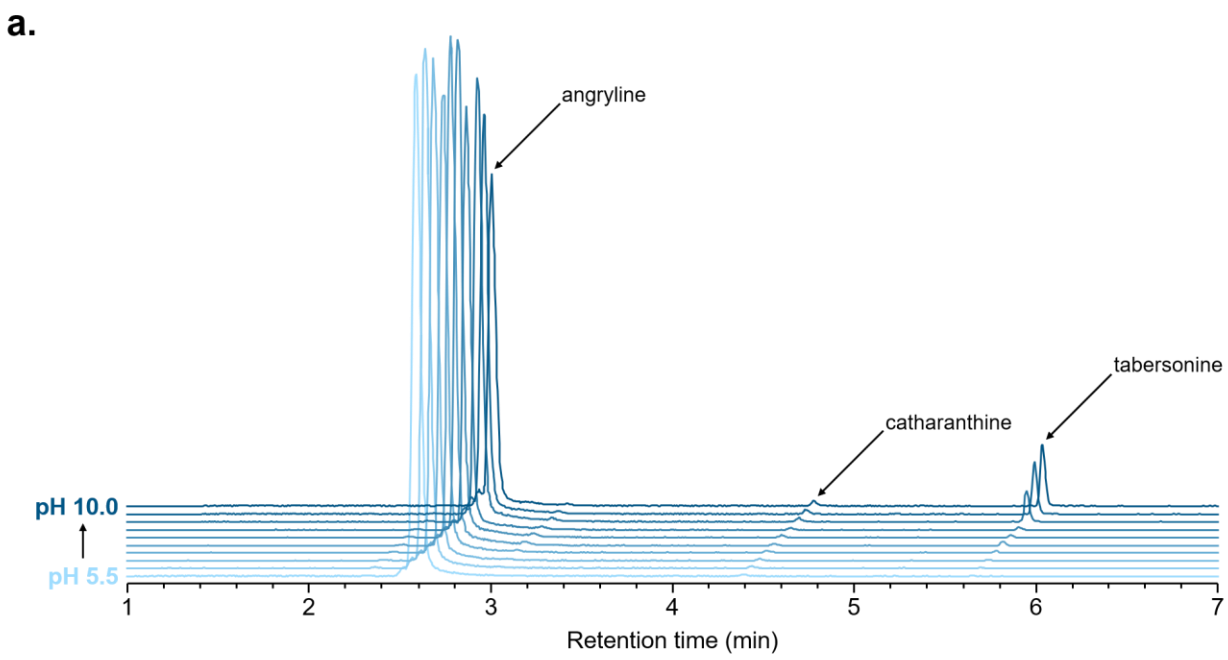


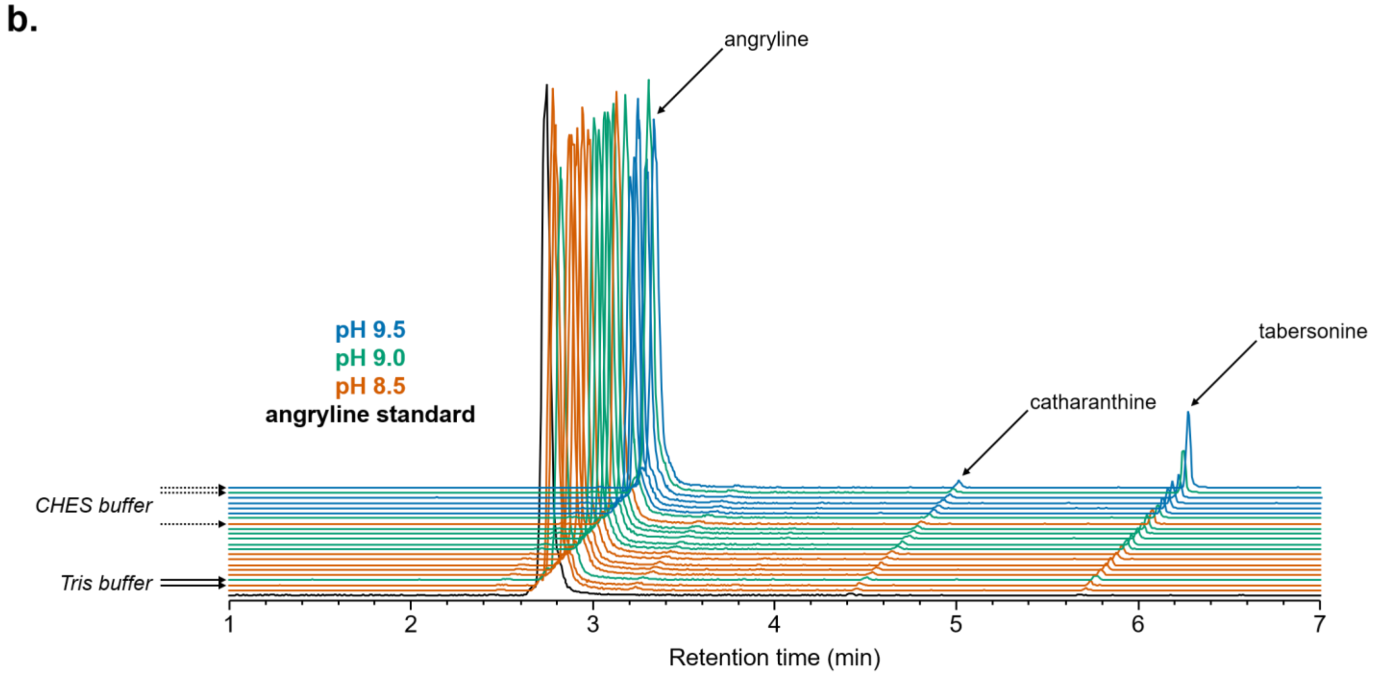


**Figure S27.** Spontaneous cyclization of angryline in solution. (**a**) LC-MS traces (EIC 337.1911 ± 0.01) of angryline (50 μM) following incubation in different buffers (50 mM) at 37 °C for 30 min. Buffers employed: MES (pH 5.5), MES (pH 6.0), MES (pH 6.5), HEPES (pH 7.0), HEPES (pH 7.5), Tris (pH 8.0), Tris (pH 8.5), CHES (pH 9.0), CHES (pH 9.5), CHES (pH 10.0). Tabersonine forms spontaneously from angryline in solution at a significant level when pH ≥ 9.0. Low levels of catharanthine are apparent in all samples, but these increase only marginally at higher pH. 16-cmc (retention time ~2.0 min) is not observed in any of the samples. (**b**) LC-MS traces (EIC 337.1911 ± 0.01) of angryline (50 μM) following incubation in different buffers (50 mM) at 37 °C for 1 h. Chromatograms are organized (front-to-back / bottom-to-top) in order of increasing levels of tabersonine: angryline standard, Glycine (pH 8.5), Tris (pH 8.5), Tris (pH 9.0), AMPD (pH 8.5), Bis-tris propane (pH 8.5), TAPS (pH 8.5), AMPSO (pH 8.5), Bicine (pH 8.5), Glycine (pH 9.0), AMPD (pH 9.0), TAPS (pH 9.0), Bis-tris propane (pH 9.0), AMPSO (pH 9.0), CHES (pH 8.5), Bicine (pH 9.0), AMPD (pH 9.5), Bis-tris propane (pH 9.5), AMPSO (pH 9.5), Glycine (pH 9.5), CHES (pH 9.0), CHES (pH 9.5). Tabersonine forms spontaneously from angryline in solution at a significant level when pH ≥ 9.0, especially in CHES buffer (dotted arrows). The lowest levels of tabersonine are observed when angryline is incubated in Tris buffer (solid arrows). Catharanthine is observed as a trace contaminant in the angryline standard, and levels appear to increase only marginally at higher pH. 16-cmc (retention time ~2.1 min) is not observed in any of the samples.

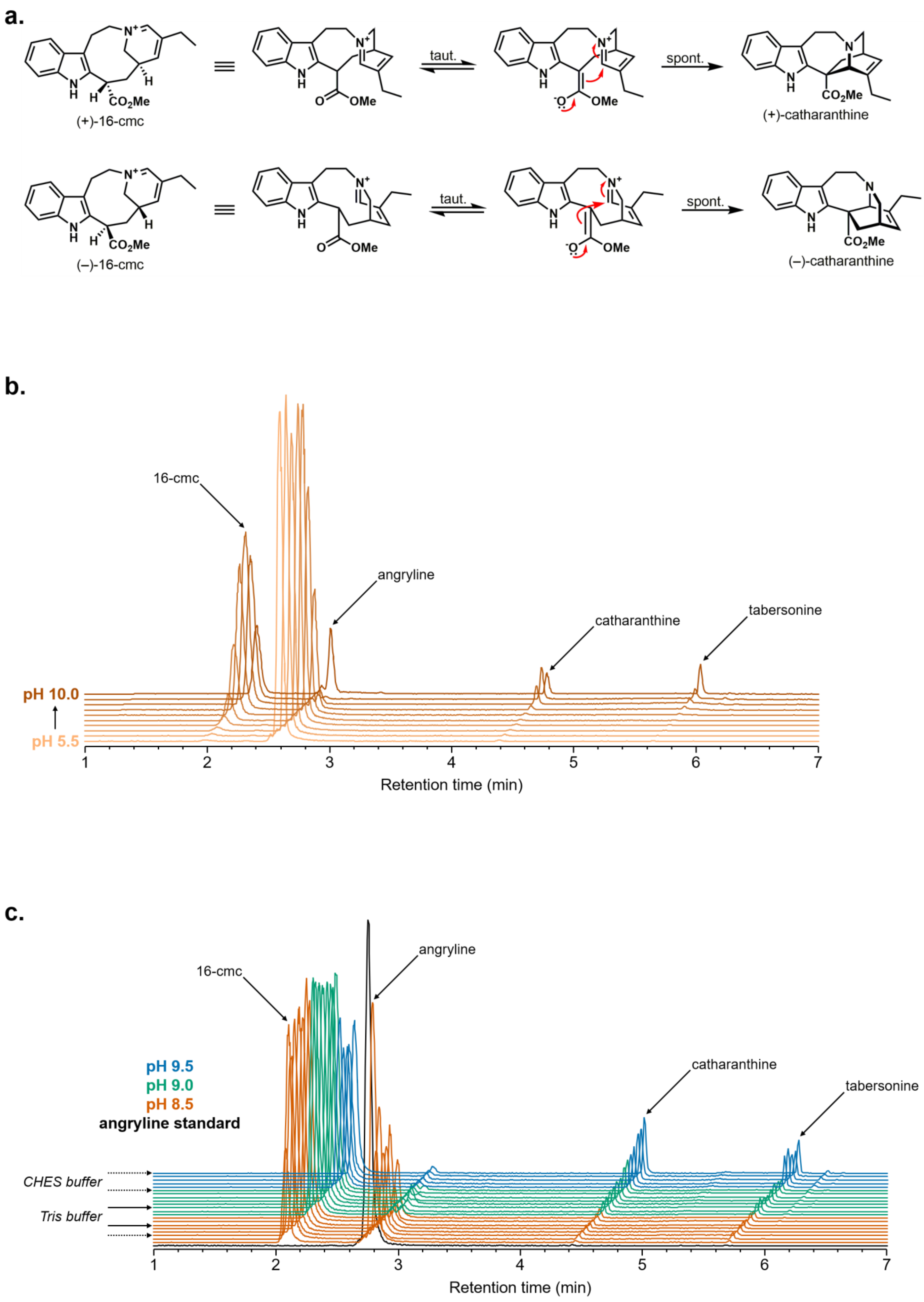


**Figure S28.** Continued on next page…

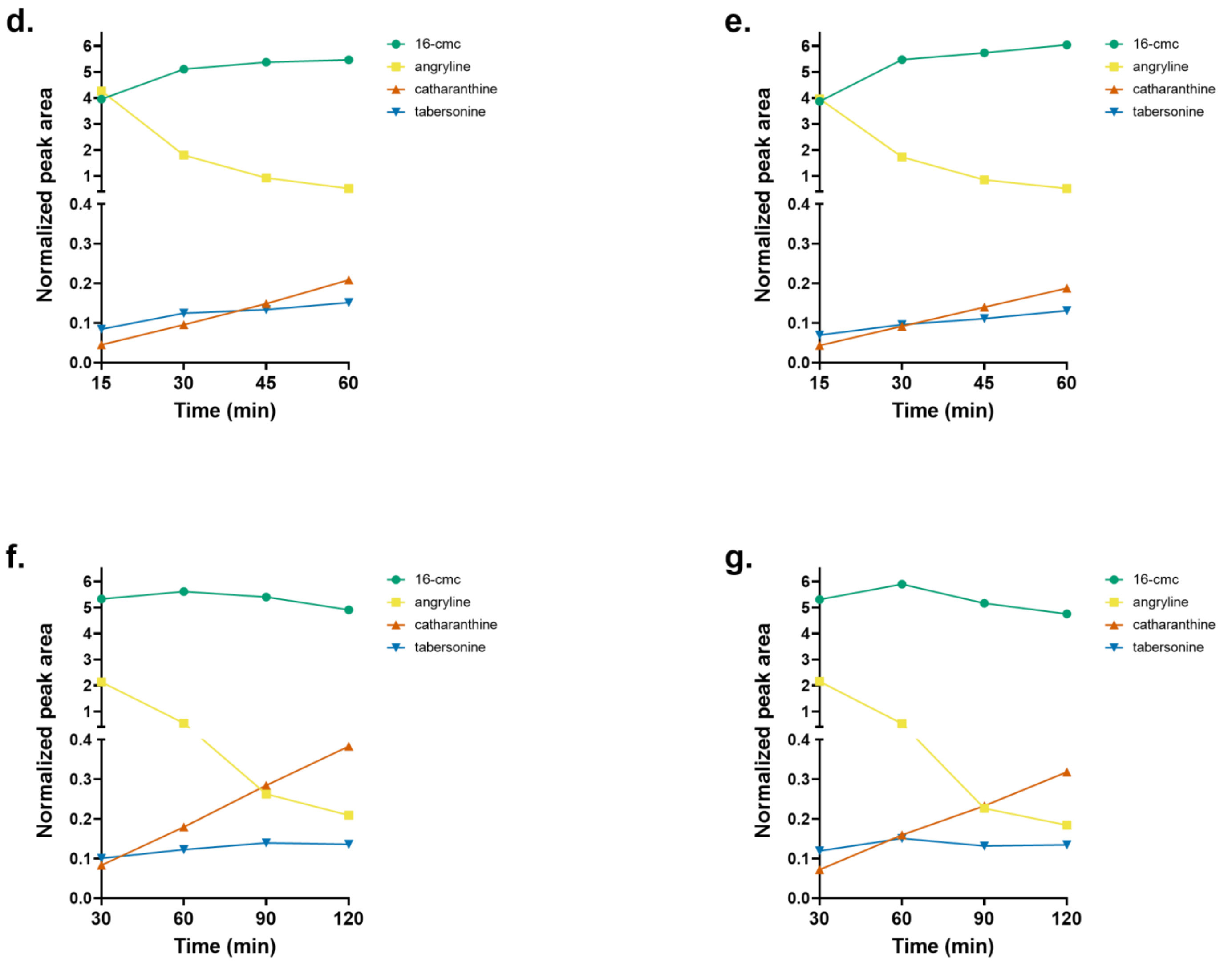


**Figure S28.** Evidence for spontaneous cyclization of 16-cmc in solution. (**a**) Spontaneous cyclization of (+)-16-cmc or (–)-16-cmc leads to (+)-catharanthine or (–)-catharanthine, respectively. As TiHID2 is known to generate (–)-16-cmc from angryline, it is hypothesized that the additional cyclization product observed in reactions with this enzyme is (–)-catharanthine. However, due to its low abundance, this compound has not been isolated and structurally characterized. Note that (+)-catharanthine is the naturally occurring enantiomer produced by *C. roseus*. (**b**) LC-MS traces (EIC 337.1911 ± 0.01) of reactions between angryline (50 µM) and TiHID2 (1 µM) performed in different buffers (50 mM) at 37 °C for 30 min. For the list of buffers employed, see **Figure S27**. Minimal reaction occurs at low pH due to the relatively high stability of angryline under these conditions. Higher pH promotes a shift in the equilibrium toward dehydrosecodine (see **Figure 3a**), which TiHID2 cyclizes to form 16-cmc. Although 16-cmc readily accumulates when pH ≥ 8.0, catharanthine appears as a significant byproduct only when pH ≥ 9.0. (**c**) LC-MS traces (EIC 337.1911 ± 0.01) of reactions between angryline (50 µM) and TiHID2 (1 µM) performed in different buffers (50 mM) at 37 °C for 1 h. Chromatograms are organized (front-to-back / bottom-to-top) in order of increasing levels of catharanthine: angryline standard, Glycine (pH 8.5), AMPSO (pH 8.5), CHES (pH 8.5), TAPS (pH 8.5), Bis-tris propane (pH 8.5), Tris (pH 8.5), Bicine (pH 8.5), AMPD (pH 8.5), TAPS (pH 9.0), AMPSO (pH 9.0), Tris (pH 9.0), Glycine (pH 9.0), Bicine (pH 9.0), Bis-tris propane (pH 9.0), AMPD (pH 9.0), CHES (pH 9.0), AMPSO (pH 9.5), Bis-tris propane (pH 9.5), AMPD (pH 9.5), Glycine (pH 9.5), CHES (pH 9.5). Significant levels of unreacted angryline remain at pH 8.5. Accumulation of catharanthine correlates with higher pH values and occurs preferentially in the presence of certain buffers (e.g., CHES (dotted arrows)). Tris (solid arrows) was selected as the buffer of choice for performing angryline cyclization assays due to the high levels of substrate consumption (at pH 9.0) and the low levels of spontaneous cyclization products observed (see also **Figure S27b**). (**d**) Normalized peak area (internal standard: vindoline) over time for reactions between angryline (50 µM) and TiHID2 (1 µM) performed in 50 mM Tris (pH 9.0) at 37 °C. (**e**) Same as **d** with the exception that reactions were performed with 5 µM TiHID2. (**f**) Same as **e** with the exception that reactions were performed over a period of 120 min. (**g**) Same as **f** with the exception that reactions were performed with 10 µM TiHID2. Maximum levels of 16-cmc are reached at 60 min. While tabersonine levels do not increase substantially after 60 min, catharanthine continues to accumulate at a constant rate even after most of the angryline has been consumed. Moreover, this rate appears to be independent of the enzyme concentration. These observations suggest that catharanthine forms spontaneously in solution from 16-cmc.

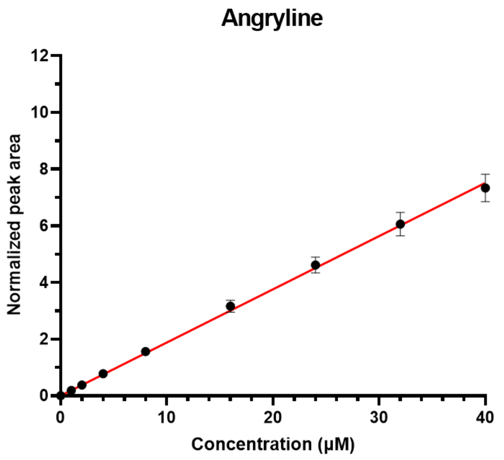


y = mx+b
m = 0.1878
b = 0
$R^2$ = 0.9905

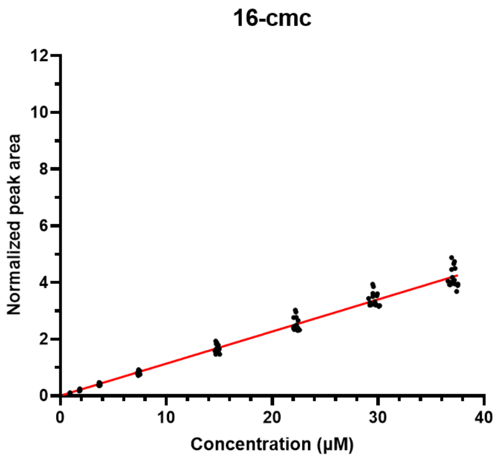


y = mx+b
m = 0.1133
b = 0
$R^2$ = 0.9868

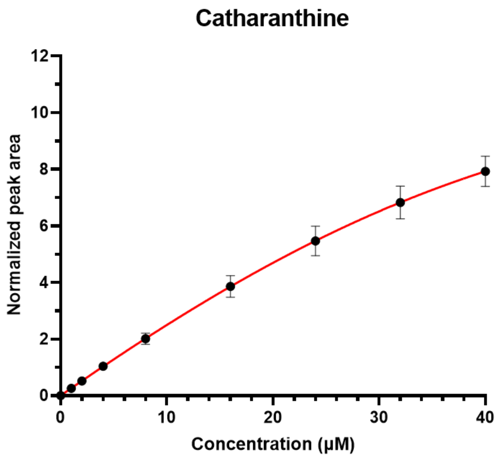


y = (a((b+x+c)-sqrt(((b+x+c)$^2$)-(4bx))))/(2b)
a = 12.93
b = 39.43
c = 9.978
$R^2$ = 0.9860

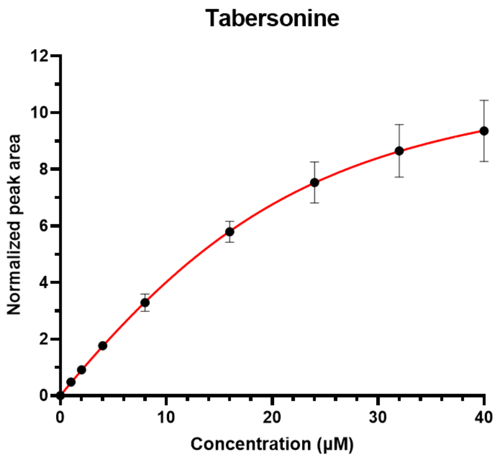


y = (a((b+x+c)-sqrt(((b+x+c)$^2$)-(4bx))))/(2b)
a = 12.12
b = 19.16
c = 7.414
$R^2$ = 0.9761

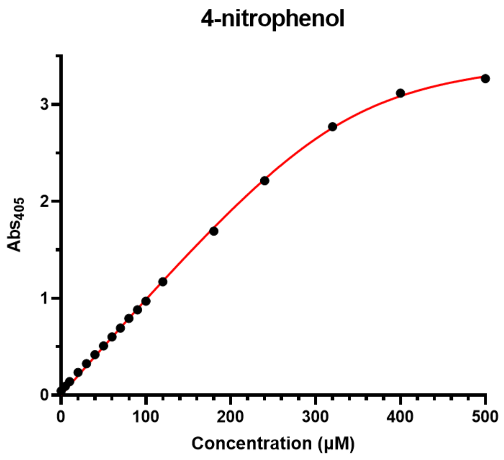


y = (a((b+x+c)-sqrt(((b+x+c)$^2$)-(4bx))))/(2b)
a = 3.611
b = 340.4
c = 18.51
$R^2$ = 0.9984

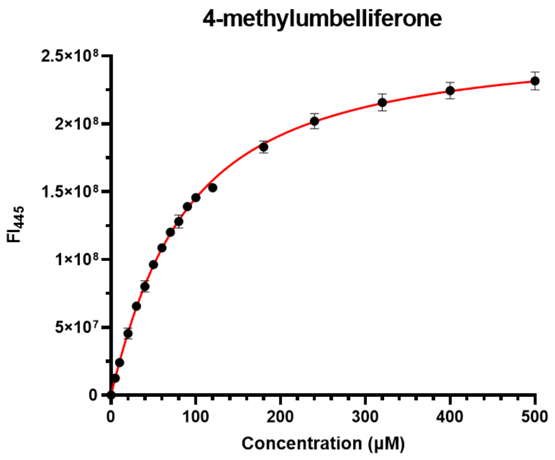


y = (a((b+x+c)-sqrt(((b+x+c)$^2$)-(4bx))))/(2b)
a = 263827277
b = 34.97
c = 65.79
$R^2$ = 0.9973

**Figure S29.** See next page for caption.

**Figure S29.** Calibration curves generated for angryline, 16-cmc, catharanthine, tabersonine, 4-nitrophenol, and 4-methylumbelliferone. For all compounds except for 16-cmc, black data points represent the mean signal intensity (error bars = SD), and the red line represents the calibration curve calculated using the given linear or nonlinear (quadratic) equation. As the estimated concentrations of 16-cmc varied each time the standards were tested, individual data points rather than mean ± SD are shown.

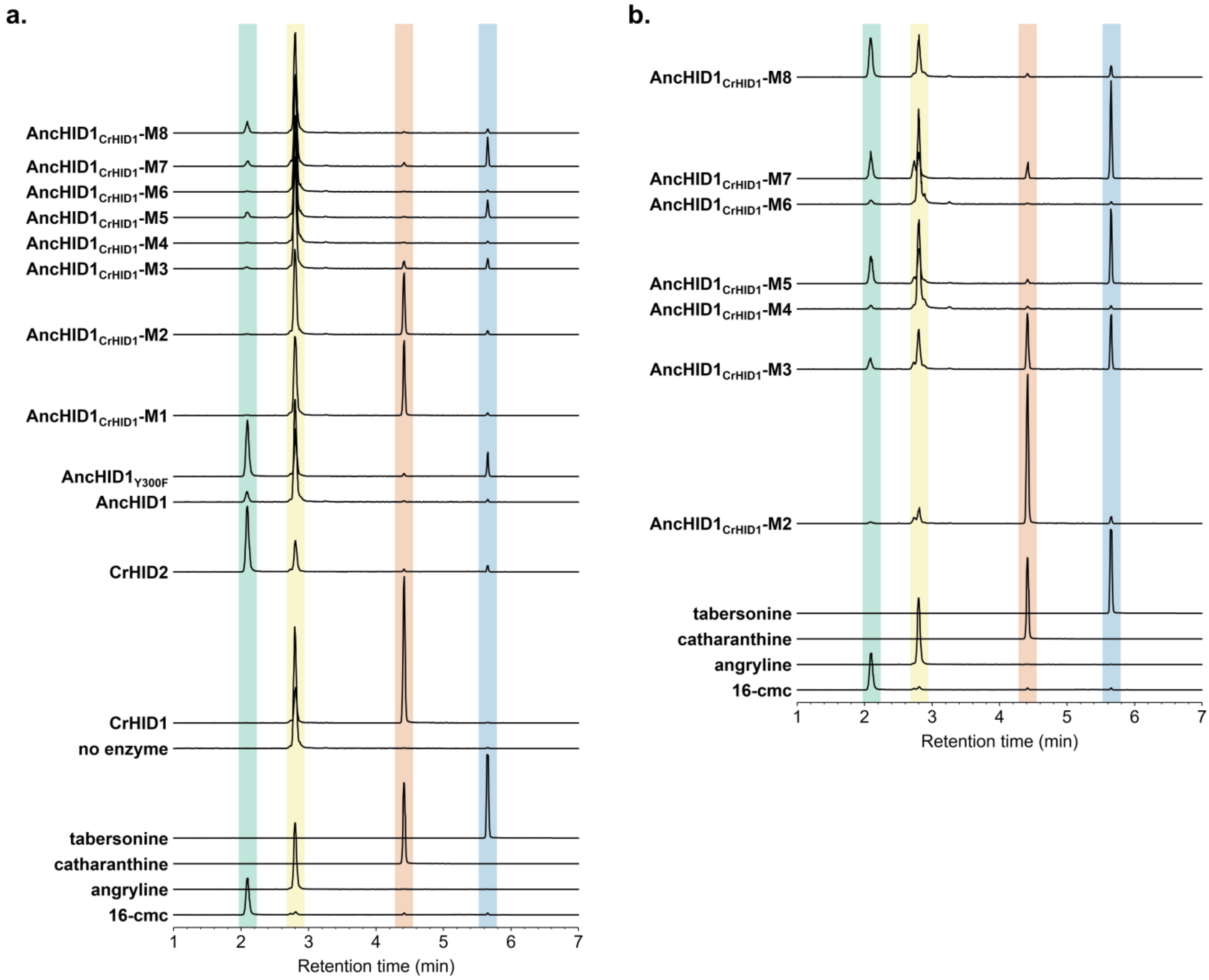


**Figure S30.** LC-MS traces (EIC 337.1911 ± 0.01) of reactions between angryline and CrHID1, CrHID2, AncHID1, and AncHID1 mutants. (**a**) Reactions performed using 1 µM enzyme. (**b**) Selected reactions performed using 5 µM enzyme.

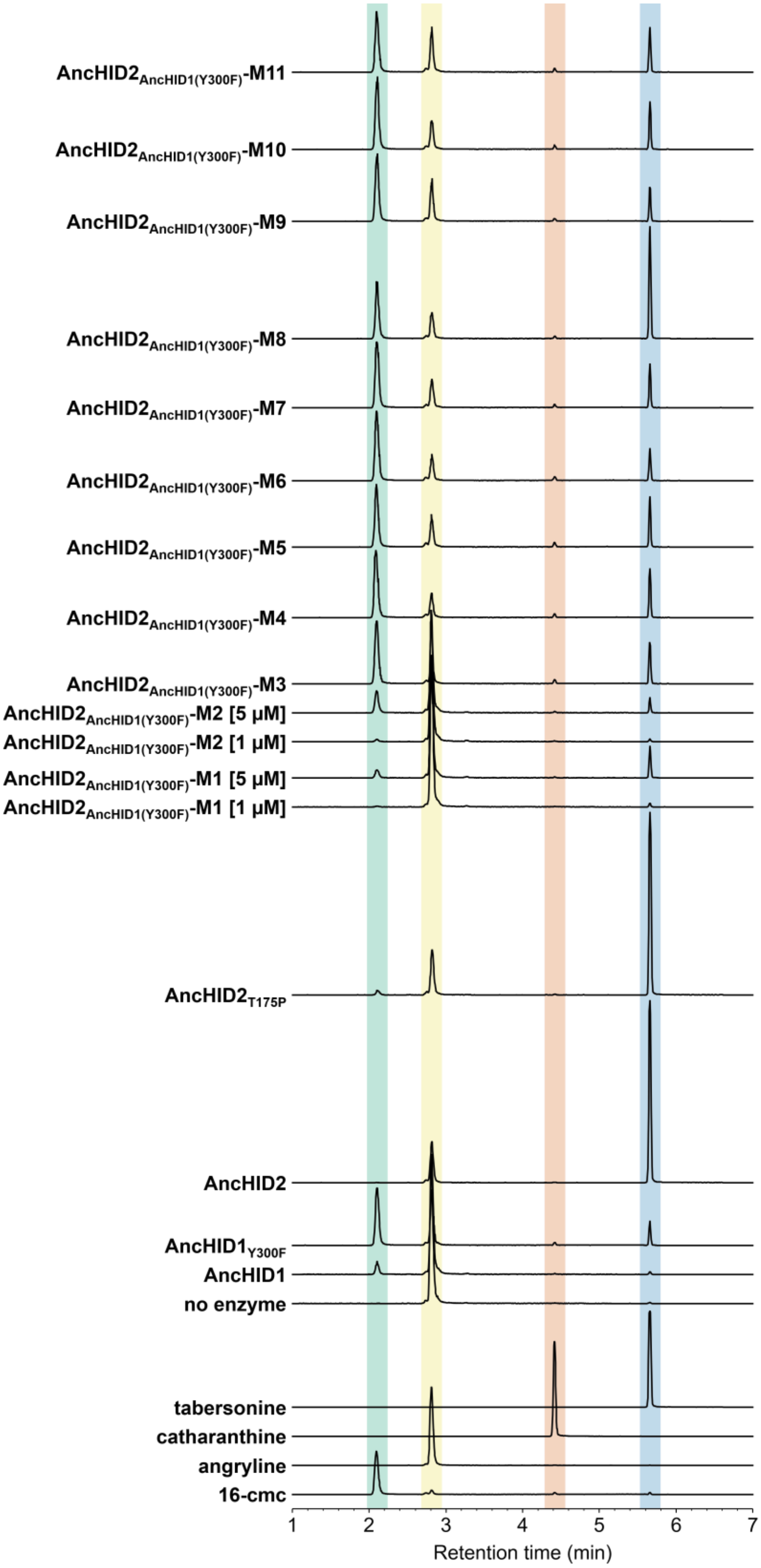


**Figure S31.** LC-MS traces (EIC 337.1911 ± 0.01) of reactions between angryline and AncHID1, AncHID1$_{Y300F}$, AncHID2, and AncHID2 mutants. Unless otherwise noted, all reactions were performed using 1 μM enzyme.

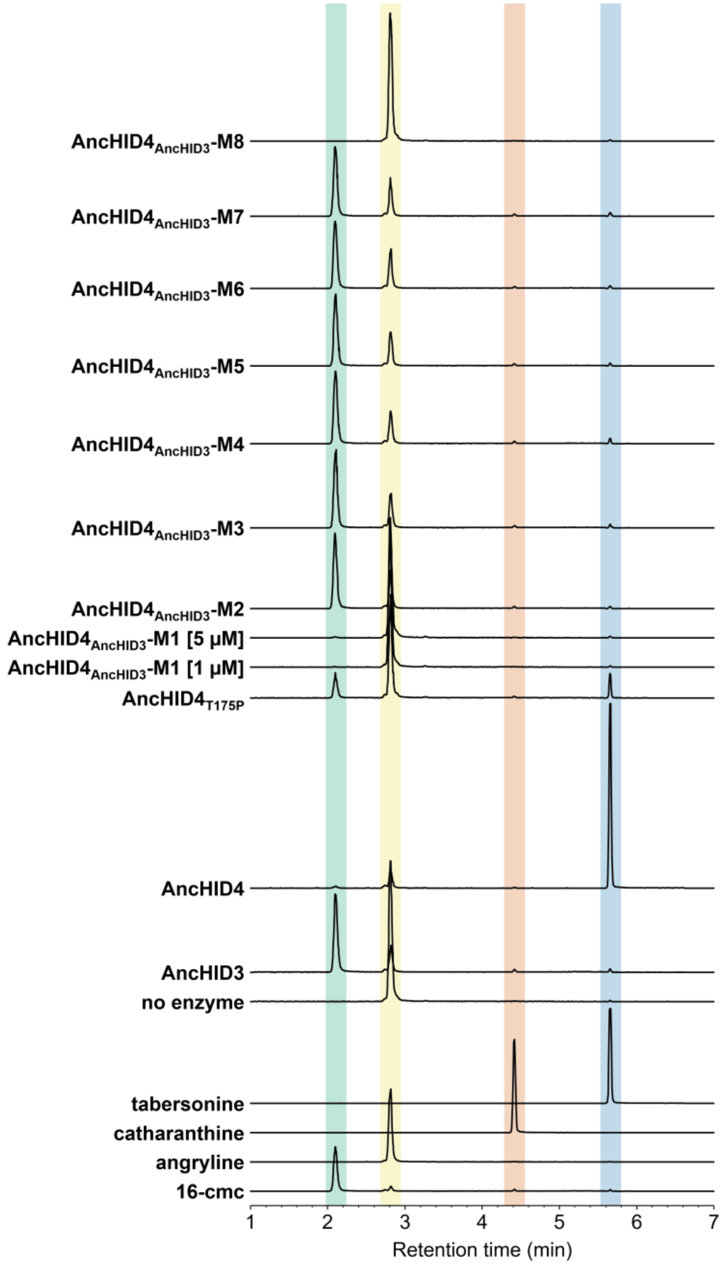


**Figure S32.** LC-MS traces (EIC 337.1911 ± 0.01) of reactions between angryline and AncHID3, AncHID4, and AncHID4 mutants. Unless otherwise noted, all reactions were performed using 1 μM enzyme.

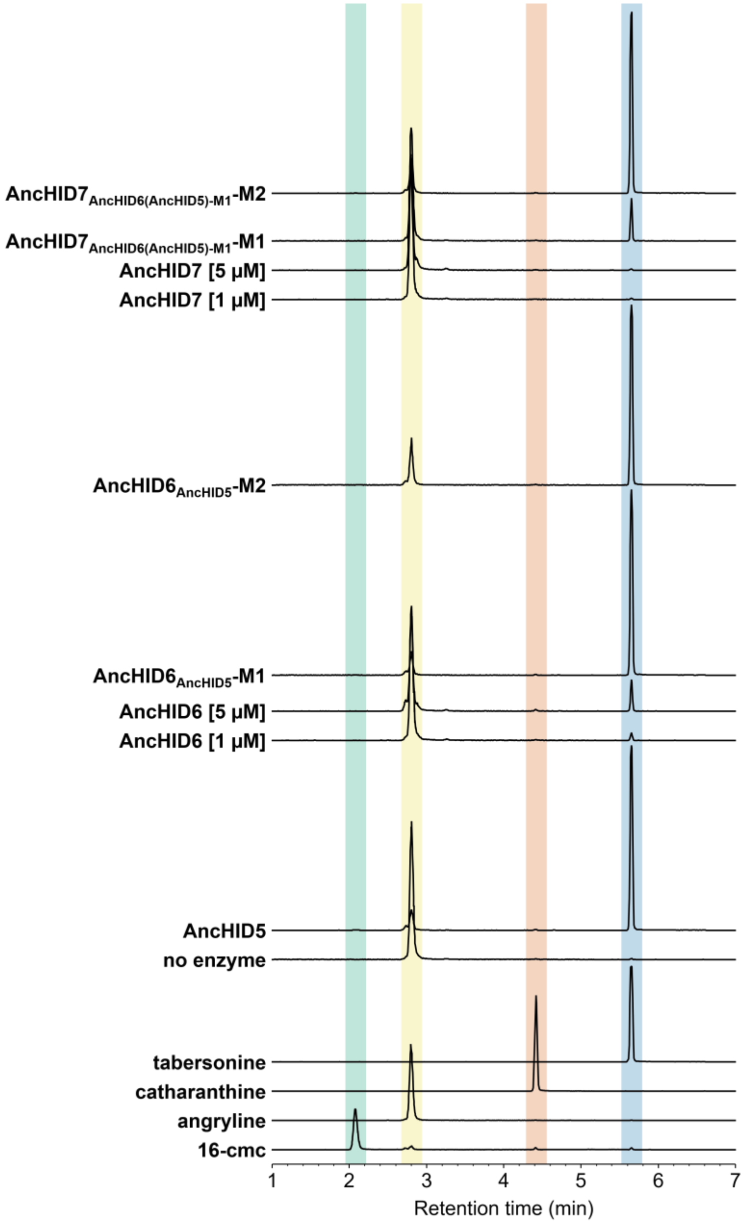


**Figure S33.** LC-MS traces (EIC 337.1911 ± 0.01) of reactions between angryline and AncHID5, AncHID6, AncHID6 mutants, AncHID7, and AncHID7 mutants. Unless otherwise noted, all reactions were performed using 1 μM enzyme.

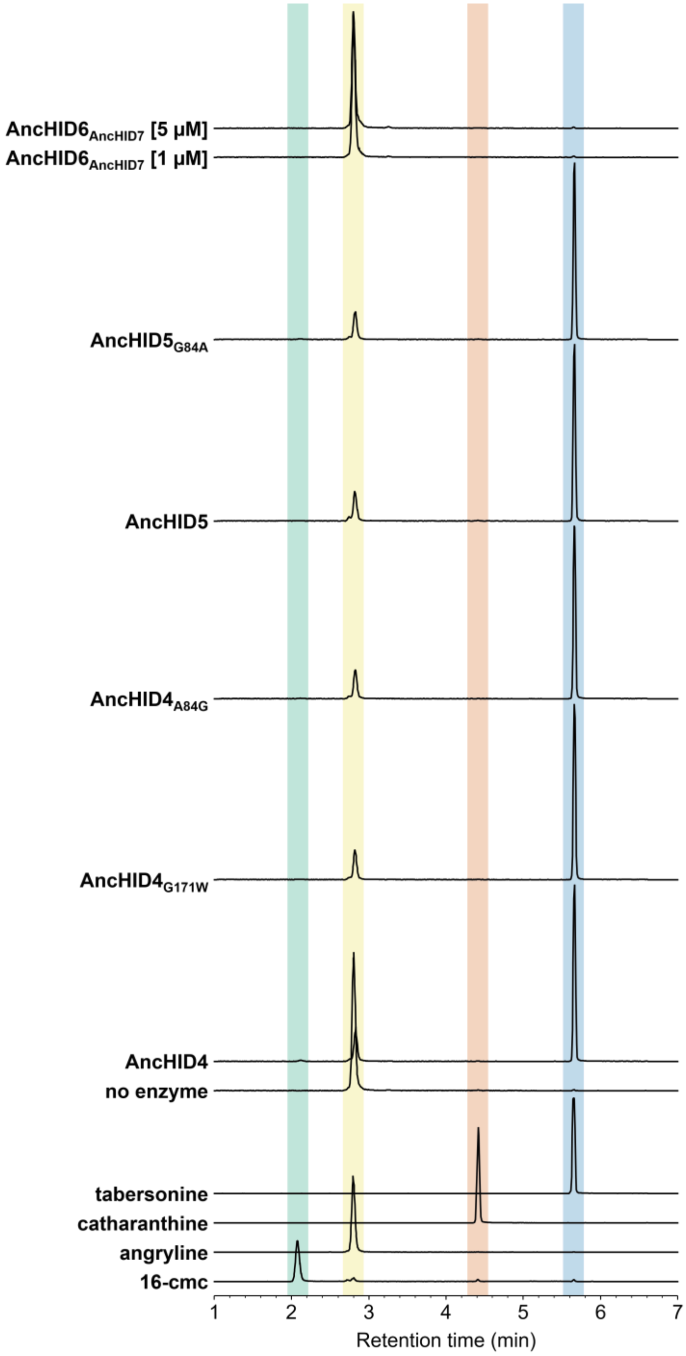


**Figure S34.** LC-MS traces (EIC 337.1911 ± 0.01) of reactions between angryline and AncHID mutants not shown in previous figures. Unless otherwise noted, all reactions were performed using 1 μM enzyme.

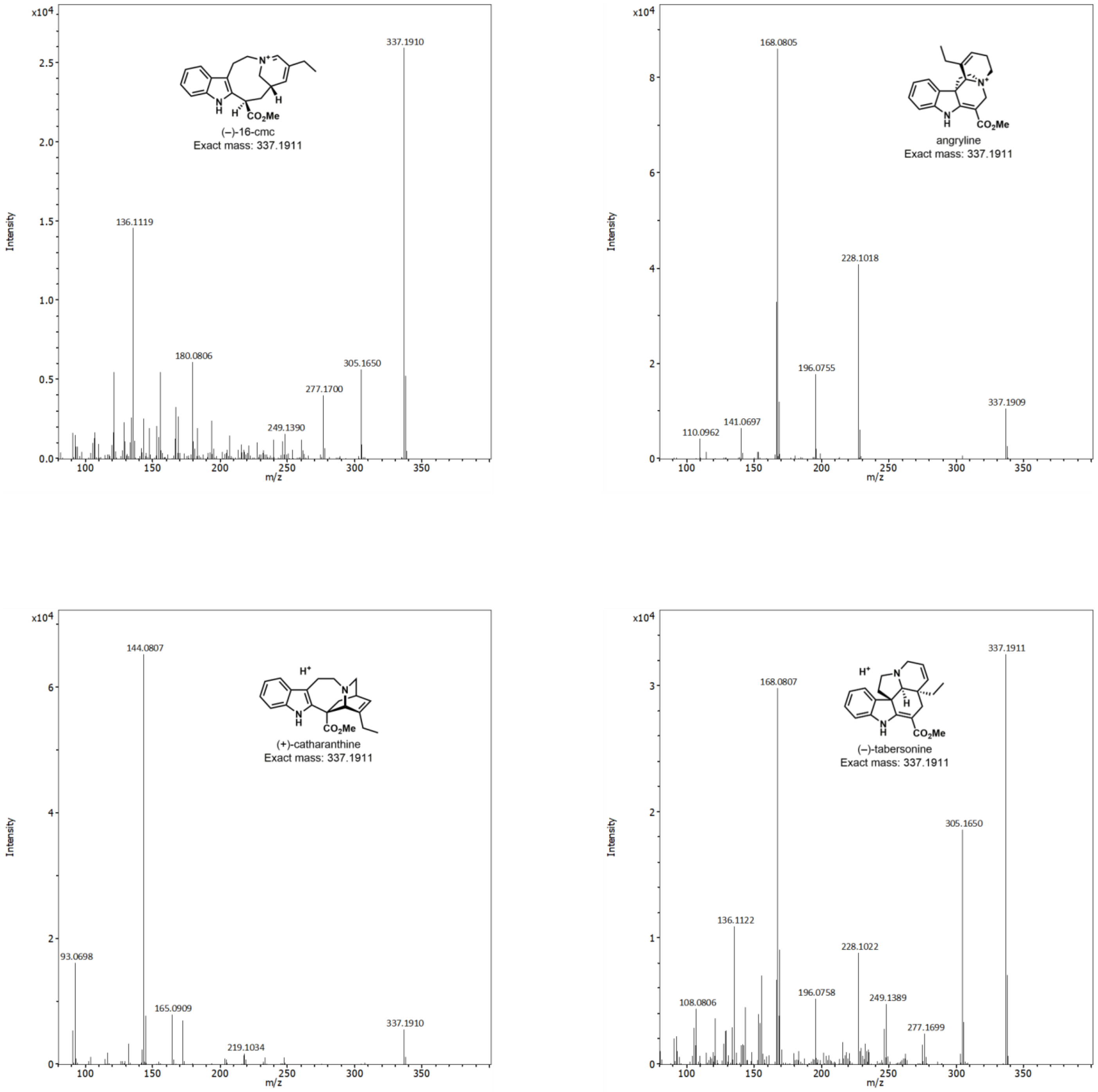


**Figure S35.** MS2 spectra of standards (–)-16-cmc, angryline, (+)-catharanthine, and (–)-tabersonine.
